# MesoSCOUT: A novel tool for revealing mesoscale organization in the white-matter connectome

**DOI:** 10.1101/2025.06.19.660631

**Authors:** Emil Dmitruk, Christoph Metzner, Volker Steuber, Shabnam N. Kadir

**Affiliations:** Biocomputation Research Group, Department of Computer Science, University of Hertfordshire, Hatfield, United Kingdom; Institute of Software Engineering and Theoretical Computer Science, Technische Universität Berlin, Germany; Department of Child and Adolescent Psychiatry, Charité – Universitätsmedizin Berlin, Berlin, Germany

## Abstract

We present MesoSCOUT (Mesoscale Structural Connectome Order-complex Unveiling Topology), a framework for characterizing white matter connectivity using a method developed from computational algebraic topology: persistent homology (PH) via clique topology. Applying this method to schizophrenia, we uncover a novel, mesoscale perspective of the differences in the white-matter connectome between healthy controls (HC) and subjects with schizophrenia (SCH). We extract and compare topological motifs found in the structural connectomes of the subjects in the two groups and find significant differences. We explore the overlap of mesoscale structures found in two different datasets, *COBRE* (Center of Biomedical Research Excellence) and *HCP* (Human Connectome Project). Differences in acquisition usually render experiments recorded on different scanners incomparable, but MesoSCOUT enables cross-dataset comparisons. Our method offers a way to establish connectomic fingerprinting that could lead to a neuroimaging-based diagnosis of schizophrenia and other psychiatric and neurological conditions as well as the development of new treatments.

## 1 Introduction

The white matter connectome, the network of structural connections between brain regions, plays a central role in shaping cognition, behavior, and vulnerability to neurological and psychiatric disorders. Advances in diffusion MRI and tractography have enabled increasingly detailed reconstructions of these networks, offering new opportunities to study brain organization at multiple scales. However, most analytical approaches focus on local or global properties, often overlooking the mesoscale architecture, namely, subnetworks and motifs that may be critical for integrative brain function and disease mechanisms.

Studies of the connectome, particularly those using graph theory, tend to be node-centric or edge-centric, (e.g. graph theory measures of the degree distribution, clustering coefficient, characteristic path length, [Stam and Reijneveld, 2007], rich club coefficient [van den Heuvel et al., 2013, Van Den Heuvel and Sporns, 2019], small-worldness [Rubinov and Sporns, 2010, Sporns, 2013], edge functional connectivity [Faskowitz et al., 2020, Faskowitz et al., 2022], centrality and controllability [Dimulescu et al., 2021]). However, these metrics are fundamentally limited to pairwise interactions and overlook higher-order relationships among brain regions.

Topological data analysis (TDA) [Zomorodian and Carlsson, 2005], a method rooted in computational algebraic topology, considers information not only from nodes and edges but also structures of all possible higher dimensions. It, therefore, allows for the characterisation of higher-order relations and the identification and quantification of more complex interrelationships between multiple brain areas.

We hypothesize that many brain disorders involve non-uniform degeneration of white matter, leading to altered relative prominence of specific connectivity patterns within the structural connectome. That is, certain white matter structures may become disproportionately dominant or diminished relative to the rest of the brain’s network. Existing methods do not capture this nuance, as they typically focus on absolute measures of connectivity or regional morphometrics, without considering how degeneration reshapes the balance of mesoscale structures across the connectome.

To address this gap, we introduce a TDA framework based on persistent homology via clique topology, MesoSCOUT, which enables the detection of mesoscale motifs that can reflect higher-order relationships among brain regions. Crucially, our method is designed to capture the relative strength of these motifs within each individual’s brain, rather than relying on absolute fibre counts. This individualized, rank-based approach allows us to quantify how prominent various structures are in a given connectome, and how their prominence differs between subjects with a disorder and those without. We demonstrate the utility of our approach on schizophrenia.

While several studies employing TDA have aimed to build a general understanding of the topology of functional connectivity [Singh et al., 2008, Ellis and Klein, 2014, Lord et al., 2016, Phinyomark et al., 2017, Anderson et al., 2018, Ellis et al., 2019, Caputi et al., 2021], others have specifically focussed on how functional brain connectivity is altered in brain degenerative diseases and psychiatric disorders, such as ADHD [Lee et al., 2011, Ellis and Klein, 2014], Parkinson’s disease [Zhang et al., 2023], schizophrenia [Stolz et al., 2017, Stolz et al., 2018, Caputi et al., 2021, Alexander-Bloch et al., 2013, Hadley et al., 2016] or early psychosis [Fournier et al., 2020]. So far, structural connectivity has only been studied in healthy subjects [Sizemore et al., 2018], leaving its role in brain disorders unexplored.

Schizophrenia is a severe and chronic mental disorder that significantly affects how an individual thinks, feels, and behaves. It has a wide range of symptoms, ranging from psychosis, hallucinations and delusions, to disturbances in cognition. Despite decades of research, the causes are still unclear [Howes et al., 2023, Shen et al., 2023].

White matter deterioration is recognised as a key contributor to the pathophysiology of schizophrenia [Konrad and Winterer, 2007,Ellison-Wright and Bullmore, 2009]. While recent reviews have reported alterations across entire fibre tracts [Vitolo et al., 2017, Male et al., 2024], these findings often lack specificity regarding which tract components are affected—an important limitation given that individual tracts can span numerous brain regions.

Furthermore, studies of schizophrenia typically focus either on global brain measures (e.g. overall cortical folding [Sasabayashi et al., 2021, Rosa et al., 2021, Kuo and Pogue-Geile, 2019]) or on local changes in specific brain areas (e.g. cortical thickness [van Erp et al., 2018]). Notably, evidence suggests schizophrenia is associated with degeneration in the white matter connectome [Fornito et al., 2011, Zalesky et al., 2011, Repple et al., 2023].

MesoSCOUT which explores the structural white matter connectome using methods of TDA on neuroimaging data, uncovers both specific differences in mesoscale-level brain structure, as well as similarities between the SCH and HC groups. We calculate structural networks of brain regions from diffusion-weighted tensor imaging (DWI) and T1-weighted MRI images by computing persistent homology obtained from the order complex derived from the white matter connectivity matrix. Through this process, we measure topological features reflecting the organisation of cliques in a person’s connectome and obtain the mesoscale-level brain structures, described by persistent homology classes of cycles. These topological structures are grouped by dimension depending on whether they correspond to clusters, holes, hollow voids, etc.

Our method of analysis provides a key insight: the relative prominence of different mesoscale structures within a connectome is important and may carry functional significance. For instance, while a set of brain regions may exhibit similar connectivity patterns and appear nearly identical on neuroimaging scans in both SCH and HC, their relative strength compared to other connections in the same brain can differ markedly. In one individual, these connections may dominate the structural connectome, while in another, they may be drowned out by stronger connections between different regions. These relative differences can be detected in the ‘persistence landscapes’, and we find significant differences between SCH and HC. Therefore, sub-networks of the brain that appear nearly identical in the two groups under some neuroimaging protocol may, nevertheless, in fact, play more dominant roles in one group than the other, thereby causing differences in behaviour and symptoms.

Importantly, we demonstrate that MesoSCOUT is robust to differences in scanning equipment and acquisition protocols - the majority of the most prominent mesoscale structures were shared between HC from the *HCP* dataset [Van Essen et al., 2012] and the *COBRE* dataset, enabling comparison across datasets.

Our approach is a first step towards the development of neuroimaging biomarkers for psychiatric and neurological conditions such as schizophrenia.

## 2 Results

### 2.1 Overview of analytical pipeline

MesoSCOUT applies TDA to investigate mesoscale structural organization in the human brain. Here, we demonstrate MesoSCOUT on schizophrenia datasets, focusing on differences between SCH and HC groups. We use DWI and T1-weighted MRI data from the publicly available COBRE dataset, comprising 44 SCH and 44 HC subjects (see Section 4.1.1 for details).

Structural connectivity matrices were derived by mapping white matter tracts between regions defined by the AAL2 atlas (see Section 4.2.1 and Figure 7). These matrices served as the basis for constructing order complexes, from which persistent homology was computed for various dimensions. Persistent homology captures the birth and death of topological features—such as connected components, loops, and voids—across a filtration of the connectivity matrix, enabling quantification of higher-order relationships beyond pairwise connections. A detailed description of the method is provided in Section 4.3, and a toy example illustrating the filtration process is shown in Figure 8 to aid intuition. To give some intuition about the relation between the connectivity structure and topological properties (cycles, birth times, and persistence), we present three toy examples in Figure 1.

**Figure 1.**
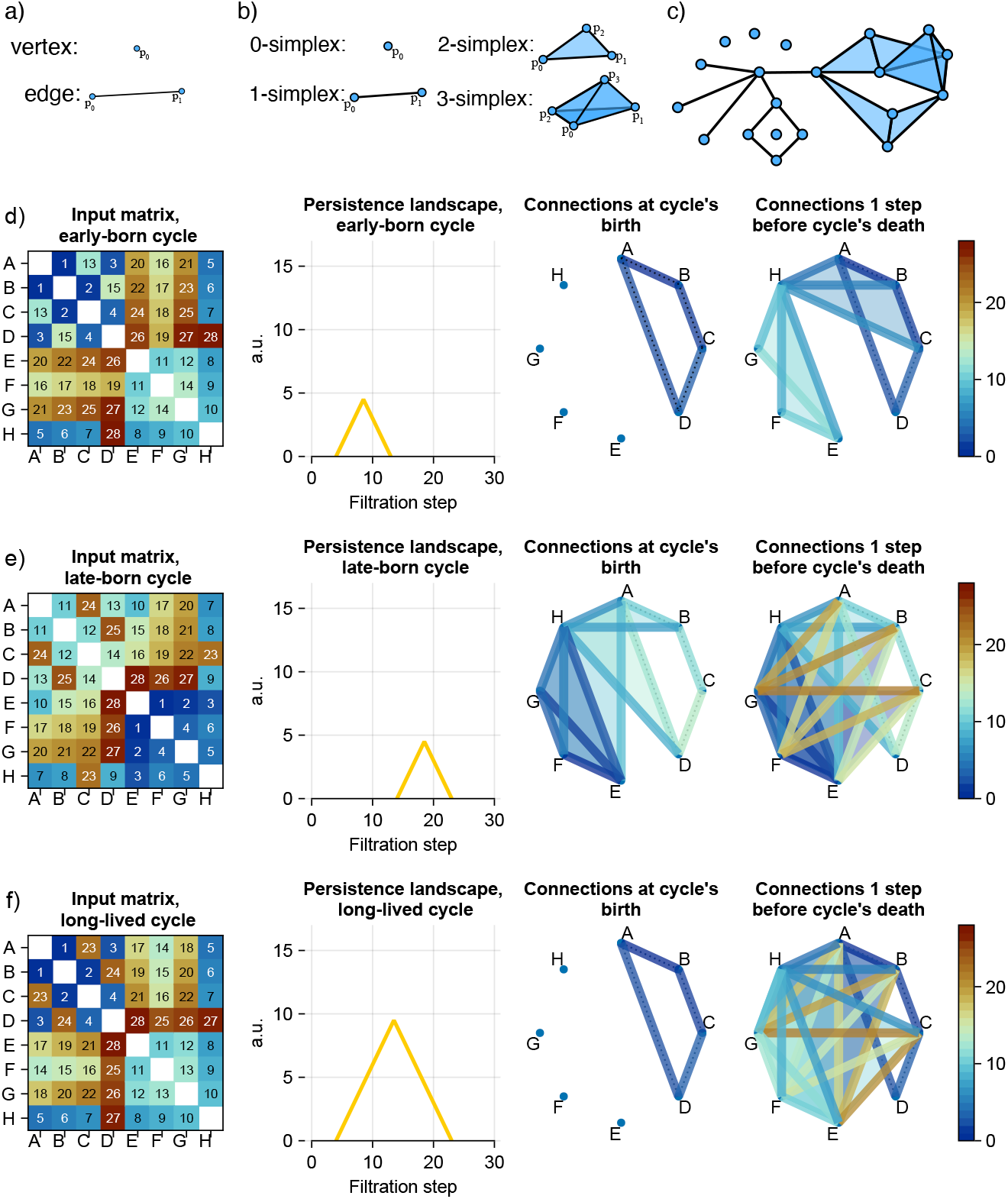
a) Building blocks of a graph (vertices and edges). b) Building blocks of a simplicial complex (a 0-simplex is a node, a 1-simplex is an edge. higher-dimensional structures convey multi-connectivity; so a 2-simplex is formed from 3 connected nodes forming a triangle; a 3-simplex is formed from 4 all-to-all connected nodes (or clique), etc. In general, an *n*-simplex corresponds to a clique with (*n* + 1) nodes. Every subclique of a clique is a lower-dimensional clique. c) A simplicial complex. d)-f) Toy examples illustrating differences in persistent sublandscapes given different birth time and persistence of a cycle that is present in all three cases. Each row presents an example of a topological structure, with d) an early-born and short-lived cycle; e) a late-born and short-lived cycles; f) early-born and long-lived cycle. For d)-e), figures in columns are (from left to right): input ordered matrix, related persistent landscape in dimension 1, visualisation of the nodes’ connectivity when a cycle is born and 1 step before it dies (the colour bar corresponds to numbers in the ordered matrix). Here, all three 8 × 8 connectivity matrices possess a 1-dimensional cycle (ABCD) with the same structure connecting the same four nodes. In (d) and (f), this cycle is born early when very few (only 4) connections have been added to the clique complex. In (e), it is born later, when there are already 15 connections in the clique complex before the formation of the cycle. The cycle, however, is more persistent in (f) because it is only ‘killed’ at step 25 of the filtration and therefore has a lifetime of 15 steps. The same cycle derived from matrices in (d) and (e) has, in both cases, a lifetime of only 10 filtration steps.

If these matrices represented connectivity structures in the brain from three different people, it would show that although each person possessed exactly the same connectivity pattern (or ‘cycle’), in other words, a physically indistinguishable set of white matter tracts between the specific nodes comprising the cycle, the strength of this pattern relative to the rest of the person’s connectome was, nevertheless, very different. The extent of this difference can be seen and quantified by the very different persistent sublandscapes associated to the cycle for each person.

To characterize the anatomical constituents of each persistent cycle, we extracted binary node participation vectors indicating which brain regions were involved at the cycle’s birth. These vectors were clustered using agglomerative hierarchical clustering, separately for each dimension (see Supplementary Section A.3 for details). Full results, including the resulting dendrograms and cycle structures are visualized in Figure 2 and Supplementary Figure Supp.B.3.

**Figure 2.**
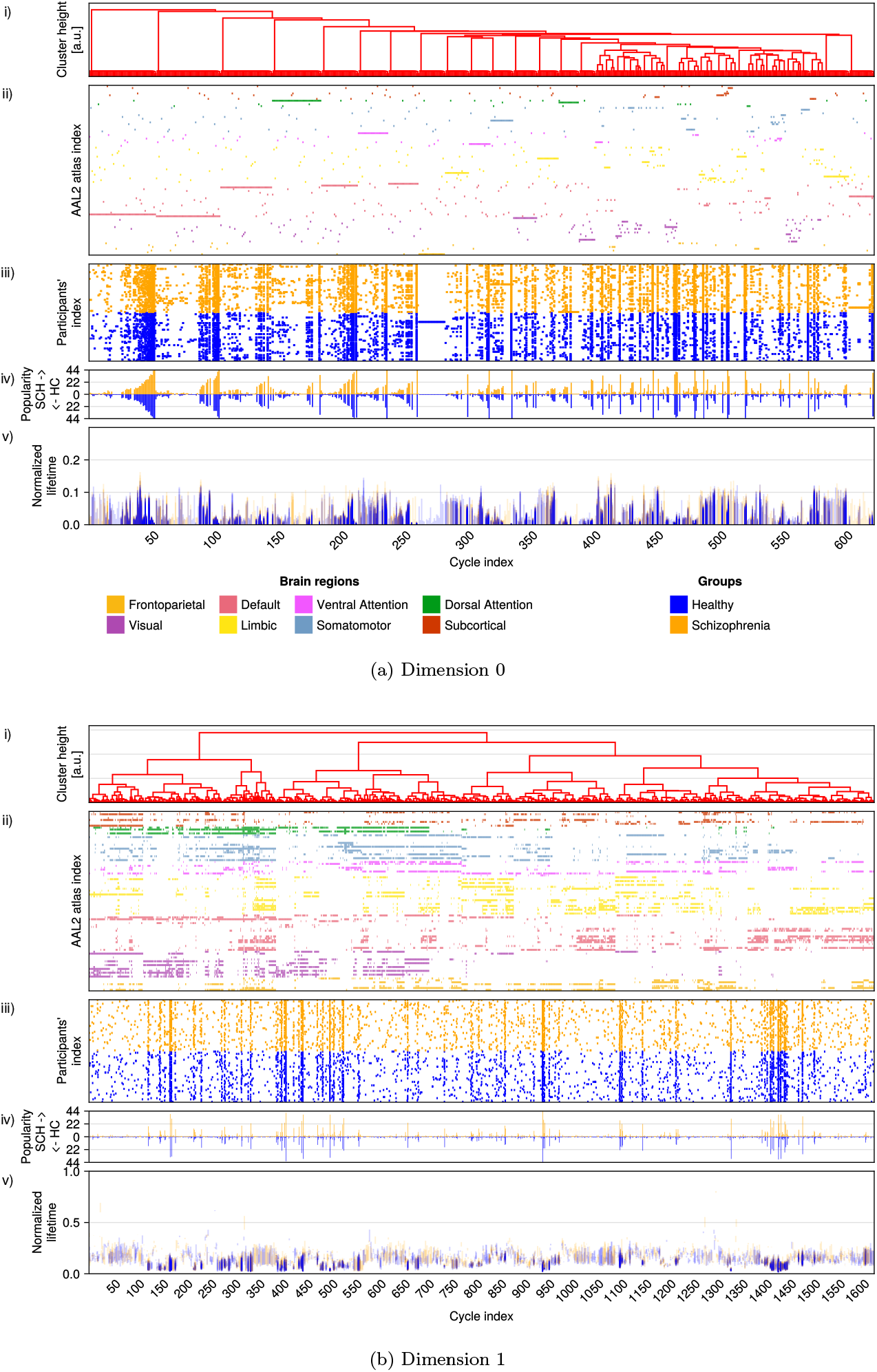
Exploration of structure, popularity, and topological properties of unique cycles in (a) dimension 0 and (b) dimension 1. Each column corresponds to a distinct persistent homology class representative cycle identified from the *COBRE* dataset (horizontal axis, shared for plots *i-iv*). (i) Dendrogram resulting from agglomerative clustering of AAL2 atlas nodes (brain regions) involved in each cycle. Clustering was performed separately for each dimension using Hamming distances on the binary node participation vectors shown in (ii), more details are presented in Supplementary Section A.3. (ii) Binary node participation vectors indicate which AAL2 brain nodes are involved in each cycle at the moment of its birth. Brain regions are colour-coded according to Yeo’s functional network atlas [Yeo et al., 2011] and ordered based on the hierarchical clustering dendrogram in (i). (iii) A subject-level raster plot showing which individuals (orange: SCH, blue: HC possess each cycle. (iv) Bar plots indicating cycle popularity, defined as the number of subjects in which each cycle appears (separated by group). (v) The (vertical) persistence barcodes for each cycle (one from each subject possessing the cycle) showing b6irth and death times normalised to the maximum filtration step. Transparency reflects the number of subjects sharing a cycle at similar filtration levels—darker regions indicate greater overlap.

An overview of the data preprocessing pipeline and topological analysis is provided in Supplementary Figure Supp.A.1.

We found that many persistent homology class representative cycles are present in multiple subjects in both the SCH and HC groups. A subset of these shared cycles exhibited significant differences in their persistence properties between groups.

### 2.2 Cycles derived from the *COBRE* data

We computed persistent homology across dimensions 0 through 3 for each subject’s structural connectivity matrix.

Intriguingly, the agglomerative clustering automatically gave rise to clusterings of persistent bar codes (shown in Figures 2a and 2b parts (v)), indicating that anatomically similar cycle classes often possess bar codes with similar birth and death times. The emergence of this correspondence is particularly striking because the hierarchical clustering only uses the vectors of brain areas and Hamming distances and is blind to the topological analysis (persistence information).

In dimension *D* = 0 shown in Figure 2a, cycles correspond to connected components present in the connectome that arise at various scales and merge with each other over the course of the filtration. For every subject, the filtration starts with 94 detached nodes (one for each AAL2 brain area). Most non-trivial zero-dimensional cycles are born near the beginning of the filtration and gradually merge, with all but one component surviving the first 16% of the way through the filtration. Representatives of the non-trivial connected components start out with 2 nodes on average. For higher dimensions, *D* = {1, 2, 3 }, cycles share multiple brain regions as shown in Figure 2b(*ii*) and Supplementary Figure Supp.B.3(*ii*).

There are cycles in dimensions 0 and 1 that were repeatedly found in multiple subjects. However, shared cycles were the minority compared to the unique cycles that were found in only single subjects (“singletons”). In dimension 1, out of 1625 distinct cycles, 1130 cycles (69.53%) were singletons. A similar observation can be made for cycles in dimension 0.

In dimensions *D* = 2 and *D* = 3, cycles were even more individualised, with the majority being singletons. Despite their uniqueness, many of these high-dimensional cycles shared structural similarities with popular dimension 1 cycles (see Figure Supp.B.3(*ii*)), often forming when lower-dimensional cycles were killed. These cycles are also constructed out of more nodes than lower-dimensional cycles (mean total nodes per cycle, for dimension *D*: 1.97 ± 0.17 (*D* = 0), 10.44 ± 5.24 (*D* = 1), 15.61 ± 8.58 (*D* = 2), 12.93 ± 3.76 (*D* = 3)). We can cluster these high-dimensional cycles into larger clusters comprising of cycles with subtle inter-individual variations (see Supplementary Figure Supp.B.4). The existence of these ‘variant’ cycles mirrors findings in the fMRI study [Gordon et al., 2017] where spatially variable ‘network variants’ were observed for all individuals and which differed in small ways from network structures obtained from group-averaged data analyses, e.g. [Yeo et al., 2011].

Interestingly, many singleton cycles, despite appearing in only one subject, exhibited structural similarity to more ‘popular’ cycles (defined as those present in at least 25% of subjects. Due to the non-uniqueness of the cycle representatives in persistent homology, these could nevertheless stem from the same underlying persistent homology class. This coherence in structure, popularity, or persistence was not observed in the Random or Geometric null models, e.g. Supplementary Figure Supp.B.25 shows the lack of consistent cycle structure for connectivity matrices derived from distances between randomly chosen points in the unit cube in ℝ^5^.

### 2.3 Persistence landscapes

We examine differences in the persistence landscapes between the SCH and HC groups of the *COBRE* dataset, firstly looking at global landscapes, followed by a more fine-grained study of sublandscapes.

#### 2.3.1 Global persistence landscapes

Figure 3 shows the averaged global landscapes computed for the two studied groups: 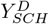 and 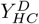, for dimensions *D* = 0, 1, 2, 3, as well as the layer-by-layer differences 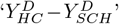 between these landscapes.

**Figure 3.**
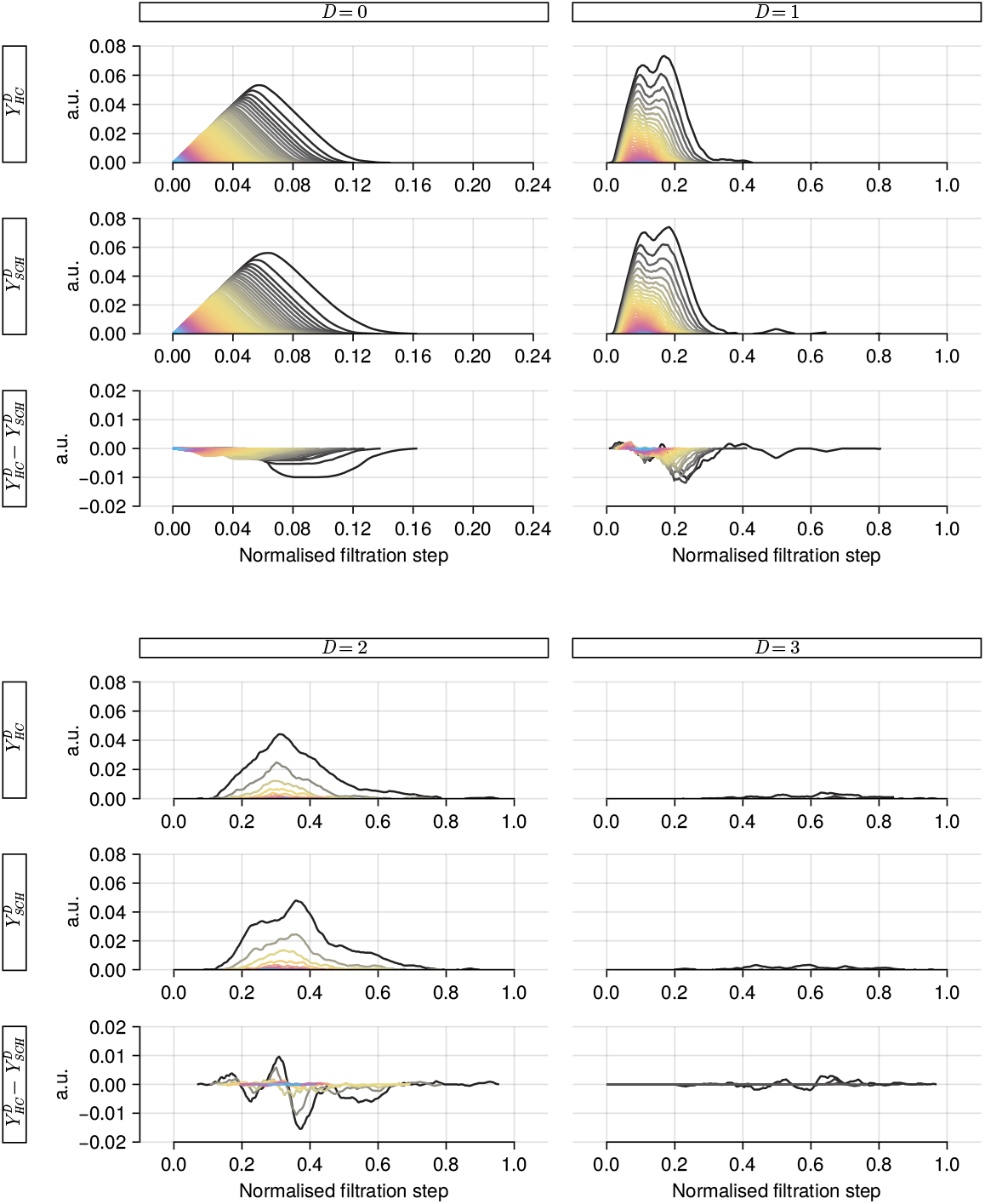
Average global persistence landscapes for healthy controls 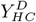, schizophrenia subjects 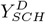 and the difference 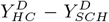 for topological features in dimensions *D* = 0, 1, 2, 3, as indicated by subtitles and labels on the left-hand side. The horizontal axis represents the filtration step normalised to the total number of steps in each filtration (the range for dimension 0 is limited to filtration value 0.24, the full range shown for other dimensions). The vertical axis is the height of the landscape expressed in arbitrary units (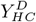 and 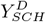 are shown in the same scale). The colours in the persistence landscapes are related to the landscape layer- the outer layers (e.g. >_0_) are denoted with black, then the colour scheme goes from yellow to pink and then to blue for deeper layers. See Figure A.2.3 for an explanation of landscape averaging.

Significant differences in the global persistence landscapes between SCH and HC cohorts were found for both dimensions 0 and 1. Specifically, the landscapes for SCH subjects denoted, 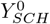 and 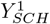, exhibited significantly greater persistence than those for HC subjects, 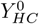and 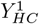, as determined by a one-sided permutation (*n* = 10000) test (dimension 0: *p* = 0.014; and dimension 1: *p* = 0.018); see Supplementary Section A.2.4 for details. These differences are visualised in Figure 3, where layer-by-layer differences 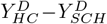 (for *D* = {0, 1 }) are predominantly negative, indicating greater persistence in the SCH group across all layers.

For dimension 1, all cycles emerged early in the filtration (mean birth time 0.088 ± 0.058), therefore, they are formed with strong connections.

No statistically significant differences were found in the average global persistence landscapes for dimensions 2 and 3 (one-sided permutation test, *p* = 0.205; and *p* = 0.509 respectively).

#### 2.3.2 Sublandscape analysis identifies candidate biomarkers

For each popular cycle, we performed permutation tests (Section A.2.4) on the associated sublandscapes to examine group differences in their persistence, controlling for multiple comparisons using the Benjamini-Hochberg (BH) procedure for False Discovery Rate (FDR). Among the significantly different cycles, three were found in at least 44 subjects (50% of the subjects) and showed significant group differences, suggesting their potential as candidate biomarkers for schizophrenia.

##### Popular cycles for dimension 1

Out of a total of 55 popular dimension 1 cycles, 22 cycles were significantly different (5 more cycles with p-values below 0.05 did not survive the FDR correction). A complete set of popular sublandscapes and sublandscape distance distributions under the permutation test can be found in Supplementary Section B.5 in Figures Supp.B.8 -Supp.B.12. The group-averaged sublandscapes corresponding to each popular cycle individually are shown in Supplementary Figure Supp.B.5.

##### Sublandscape centroids reveal distinct patterns of disruption

Centroids (Section A.3) were used to characterise qualitative differences between the sublandscapes of the two groups (see Figure 4a). Vectors were drawn from the HC centroid to the SCH centroid for each cycle and plotted in polar coordinates in Figure 4b. Agglomerative clustering of these vectors based on cosine distance (Figure 4c) confirms two distinct clusters reflecting two types of changes in brain organisation seen in schizophrenia.

**Figure 4.**
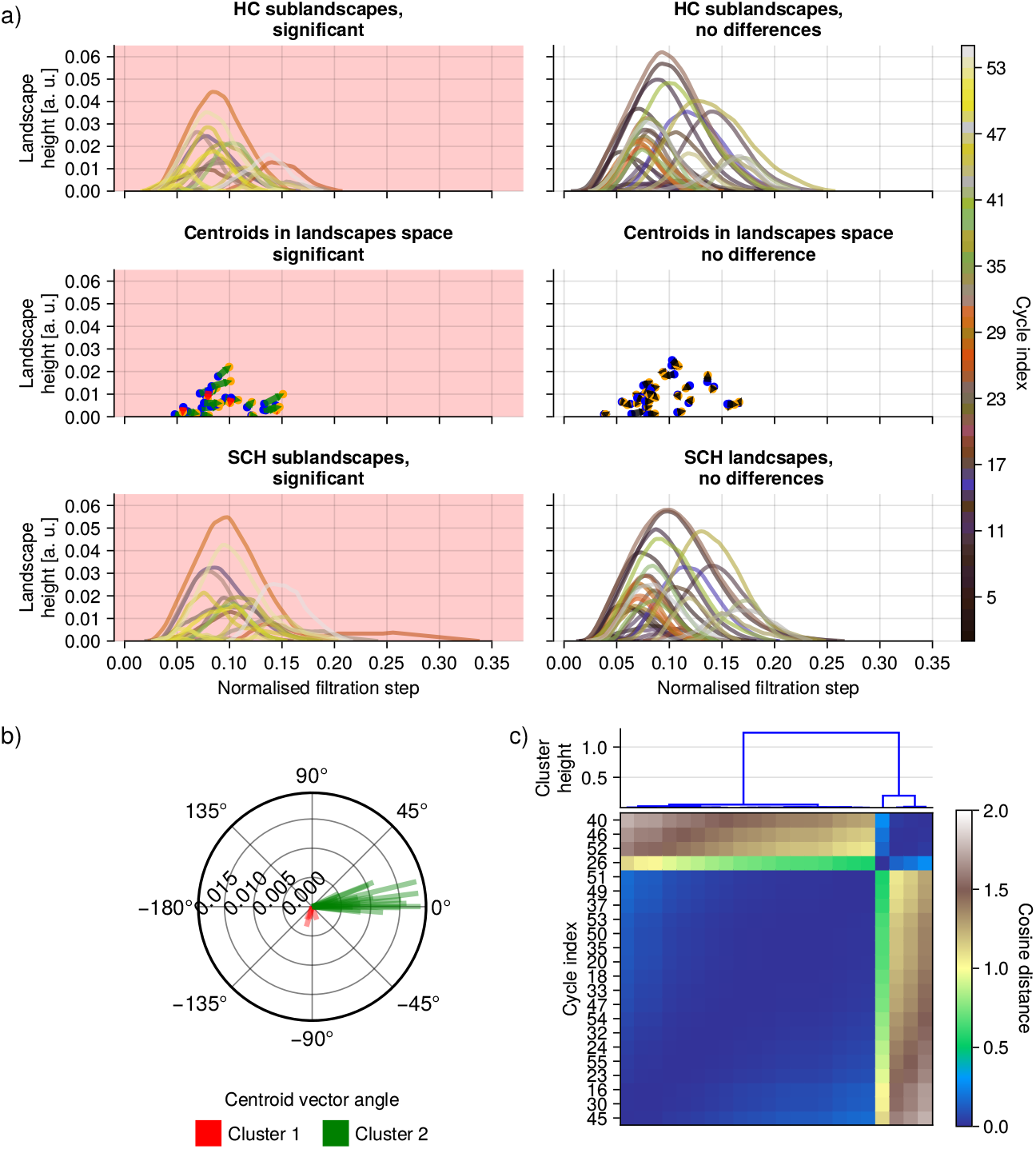
Decomposition of the global landscape in terms of popular sublandscapes (a) (data for cycles in dimension 1), the polar plot of centroid difference vectors (b) and cosine distance matrix with clustering (c). In the middle plot of (a), each arrow indicates how the centroids in dimension 1 are displaced for the 22 popular persistence sublandscapes that were found to be significantly different between the two groups (popular cycles were cycles found in at least 22 subjects). In the top and bottom rows of (a), persistence sublandscapes are presented for the HC group (top) and the SCH group (bottom), for statistically different cycles (first column) and for cycles where no significant difference was found (second column). Persistence sublandscapes are coloured according to the cycle they are derived from, i.e. the same colour in HC and SCH is used for the same cycle. The vectors shown in (b) indicate the direction of the centroids shift in the cycles in dimension 1 that are significantly different between HC and SCH group. Clustering (c) indicates two distinct groups, cluster 1 (cycles 26, 40, 52, 56) marked as red, cluster 2 with the remaining cycles. Vectors in the raster plot indicate how the centroid of SCH is placed when compared to equivalent cycle in the HC group.

The first cluster in red (cycles 26, 40, 46, 52) comprises persistence sublandscapes that are all ‘lower’ for SCH than in HC due to earlier death times for SCH cycles. The second cluster in green (18 cycles) comprises of persistence sublandscapes that are mostly shifted upwards and right for SCH compared to HC. Consult Supplementary Section B.3 for a more detailed discussion on sublandscape differences. Structures found in schizophrenia-affected patients are therefore ‘delayed’ (both are dying later, while some are also born later). The shift is consistent with the fact that inner layers of the dimension 1 global landscape, to which sublandscapes contribute, are formed later for SCH than for HC due to weaker connections (both birth and death of cycles for SCH are occurring later in the filtration).

#### 2.3.3 Dimension 1 cycle structures and brain regions

We note that many of the brain regions partaking in the significant dimension 1-cycles are frequently mentioned in existing literature about structural brain alterations induced by schizophrenia. In general, cycles differ in both their size (total number of brain regions participating in a cycle) and in the regions involved. However, some regions are repeatedly found in many of the significant cycles - 9 brain regions were found in at least 4 cycles. We summarise these in Table 1; for further discussion on the connections to existing literature, see Supplementary Results Section B.4. A full list of cycles and their constituent regions can be found in Supplementary Table B.11.

**Table 1:**
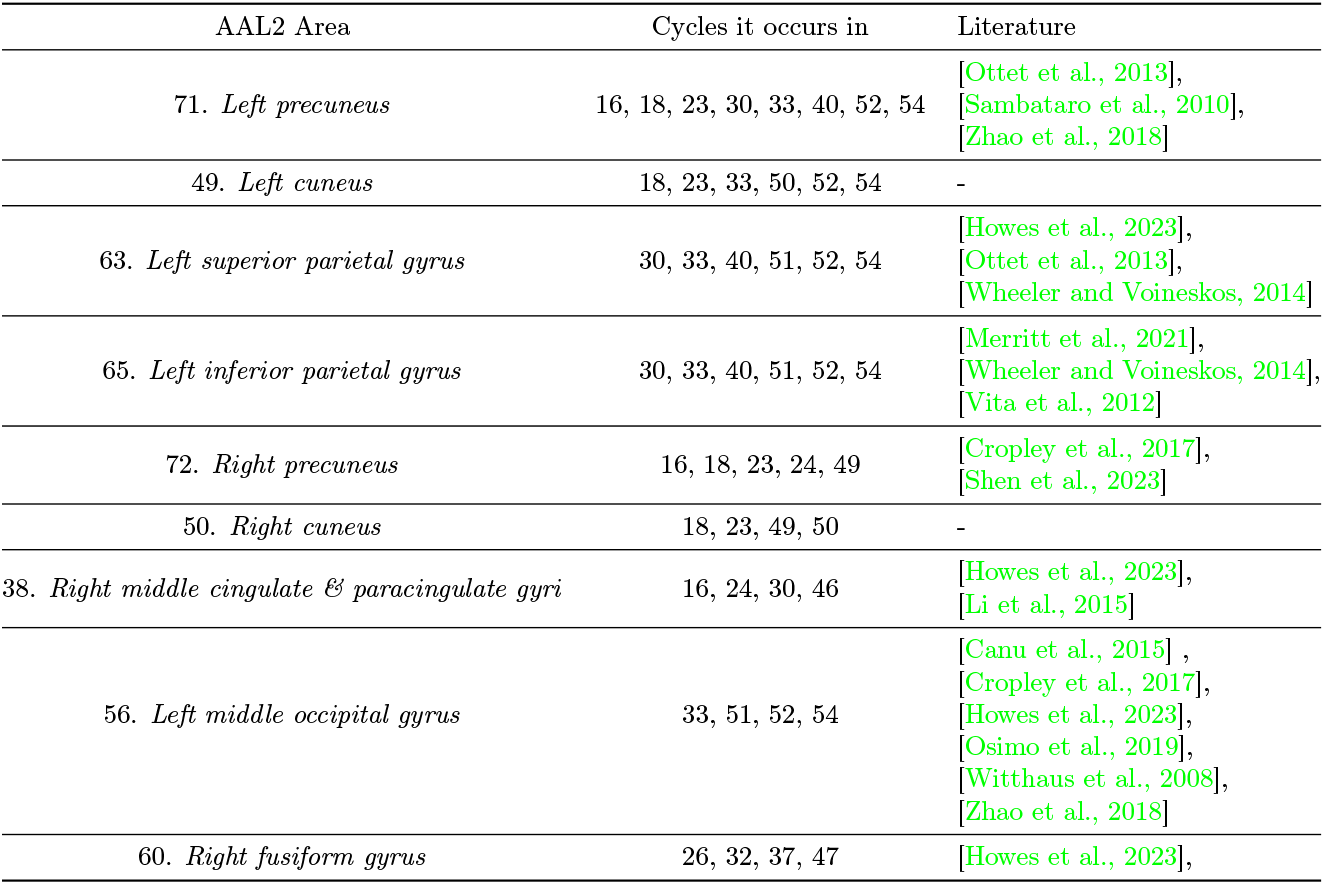
Areas that partake in significant Dimension 1 cycles and the schizophrenia literature.

| AAL2 Area | Cycles it occurs in | Literature |
| --- | --- | --- |
| 71. *Left precuneus* | 16, 18, 23, 30, 33, 40, 52, 54 | [Ottet et al., 2013],<br>[Sambataro et al., 2010],<br>[Zhao et al., 2018] |
| 49. *Left cuneus* | 18, 23, 33, 50, 52, 54 | - |
| 63. *Left superior parietal gyrus* | 30, 33, 40, 51, 52, 54 | [Howes et al., 2023],<br>[Ottet et al., 2013],<br>[Wheeler and Voineskos, 2014] |
| 65. *Left inferior parietal gyrus* | 30, 33, 40, 51, 52, 54 | [Merritt et al., 2021],<br>[Wheeler and Voineskos, 2014],<br>[Vita et al., 2012] |
| 72. *Right precuneus* | 16, 18, 23, 24, 49 | [Cropley et al., 2017],<br>[Shen et al., 2023] |
| 50. *Right cuneus* | 18, 23, 49, 50 | - |
| 38. *Right middle cingulate & paracingulate gyri* | 16, 24, 30, 46 | [Howes et al., 2023],<br>[Li et al., 2015] |
| 56. *Left middle occipital gyrus* | 33, 51, 52, 54 | [Canu et al., 2015],<br>[Cropley et al., 2017],<br>[Howes et al., 2023],<br>[Osimo et al., 2019],<br>[Witthaus et al., 2008],<br>[Zhao et al., 2018] |
| 60. *Right fusiform gyrus* | 26, 32, 37, 47 | [Howes et al., 2023], |

##### Significant cycles with low persistence

Persistence landscapes 20, 35 and 53 are constructed from small cycles involving four brain regions each, and exhibit very low persistence along with significant variability in their landscapes. Although low persistence cycles are typically dismissed as noise in TDA, their consistent appearance across multiple subjects (observed in 44, 32, and 23 subjects, respectively) suggests they may be meaningful biomarkers. Notably, at least two regions in each cycle have been previously associated with schizophrenia (see Supplementary Table 5), further supporting their potential relevance.

The importance of low-persistence cycles has been highlighted in prior applications of TDA to medical data [Kanari et al., 2018, Bendich et al., 2016], in particular, in a study of connectivity in functional magnetic resonance imaging [Stolz et al., 2017].

#### 2.3.4 Cycle Laterality

We examined the hemispheric distribution of the 55 popular dimension-1 cycles to assess structural asymmetry. Table 2 categorizes which significant cycles were left-lateralized, right-lateralized, symmetric, or could not be classified. Notably, we observed a strong asymmetry, as 16 cycles were confined to a single hemisphere, and none of these had a corresponding “mirror cycle” composed of anatomically homologous regions in the opposite hemisphere (see Supplementary Results B.2.2).

**Table 2:** Laterality of significant dimension 1 cycles.

| Laterality | Cycles exhibiting property |
| --- | --- |
| Left | 20, 33, 40, 51, 52, 53, 54, 55 |
| Right | 24, 26, 32, 35, 37, 45, 47, 49 |
| Symmetric | 16, 18, 23, 50 |
| Mixed | 30, 46 |

**Table 3:** Illness information. The *COBRE* dataset comes with the evaluation of the state of SCH patients done with a Positive and Negative Affect Schedule (PANSS) scale [Zevon and Tellegen, 1982, Watson et al., 1988].

| | Mean (± std. dev.) | Min | Max |
| --- | --- | --- | --- |
| Age | 38.30(±13.96) | 19 | 65 |
| Age illness onset | 21.33(±7.04) | 12 | 47 |
| Illness duration | 16.96(±13.14) | 1 | 41 |
| Medication- CPZ equivalent dose | 373.52(±331.86) | 25 | 1800 |
| PANSS- Positive | 14.34(±4.99) | 7 | 28 |
| PANSS- Negative | 14.45(±4.96) | 8 | 29 |
| PANSS- Gen. Psychosomatic | 28.48(±8.26) | 16 | 56 |

Across the full set of popular cycles, we identified eight “mirror twin” pairs of cycles, each composed of homologous regions in opposite hemispheres. Of these, three pairs consisted of one cycle that was significantly different between groups (cycles 26, 37, and 47—all right-lateralized) and one non-significant counterpart (cycles 42, 48, and 34, respectively). The remaining five pairs were composed of non-significant cycles. These patterns are visualized in the frequency scaffolds in Supplementary Figure Supp.B.7.

### 2.4 Null models

Given the highly non-random architecture of the human brain [Bullmore and Sporns, 2012, Sporns, 2013], we anticipated the presence of recurring structural motifs across individuals, alongside subject-specific variability. Null models (NM) (see Section 4.4 for descriptions) enabled us to probe the underlying structure of the data and assess whether the observed cycles could arise by chance or reflect meaningful biological organization. Full details of this analysis are provided in Supplementary Results Section B.8.

### 2.5 Cross-Dataset Structural Consistency

Despite differences in acquisition protocols between the *COBRE* and *HCP* datasets (see Table 4), we observed a high degree of structural overlap. All 55 popular cycles identified in *COBRE* were also present in *HCP*, indicating strong cross-dataset reproducibility. This is visually evident in Figure 5(*iii*), where vertical lines representing frequently observed cycles in HC (blue) and SCH (orange) subjects from *COBRE* also appear in *HCP* subjects (red). For full sublandscape results, see Supplementary Section B.6.

**Table 4:**
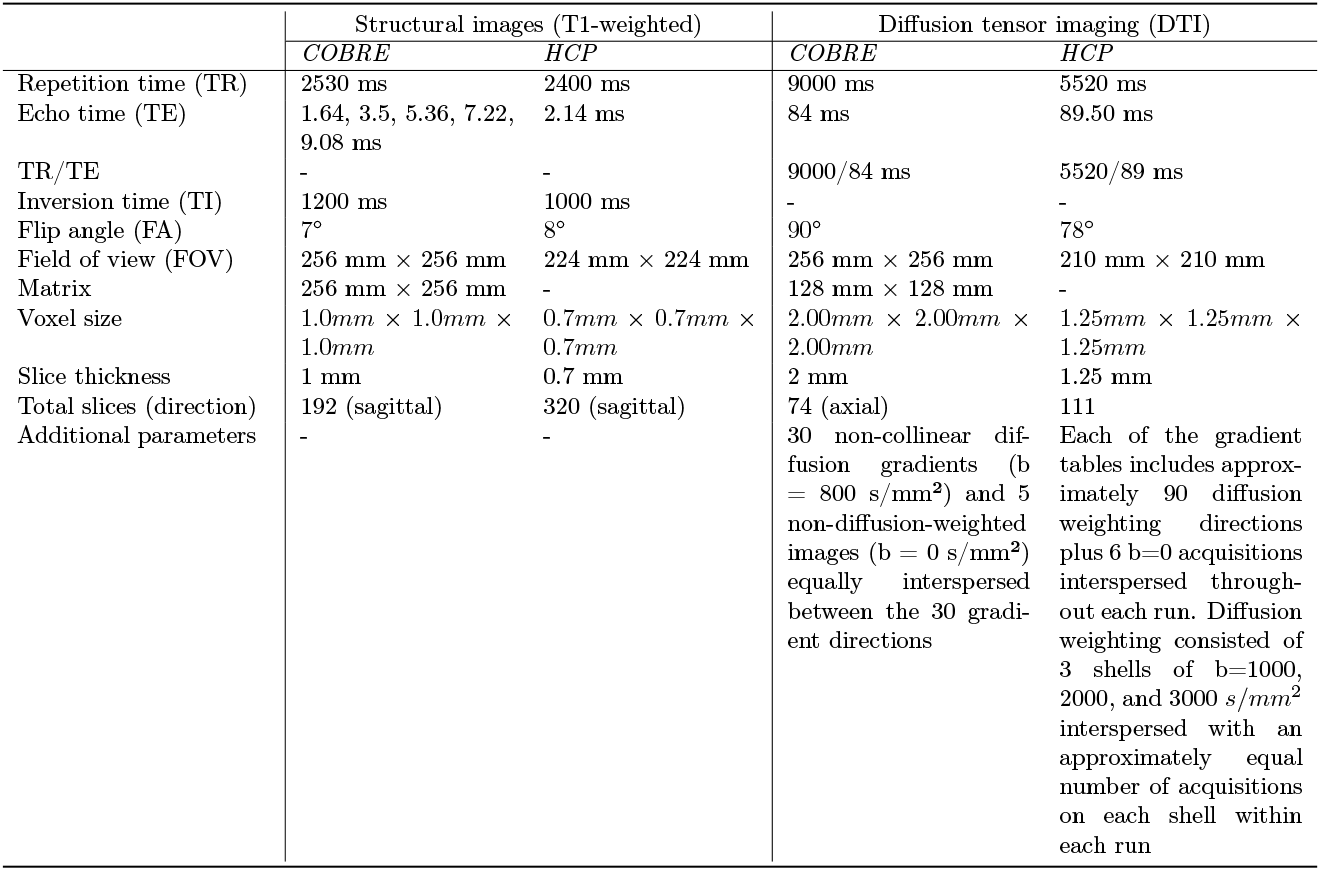
Acquisition parameters for structural and diffusion tensor imaging for *COBRE* and *HCP* datasets.

|  | Structural images (T1-weighted) |  | Diffusion tensor imaging (DTI) |  |
| --- | --- | --- | --- | --- |
|  | *COBRE* | *HCP* | *COBRE* | *HCP* |
| Repetition time (TR) | 2530 ms | 2400 ms | 9000 ms | 5520 ms |
| Echo time (TE) | 1.64, 3.5, 5.36, 7.22, 9.08 ms | 2.14 ms | 84 ms | 89.50 ms |
| TR/TE | - | - | 9000/84 ms | 5520/89 ms |
| Inversion time (TI) | 1200 ms | 1000 ms | - | - |
| Flip angle (FA) | 7° | 8° | 90° | 78° |
| Field of view (FOV) | 256 mm × 256 mm | 224 mm × 224 mm | 256 mm × 256 mm | 210 mm × 210 mm |
| Matrix | 256 mm × 256 mm | - | 128 mm × 128 mm | - |
| Voxel size | 1.0mm × 1.0mm × 1.0mm | 0.7mm × 0.7mm × 0.7mm | 2.00mm × 2.00mm × 2.00mm | 1.25mm × 1.25mm × 1.25mm |
| Slice thickness | 1 mm | 0.7 mm | 2 mm | 1.25 mm |
| Total slices (direction) | 192 (sagittal) | 320 (sagittal) | 74 (axial) | 111 |
| Additional parameters | - | - | 30 non-collinear diffusion gradients (b = 800 s/mm<sup>2</sup>) and 5 non-diffusion-weighted images (b = 0 s/mm<sup>2</sup>) equally interspersed between the 30 gradient directions | Each of the gradient tables includes approximately 90 diffusion weighting directions plus 6 b=0 acquisitions interspersed throughout each run. Diffusion weighting consisted of 3 shells of b=1000, 2000, and 3000 s/mm<sup>2</sup> interspersed with an approximately equal number of acquisitions on each shell within each run |

**Figure 5.**
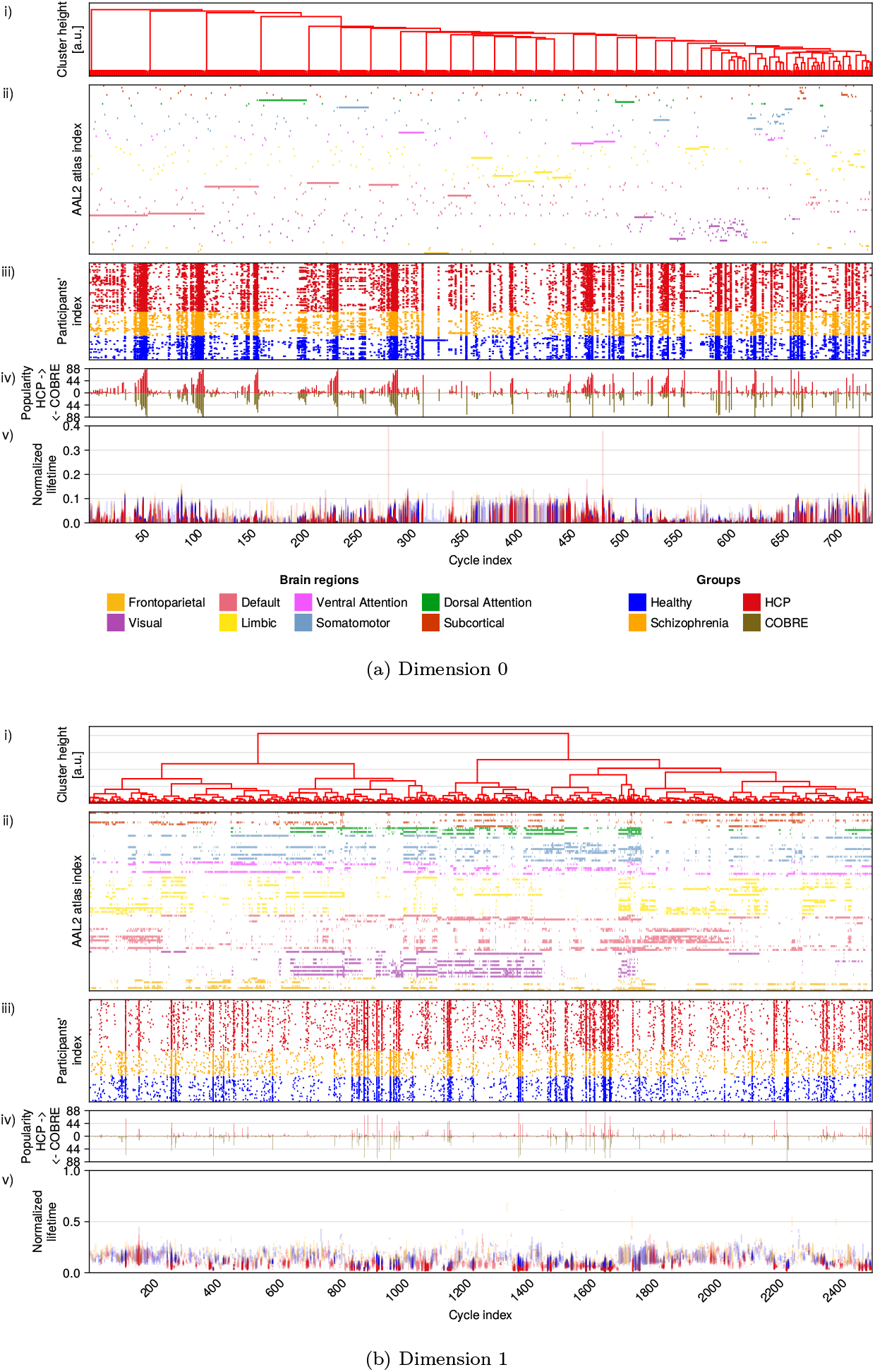
Unique cycle structures and popularity presented as a raster plot for dimensions *D* = 0, 1 for Human Connectome Project (*HCP*) and *COBRE*. For each figure, the plots are structured as described in 2. Plots (ii) are raster plots showing which brain regions participated in the unique cycles at the moment of the cycle’s birth. All unique cycles are aligned with the horizontal axis and brain regions are marked with 7 colours according to Yeo’s functional network atlas. The brain regions are ordered according to the hierarchical clustering presented at the top part of the plot. Plots (iii) are raster plots showing in which subjects a cycle was found (orange is used for SCH, blue for HC). Plot (iv) summarizes individual cycle popularity, i.e. total number of subjects, in which a cycle was found. (v) Vertical persistence barcodes for each cycle. The vertical axis shows the lifetime normalised to the maximal filtration step in the filtration. All horizontal axes are the same, using cycle indexing from hierarchical clustering.

In dimension *D* = 1, 64 cycles showed significantly different popularity between *COBRE* and *HCP*. Of these, 13 cycles (16, 20, 23, 24, 26, 30, 37, 45, 47, 49, 51, 54, 55) also exhibited significant differences in persistence sublandscapes between SCH and HC groups. An additional 9 cycles (18, 32, 33, 35, 40, 43, 50, 52, 53) were significantly different between SCH and HC in sublandscape structure but did not differ in popularity across datasets. For full results of popularity significance tests, see Supplementary Results Section B.7.

Popular cycles also showed low variability in birth time compared to non-popular cycles (Mann-Whitney U test on the distribution of standard deviation of birth times of cycles found in more than 22 subjects vs those found in only 4-22 subjects, *U*_*COBRE*_ = 5666.0, *p*_*COBRE*_ = 0.009, *U*_*HCP*_ = 4180.0, *p*_*HCP*_ = 0.0005). This suggests that popular cycles represent stable features of brain architecture.

The presence of common cycles across datasets acquired using different scanners and protocols supports the generalizability of our findings and enables meaningful comparisons of connectomes across studies.

#### 2.5.1 Reverse filtration

The weakest connections in a connectivity matrix are usually attributed to measurement noise rather than true anatomical connections [Fornito et al., 2016]. Consequently, we hypothesized that the number and structure of cycles emerging from a reverse filtration, starting with the weakest connections, would resemble those arising from random NMs.

Figure 6, shows Betti curves (see equation A.3 for a definition) from the data matrices and NMs for both forward and reverse filtrations in dimensions *D* = [0, 4],*D* ∈ ℤ. Due to the sparsity of the connectivity matrices for the *COBRE* data (mean of 2238.68 ± 501.78 zero entries), in the reverse filtration of *COBRE, β*_0_ drops to a very low value on the very first step of the filtration as the zero entries immediately induce the construction of a large connected component consisting of many nodes, unlike the case for the other models where there is a more gradual drop off from 94 detached starting nodes. On the other hand, the (reverse) *COBRE β*_1_ curve starts with a few cycle representatives, whereas all others start with no dimension 1 cycle representatives.

**Figure 6.**
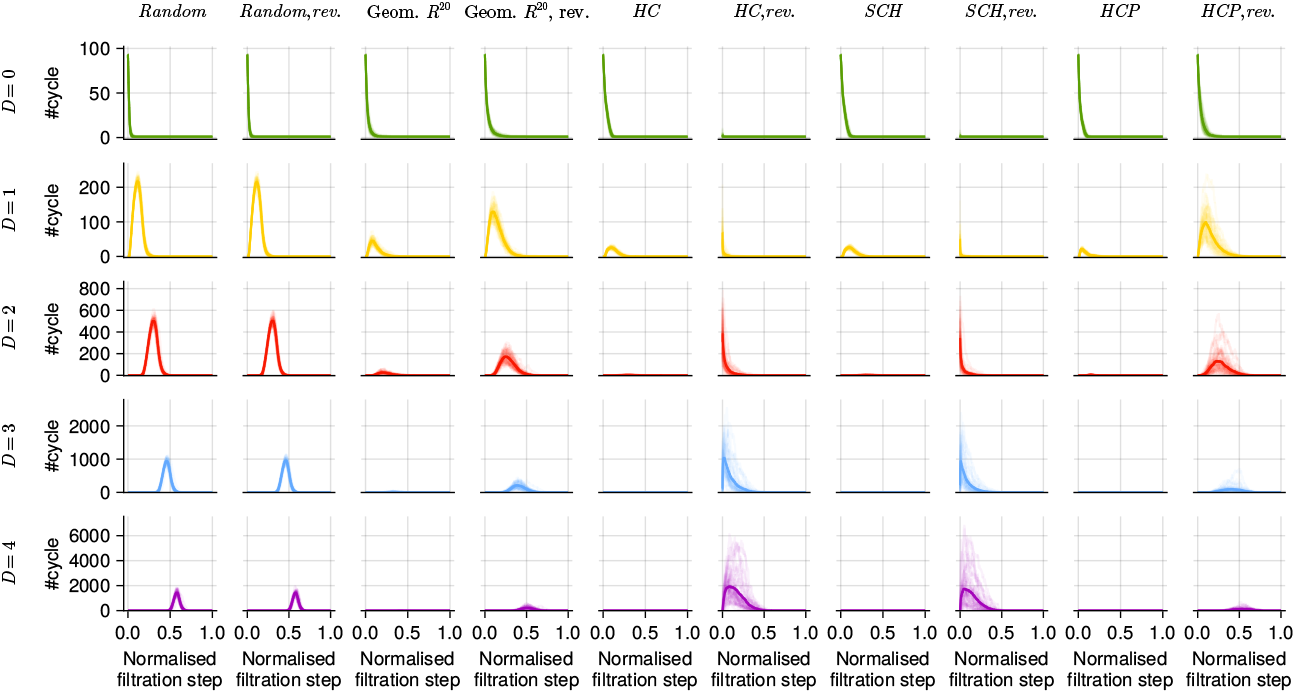
Comparison of Betti curves for Random NM, Geometric *R*^20^ NM, *COBRE, HCP* and their reverse filtration for dimension in range 0 ≤*D* ≤ 4,*D* ∈ ℤfor matrix size 94 ×94. Solid curve shows the average Betti curve computed from 44 samples; each individual sample is shown with a faint line.

**Figure 7.**
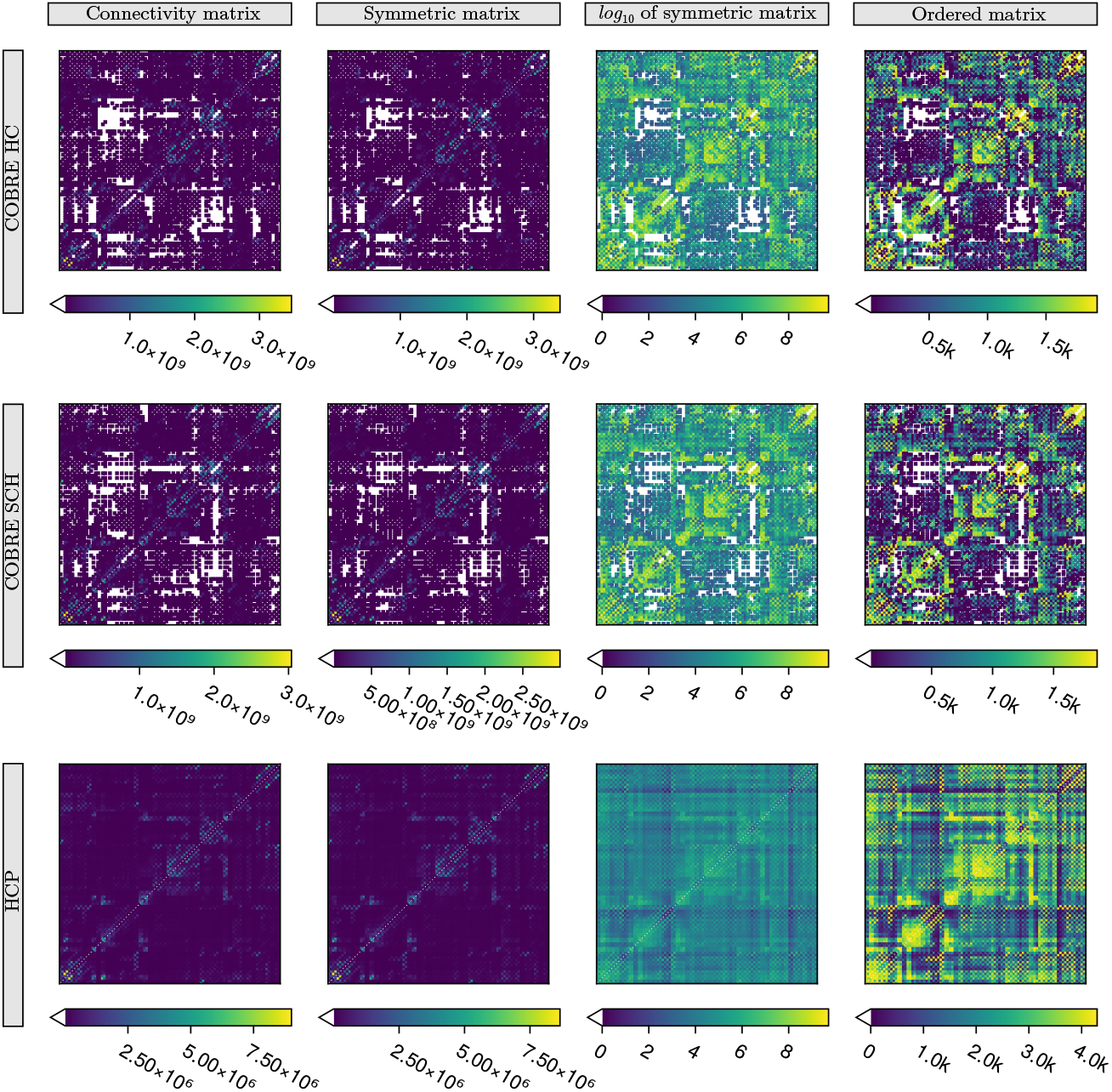
Heatmaps of sample matrices from the *COBRE* dataset (*HC* in the top row, *SCH* in the middle row) and *HCP* dataset (in the bottom row). The first column presents connectivity matrices obtained from probabilistic fibre tracking; the second corresponds to the symmetrised matrix; the third demonstrates the result of non-linear-transformation of the symmetrised matrix (*log*_10_); the fourth corresponds to the ordered matrix. In each plot (except non-linear transformation), colours are fitted on a different range of values.

**Figure 8.**
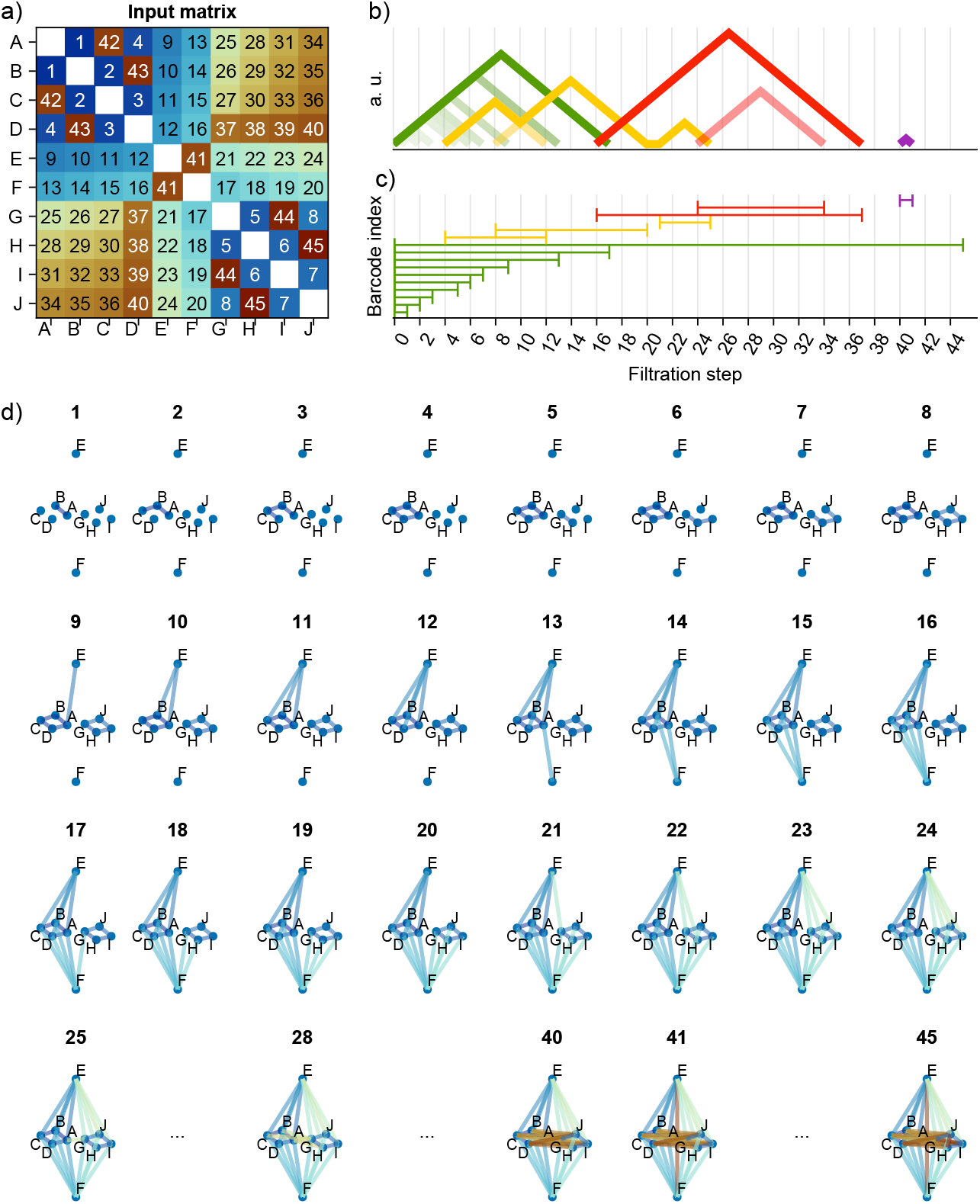
A toy example of weighted-rank clique filtration derived from an ordered matrix. a) An example of the ordered matrix; colours correspond to the filtration step; (b) persistence landscapes derived from the filtration; (c) barcodes derived from the filtration (the results for dimensions *d* = 0, 1, 2, 3, 4 are colour-coded in (b) and (c) as green, yellow, red, blue, purple, respectively); (d) simplicial complexes created at the filtration step are annotated above the figures; some steps are omitted, i.e. after all cycles died in dimension 1 (step 25) and before a fully connected graph is formed (step 45), with a few exception, e.g. when a cycle in dimension 4 is formed and dies (steps 40 and 41 respectively). The horizontal axis for (e) and (f) are filtration steps, and there are as many filtration steps, as there are unique numbers in the ordered matrix.

The distribution of Betti curves for the reverse filtration for *COBRE* resembles that of the Random NM in that they exhibit unimodal Betti curves with peak values that increase with dimension, *D*. This contrasts with the Geometric NM Betti curves, where peaks decrease with increasing *D* for *D* ≥ 1, as previously reported in [Kahle, 2013, Giusti et al., 2015b]. Interestingly, reverse *COBRE* has more cycles in dimensions *D* = 3, 4 than the Random NM, some of which stem from the large number of zeros present in the original matrix, which leads to large structures early in the reverse filtration.

To our knowledge, matrices where the Betti curves are higher than those of random matrices have not been systematically studied, suggesting that white matter connectivity may possess previously uncharacterized mathematical properties.

To explore higher-dimensional structures, we computed persistent homology for submatrices of size 30 × 30 from both datasets, obtained by taking only the first 30 AAL2 brain regions. This size limitation allowed for computational tractability up to dimension *D* = 8. As shown in Supplementary Figure Supp.B.22, the distributions of Betti curves for both datasets in this reduced form align more closely with the Random NM.

There are always more popular cycles for the data (details are presented in Supplementary Table 9) as compared to null models, especially in the forward filtration. This is particularly visually evident in Supplementary Figures Supp.B.25, Supp.B.27 for the randomised models.

##### Robustness to the presence of weak connections

To assess the impact of weak connections on topological structure, we performed a controlled shuffling of the weakest *k* ∈ {10, 20, 30, 40, 50, 60, 80, 90} per cent of edges in a representative *HCP* matrix (see Supplementary Section B.9.1). As predicted, increasing the proportion of shuffled edges caused the distribution of Betti curves to converge to that of the Random NM.

In the forward filtration, Betti curve shapes remained stable until *k* = 30% of the weakest connections were randomized. Even at this threshold, the curve shifted slightly toward earlier filtration steps but retained its overall form, indicating robustness to noise from the presence of weak connections. Notably, this 30% threshold corresponds to the average proportion of zero-weight connections in the *COBRE* dataset, demonstrating that despite stark differences in sparsity, the same topological features are recovered in both *COBRE* and *HCP*.

Further increasing the proportion of shuffled connections results in the emergence of higher-dimensional cycles. The latter are absent from the original matrix, but are expected to be present in random matrices of the same size. The shape of the Betti curves from *HCP* remains largely unchanged until the *k* = 60% level, where it starts to be overrun by a ‘tsunami of randomness’ (Figure Supp.B.23), after which there is a significant rise in the number of cycles in the randomised part of the filtration, and convergence to the Random NM profile. This transition highlights the boundary at which biologically meaningful structure gives way to noise-induced topology.

## 3 Discussion

We introduced a general and robust framework, MesoSCOUT, for characterizing white matter connectivity in disease using TDA. Instead of focussing on how a disease affects individual regions, captured as deviations in brain morphometrics, it examines connectivity patterns at multiple scales specified by the filtration steps of the order complex. While we illustrated its utility in the context of schizophrenia, the method is broadly applicable to any condition involving alterations in white matter connectivity, including other neuropsychiatric, neurodevelopmental, and neurodegenerative disorders.

By using an ordered matrix instead of directly using the output matrix produced by the probabilistic fibre tracking algorithm (Figure 7), we ensured that the derived topological properties rely on the relative connections within a participant’s brain and not on estimates of absolute fibre count. Using absolute fibre counts may have contributed to the variability of reports of schizophrenia-related brain alterations in previous studies([Wheeler and Voineskos, 2014,Howes et al., 2023]). In contrast, using ordered connections, as MesoSCOUT does, resolves this issue and makes the comparison between different groups more reliable.

Investigating the ranking of connections within individuals allows us to examine the sequence in which connections are integrated into the subject’s connectome. This individualized framework facilitates a subject-specific characterization of brain connectivity. Notably, despite inter-individual variability, connectomes exhibit a substantial number of shared topological features across datasets. Many cycles emerged at consistent ranks across subjects, suggesting that their formation follows overarching principles of brain organization (Figure 2).

In the case of schizophrenia, differences between the SCH and HC groups were evident both globally, via persistence landscapes (Figure 3), and locally, through sublandscape analysis (Figure 4). In SCH subjects, cycles exhibited later birth times and delayed deaths, consistent with the white matter disconnectivity hypothesis [Glahn et al., 2008, Ellison-Wright and Bullmore, 2009,Fornito et al., 2009,Schmitt et al., 2011,Vitolo et al., 2017,Kelly et al., 2018]. Interestingly, the significantly different sublandscapes involved cycles containing regions previously implicated in schizophrenia (see Table 1).

We note that the cycles with significantly different persistence landscapes are always found in subpopulations, never in all participants, however, all schizophrenia participants possess at least one of the significant cycles. This heterogeneity reflects the heterogeneity of the vast spectrum of symptoms. Future work involving larger groups of patients with schizophrenia will enable us to link specific mesoscale topological motifs to symptoms, disease subtypes, or treatment responses.

The recurrence of homology class representatives (and their birth and death times) across individuals in both datasets (Figure 5), despite differences in scanning equipment, demonstrates that cycles are meaningful descriptors of white matter connectivity patterns. Each individual has a set of cycles and persistent bar codes which constitute a ‘connectomic fingerprint’. These features could inform future experimental designs aimed at probing schizophrenia-related brain changes and serve as starting points for novel neuropathological investigations.

Persistent homology offers a novel perspective on brain connectivity by focusing on cycles—topological holes that persist across filtration steps. This enables us to indicate which sets of brain regions, and not just single regions, exhibit different properties between the two groups. This is particularly well-suited to studying white matter nerve bundles, which inherently span multiple regions. In the case of schizophrenia, the literature tends to report only which bundles are affected [Kelly et al., 2018] or that the bundles change with age [Cropley et al., 2017]; whereas they rarely specify the precise inter-regional connections involved. Our method enables us to localise the alterations to subsegments of tracts, thereby providing a more precise anatomical characterisation of the affected pathways, rather than reporting only whole tract abnormalities.

Although MesoSCOUT, by design, is robust to monotonic transformations, which mitigates some of the dependency on the quality of the connectivity matrix and, in turn, its dependency on choices of scanning hardware, scanning protocol, acquisition parameters, and preprocessing steps, etc., potential sources of error remain. Further investigation of the susceptibility of our approach to the preprocessing steps would be useful, but would require the availability of datasets where scans have been performed on the same individual with different scanner settings and preprocessing steps, ideally on the same day. We discuss these considerations, along with current limitations and opportunities for refinement, in Supplementary Section B.13.

In summary, MesoSCOUT is a novel, robust approach for producing detailed connectivity profiles of the human brain, providing “connectomic fingerprints” that could serve as a foundation for future biomarker development across a range of brain disorders. Future directions include establishing normative topological connectomic fingerprints in large-scale datasets and exploring their utility as biomarkers across diverse clinical contexts, including early detection, stratification, and treatment monitoring.

## 4 Methods and Materials

We used two datasets: (1) the Centre for Biomedical Research Excellence (*COBRE*) dataset (https://fcon_1000.projects.nitrc.org/indi/retro/cobre.html) and (2) the Human Connectome Project (*HCP*) dataset (https://humanconnectome.org).

### 4.1 Data sets

#### 4.1.1 COBRE dataset

The *COBRE* data set [Aine et al., 2017, Cetin et al., 2014] consists of structural magnetic resonance imaging scans of healthy controls (HC) (*N* = 44, 11 female) and schizophrenia subjects (SCH) (*N* = 44). Data were obtained by querying the SchizConnect database (http://schizconnect.org). The HC group were matched by age and sex to the SCH group as in [Dimulescu et al., 2021].

Data was collected on a Siemens Magnetom Trio 3T MR device with the acquisition parameters presented in Table 4. This study used diffusion-weighted tensor image (DWI) data and high-resolution T1-weighted images; details of data acquisition and initial data preprocessing are outlined in detail in [Dimulescu et al., 2021, Cetin et al., 2014].

#### 4.1.2 Human Connectome Project

Data from 88 healthy subjects (33 female) from the Human Connectome Project [Van Essen et al., 2012] were used (the size matched the number of subjects in the *COBRE* dataset). It was not possible to match subjects according to age as only 13 out of 1133 *HCP* subjects were over 36, while the average age for COBRE subjects is 38.3 years.

A modified Siemens Skyra 3T MRI scanner was used [Sotiropoulos et al., 2013], and the acquisition parameters are presented in Table 4. We refer the reader to [Van Essen et al., 2012] for full details.

### 4.2 Data preparation/processing

#### 4.2.1 Computing brain connectivity matrix

T1-weighted and DTI images were preprocessed with a semi-automatic pipeline from the *FSL* toolbox [Smith et al., 2004, Jenkinson et al., 2012](fsl.fmrib.ox.ac.uk). For the T1 images, this involved the following steps: removal of non-brain tissue, brain extraction, brain mask generation, and brain parcellation according to the AAL2 atlas [Rolls et al., 2015]. Brain extraction was applied to the DTI data, followed by head movement corrections and the removal of eddy current distortions. The probabilistic diffusion model was fit to the data with Bayesian Estimation of Diffusion Parameters Obtained using Sampling Techniques (BEDPOSTX) [Hernández et al., 2013] *FSL* toolbox.

Subjects’ DTI images were linearly registered to their corresponding anatomical T1 images; the anatomical high-resolution masks were transformed into the subjects’ diffusion space, and this was followed by running probabilistic tractography with 5000 samples per voxel using the PROBTRACKX algorithm [Behrens et al., 2007]. The output of this pipeline is a 94 × 94 structural connectivity matrix *M*. The elements of this matrix, *M*_*ij*_, describe the estimated number of connections from brain region *i* to region *j* from the AAL2 atlas, which has 94 distinct brain regions (see Supplementary Table 72).

#### 4.2.2 Symmetrizing and ordering brain region connectivity

The topological features are obtained from the structural connectivity of each subject. This involves three steps described below: (1) computing the connectivity for every pair of regions, (2) ordering/ranking the matrix entries, and (3) running weighted-rank clique filtration on the ordered matrix.

##### Symmetrizing brain region connectivity

In general, a probabilistic fibre-tracking algorithm results in a non-symmetric matrix, *M*, describing the probability of two regions being connected, i.e., the probability of a path from region *i* to region *j* is not the same as that of a path from region *j* to region *i* [Fornito et al., 2016]. However, these matrices are very nearly symmetric (see Figure 7), which justifies our approach of building undirected simplicial complexes instead of directed flag complexes for our analysis. A symmetrized connectivity matrix 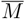 is obtained from *M* the usual way: 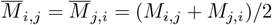.

##### Ordered matrix of brain region connectivity

The order matrix *Ô* is a symmetric matrix of positive integers obtained by replacing the above (or below) diagonal entries of the symmetric matrix 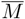 by their ranking from the largest to the smallest (or vice versa):

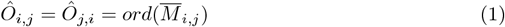

The strongest connection from 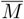 in *Ô* is numbered 1, the second strongest 2, etc (see Figure 8 for an example).

### 4.3 Persistent homology

At the core of persistent homology is the idea of homology groups, which are key topological features of a space. Precise details of the tools (e.g. homology, persistent homology, simplicial complexes, frequency scaffolds) from algebraic topology that we are using are presented in Supplementary Methods Section A.2. Here we outline the main ideas and intuition. Further rigorous mathematical definitions can be found in [Giusti et al., 2015b, Otter et al., 2017, Stolz et al., 2018, Bubenik and Dłotko, 2017]).

#### Simplicial complexes

The structures we work with are simplicial complexes derived from the DTI-data. The building blocks of simplicial complexes are called *n*-simplices, where *n* denotes the dimension of the building block. As shown in Figure 1 a 0-simplex is a node, a 1-simplex is an edge, as is the case in graph theory. TDA goes beyond this and also works with higher-dimensional structures conveying multi-connectivity; so a 2-simplex is formed from 3 connected nodes forming a triangle; a 3-simplex is formed from 4 all-to-all connected nodes (or clique), etc. In general, an *n*-simplex corresponds to a clique with (*n* + 1) nodes. Every subclique of a clique is a lower-dimensional clique.

A simplicial complex *K* is a finite collection of simplices. If a simplex *σ* ∈ *K*, and *σ* ^′^, ≤ *σ*it implies that *σ* ^′^ ∈*K* (here we use ≤ to denote ‘subset of’). Also, if and ^*0*^ are two simplices of *K*, their intersection is either empty or a common face of *σ* and *σ* ^′^.

Homology assesses the existence of features such as connected components, holes, and voids of various dimensions of a space; it keeps track of cavities of various dimensions via homology classes, each of which has a representative ‘cycle’. Cycles are *d*-dimensional loops surrounded by cliques. For any simplicial complex, we can compute its homology classes for each dimension.

#### Weighted Rank Clique Filtration

In this study, the nodes are AAL2 brain regions, and we derive simplicial complexes from the symmetric connectivity matrix (see Section 4.2.2) using a process called the Weighted Rank Clique Filtration (WRCF). A “filtration” refers to a process of creatinga sequence of increasingly detailed simplicial complexes, each containing its predecessor, where the starting simplicial complex consists of *N* disconnected nodes. For the WRCF, at each step, the next connection from the ordered connectivity matrix is added. After a maximum 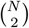 of steps, we obtain a fully connected (*N -* 1)-clique. For our filtrations, the ordering always starts with the strongest connection unless we specify the ‘reverse filtration’.

Homology groups can be computed for each simplicial complex in the filtration. The inclusion maps between successive simplicial complexes induce maps between these homology groups, resulting in a notion of a ‘persistent homology group’ [Zomorodian and Carlsson, 2005] which tracks homology classes that remain non-trivial across the filtration. Cavities corresponding to homology classes and persistent homology classes can be specified by a representative ‘cycle’. PH makes it possible to track the evolution of topological cavities through the filtration

Figure 8 shows a toy example of an ordered 10 × 10 symmetric matrix together with a visualisation of the evolution of the simplicial complexes throughout the filtration. The filtration starts with 10 disconnected nodes; the strongest connection is between nodes *A* and *B* (indicated by ‘1’ in the matrix of integers), the second strongest is between nodes *C* and *D* (indicated by ‘2’), etc. As the filtration progresses and connections are added step-by-step, structure in the form of ‘cycles’ emerges and vanishes.

There are as many steps of the filtration as there are unique values in the ordered matrix *O*, so a maximum of 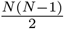 for an *N* × *N* -matrix. For the matrices obtained from the data, the filtration steps were normalised by dividing by the total number of filtration steps, resulting in filtration steps indexed by (rational) numbers in the range [0, 1], enabling comparison between filtrations of different length.

#### Persistence barcodes and persistence landscapes

Given a cycle indexed *i* with birth time *b*_*i*_ and death time *d*_*i*_ in terms of (normalised) filtration steps, we can associate it to an interval called a ‘barcode’, (*b*_*i*_, *d*_*i*_). Hence, for each topological structure (cycle) uncovered using persistent homology, we have its associated barcode, and the results can be summarised in a multiset of *P* persistence barcodes 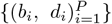. We shall segregate the barcodes by dimensionality of cycle.

It is helpful to further convert barcodes into persistence landscapes [Bubenik and Dłotko, 2017]. This information-preserving transformation enables statistical analyses, which are not possible with a discrete multiset of barcodes. For a barcode with birth time *b* and death time *d*, (*b, d*) with *b*< *d*, a piecewise linear function *f*_(*b,d*)_ : ℝ → [0,*M*] is defined as:

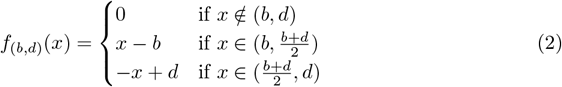

where M is a real number representing the end of the filtration, e.g. number of filtration steps. In this study, we have normalised this to 1. It is now possible to construct a persistence landscape for a multiset of barcodes {(*b*_*i*_, *d*_*i*_), *i* = 1,..., *P* }, as a set of functions λ_*k*_ : ℝ → ℝ defined pointwise, such that for each *x*, λ_*k*_ (*x*) is the *k*-th largest value of 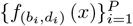 and λ_*k*_ (*x*) = 0 when there is no *k*-th largest value (the index *k* denotes the layer of the landscape).

Unlike a multiset of barcodes, a collection of persistent landscapes can be averaged; the averaging operation is performed layer by layer on the functions _*k*_ for each *k* to obtain an averaged persistence landscape, see Supplementary Figure Supp.A.2 for an example. This allows for statistical analysis of the differences in the persistence properties between various groups.

Differences in landscape structure were investigated using permutation tests on landscape distances [Bubenik, 2015]. The test statistic is the *L*_1_-distances between two averaged landscapes for each dimension *d* compared to the distribution of *L*_1_-distances obtained from randomized landscapes. The details of this procedure are presented in Supplementary Methods Section A.2.4.

### 4.4 Null models

We compared the topological descriptors derived from the data with four types of null models: The ‘Frankenstein-brain’, the “Euclidean brain”, the “Geometric” null model and the Random null model. See Supplementary Methods Section B.8 for details.

For all models (‘Frankenstein-brain’ NM, ‘Euclidean-brain’ NM, Geometric NM, and Random NM), the matrices were of the same size as the connectivity matrices computed from the data (94 × 94).

#### ‘Frankenstein-brain’ null model (random weight permutation)

This null model (NM) utilizes (*COBRE*) population data by altering connectivity matrices through random weight swaps. That is, in a null connectivity matrix of a (simulated) subject, every edge was replaced with a probability *p* with a connection between the same regions, but from a randomly selected participant from within the group, with the selection being made such that all other subjects have the same chance of being chosen. We generated 6 populations of 88 matrices for a range of probabilities *p* ={0.05, 0.1, 0.2, 0.5, 0.9, 0.99 }. The matrices created for the data-based null model were generated and organised into two groups, one from each group of subjects in the *COBRE* dataset. Swapping connections between subjects removes individual contribution while preserving the global relation between brain regions (if there is any).

#### ‘Euclidean-brain’ null model

This NM was constructed based on the concept of a minimally wired brain network [Betzel and Bassett, 2017, Sizemore et al., 2018]. In this model, the connectivity is given by the Euclidean distance between the brain regions of the used atlas. Firstly, 94 brain regions from the AAL2 atlas were represented as nodes, with their spatial positions defined by three-dimensional coordinates given by the centre of the brain region in MNI space [Evans et al., 1992, Brett et al., 2002] The coordinates of the regions from the MNI brain were normalised to lie within the unit cube [0, 1]^3^ for consistency. Secondly, 6 populations, each consisting of 88 spatial distribution matrices, were generated by adding Gaussian noise to the coordinates to introduce variability, each population with mean *µ* =0 but different standard deviation levels, *σ* = {0.0005, 0.005, 0.01, 0.02, 0.05, 0.1 }. Finally, connectivity matrices were constructed with the weight determined by the inverse of the Euclidean distance between the paired nodes: 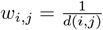, where *d*(*i, j*) is the Euclidean distance between nodes *i* and *j*. This approach emphasises connections between spatially proximal regions, embodying the minimally wired aspect of the network and reflecting spatial constraints in brain wiring. The visualisation of node distribution is presented in the Supplementary Figure Supp.B.30.

#### Geometric null model

This NM, also used in [Giusti et al., 2015b], is where *N* × *N* connectivity matrices *M*^*geom*^ consist of pairwise Euclidean distances between a set of *N* points uniformly distributed in a unit cube in ℝ^*d*^.

#### Random null model

In the Random NM, a connectivity matrix *M*^*rand*^ is a symmetric order matrix where (without loss of generality) the upper diagonal entries have been permuted randomly.

## Supporting information

Supplementary Methods and Results

## Acknowledgements

CM was supported by the Einstein Stiftung Berlin (A-2020-613).

Data used in the preparation of this article were obtained from the SchizConnect database (http://schizconnect.org). Data were provided (in part) by the Human Connectome Project, WU-Minn Consortium (Principal Investigators: David Van Essen and Kamil Ugurbil; 1U54MH091657) funded by the 16 NIH Institutes and Centers that support the NIH Blueprint for Neuroscience Research, and by the McDonnell Center for Systems Neuroscience at Washington University.

All the code was implemented in the Julia [Bezanson et al., 2017] programming language and the persistence homology calculations were performed using the *Eirene* library [Henselman and Ghrist, 2017], supported by the “PersistenceLandscapes.jl” [Dmitruk et al., 2021, Bubenik and Dłotko, 2017] and “DrWatson.jl” [Datseris et al., 2020] libraries. Code reproducing the results of this paper is publicly available in the following GitHub repository: https://github.com/edd26/mesoSCOUT.

