## Supplementary Methods and Results for "MesoSCOUT: A novel tool for revealing mesoscale organization in the white-matter connectome"


October 10, 2025

1046 # A Supplementary methods

1047 ## A.1 MesoSCOUT: Topological Connectome Pipeline

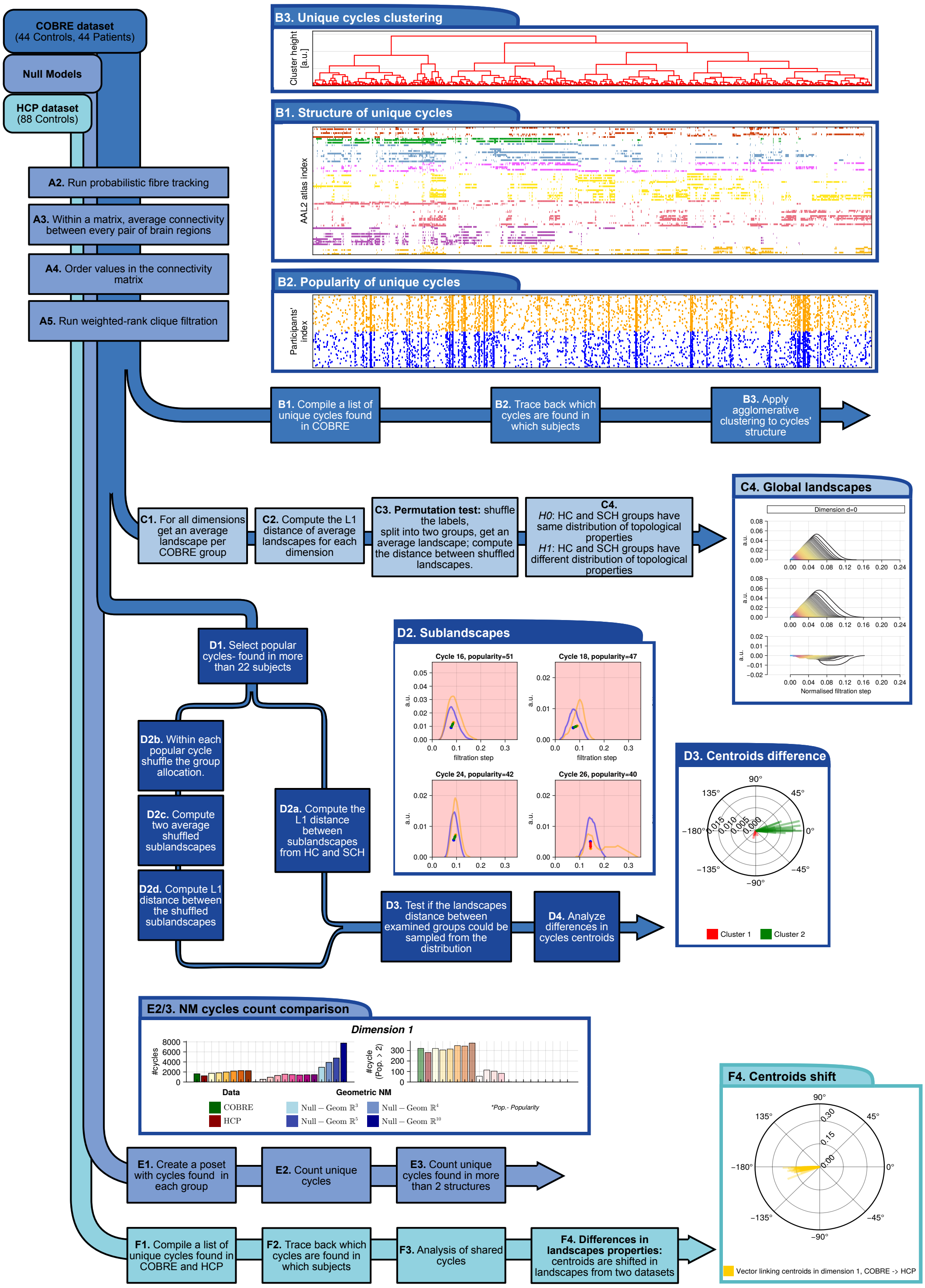


Figure Supp.A.1: MesoSCOUT processing pipeline illustrated for schizophrenia. Boxes represent the stages of data processing. Two datasets were used in the study- *COBRE* and *HCP*; for details, please see Sections 4.1.1 and Section 4.1.2. Processing is grouped into the following sections: A- Data preprocessing leading to topological invariants derived from the connectivity matrices; B- Analysis of cycle classes in *COBRE* and *HCP*; C- Analysis of global landscapes for *COBRE* data; D- Study of topological properties of popular cycle classes in *COBRE*, dimension 1. E- Study of topological properties of cycles found in multiple subjects. The plots presented here are explained in detail in the following Figures: B1, B2, B3- Figure 2; C4- Figure 3; D2- Figure Supp.B.5; D3- Figure 4; E2/3- Figure Supp.B.20; F4 Figure Supp.B.14.

#### A.2 Persistence homology

##### A.2.1 Filtrations of simplicial complexes and persistent homology

Any $N \times N$ symmetric (real) connectivity matrix $M$, can be related to a filtration of simplicial complexes. We shall briefly outline the main steps here, a detailed exposition can be found in [Giusti et al., 2015b, Giusti et al., 2015a]. Whereas $M$ can be considered the adjacency matrix of an undirected weighted network graph, we shall examine a nested sequence of undirected graphs that can emerge from $M$ based on a parameter, connectivity strength. We then obtain the clique complex associated to each graph in this nested sequence, and catalogue the persistent homology cycles.

The distinct upper-diagonal entries of $M$, $\{M_{ij}\}_{i<j}$ can be ordered from the largest to the smallest (or vice versa) and let $q$ be the number of distinct entries (so $q \leq \binom{N}{2} = \frac{N(N-1)}{2}$). This gives us a sequence of nested simple graphs $G_r$ that comprise the order complex of $M$, $\mathrm{ord}(M)$:

$$\mathrm{ord}(M) = (G_0 \subset G_1 \subset \ldots \subset G_r \subset G_{r+1} \subset \ldots \subset G_q), \tag{A.1}$$

where $G_r$ is the graph consisting of $N$ nodes and all the edges ranked 1 to $r$. $G_0$ is the graph with no edges. The integer $r$ is denoted the filtration step and $0 \leq r \leq q$. The subsequent graph, $G_{r+1}$ can be obtained from $G_r$ by adding the edge corresponding to the next-largest entry in $M$, so $G_r \subset G_{r+1}$. This process is continued until the fully connected graph, $G_q$, is reached. Note that for each filtration step, several edges may be added to the graph. In practice, for real-valued matrices obtained from DTI connectivity data, the non-zero entries are in general distinct and the final step $q$ where all the last ranking (zero entries) are added at once, can differ. We normalise our filtration steps to values in $[0, 1]$ by dividing $r$ by $q$.

The clique complex associated to a graph $G$ is a simplicial complex with the same vertex and edge set as the graph. The clique complex, denoted $X(G)$, of a graph $G$ with $N$ vertices, is the set of all cliques (all-to-all connected vertices) of $G$:

$$X(G) = \{\sigma \subset [N] \mid \sigma \text{ is a clique of } G\}.$$

Let $X_r(G)$ denote the set of $(r+1)$-cliques of $G$. ($[N] = \{1, 2, \ldots, N\}$ is the ordered set of $N$ vertices). Formal linear combinations of cliques are called chains and form vector spaces over a field $\mathbf{k}$. Given a set of $(i+1)$-cliques of G, $\sigma_1, \ldots, \sigma_l$, we can define the $i$-th chain group, a $\mathbf{k}$-vector space as follows:

$$C_i(X(G), \mathbf{k}) := \left\{ \sum_{j=1}^{\ell} a_j c_{\sigma_j} \mid \sigma_j \in X_i(G) \text{ and } a_j \in \mathbf{k} \text{ for each } j = 1, \ldots, \ell \right\},$$

where $c_{\sigma_j}$ is the basis element of the vector space $C_i(X(G), \mathbf{k})$ corresponding to the clique $\sigma_i$.

The maps $\partial_i$ are the standard boundary maps $\partial_i : C_i(X(G)) \to C_{i-1}(X(G))$, for $i > 0$ are defined as the following alternating sum of sub-(i-1)-cliques:

$$\partial_i(c_{v_0 v_1 \ldots v_i}) = \sum_{k=0}^{i} (-1)^k c_{v_0 v_1 \ldots \hat{v}_k \ldots v_i},$$

where $c_{v_0 v_1 \ldots \hat{v}_i \ldots v_i} = c_\sigma$ and the $i$-clique $\sigma = \{v_j\}_{j=0}^{i}$ has been specified by its constituent vertices. Note that $v_0 v_1 \ldots \hat{v}_k \ldots v_i$ denotes the vertices $v_0 v_1 \ldots v_i$ with the vertex $v_k$ missing. This map, defined on the basis elements, can be extended to general chains via linearity. The map $\partial_0$ is the zero map.

The boundary of a boundary is always zero, i.e. $\partial_i \circ \partial_{i+1} = 0$; this enables us to define homology groups of $X(G)$ in the usual way:

$$H_i(X(G), \mathbf{k}) := \frac{\ker \partial_i}{\operatorname{im} \partial_{i+1}}. \tag{A.2}$$

Cycles are representatives of homology classes. Note that cycles are defined to be chains with no boundary that are not themselves the boundary of a higher-dimensional chain.

Given a symmetric matrix $M$ with order complex $\mathrm{ord}(M)$, the $i$-th Betti curve of $\{X(G_0), X(G_1), \ldots, X(G_q)\}$ derived from is defined by:

$$\beta_i(r) = \beta_i(X(G_r)) := \text{rank}\, H_i(X(G_r), k),\ \ j \in [0, q], \tag{A.3}$$

where $H_i(X(G_r), \mathbf{k})$ is the $i$th homology group of $X(G_r)$ with coefficients in the field $\mathbf{k}$ (the software implementation uses $\mathbb{Z}_2$ coefficients).

The inclusion maps $\iota_r : X(G_r) \hookrightarrow X(G_{r+1})$ induce maps on the homology

$$(\iota_r)_i : H_i(X(G_r), k) \to H_i(X(G_{r+1}), k).$$

These induced maps allow us to follow cycle representatives of homology classes as they evolve through the filtration, from one complex to the next. In particular, we note when they are 'born' 'birth time' $b$ (the value of the filtration when it first appears) and when they die 'death time' $d$ (the value of the filtration when it ceases to exist).

##### A.2.2 Frequency scaffolds

The frequency homological scaffold $\mathcal{H}_G^f$ [Petri et al., 2014] is an object that was used to compactly represent the homological features of popular cycles from the *COBRE* dataset ($f$ is used to note the frequency homological scaffold, with $p$ being a possible alternative for the persistent homological scaffold.) For a given graph $G$ (e.g. a weighted graph where weight is the count of connections in cycles), the frequency scaffold quantifies edge importance by counting the number of distinct topological cycles in which each edge participates. The edge weight $\omega_e^f$ is defined as:

$$\omega_e^f = \sum_{g_i | e \in g_i} \mathbf{1} e \in g_i \tag{A.4}$$

where $\mathbf{1}e \in g_i$ represents as the indicator function for the set of edges composing a cycle generator $g_i$. This formulation allows for a comprehensive quantification of an edge's topological significance by measuring its participation across multiple homological cycles.

##### A.2.3 Persistence (sub)-landscapes averaging

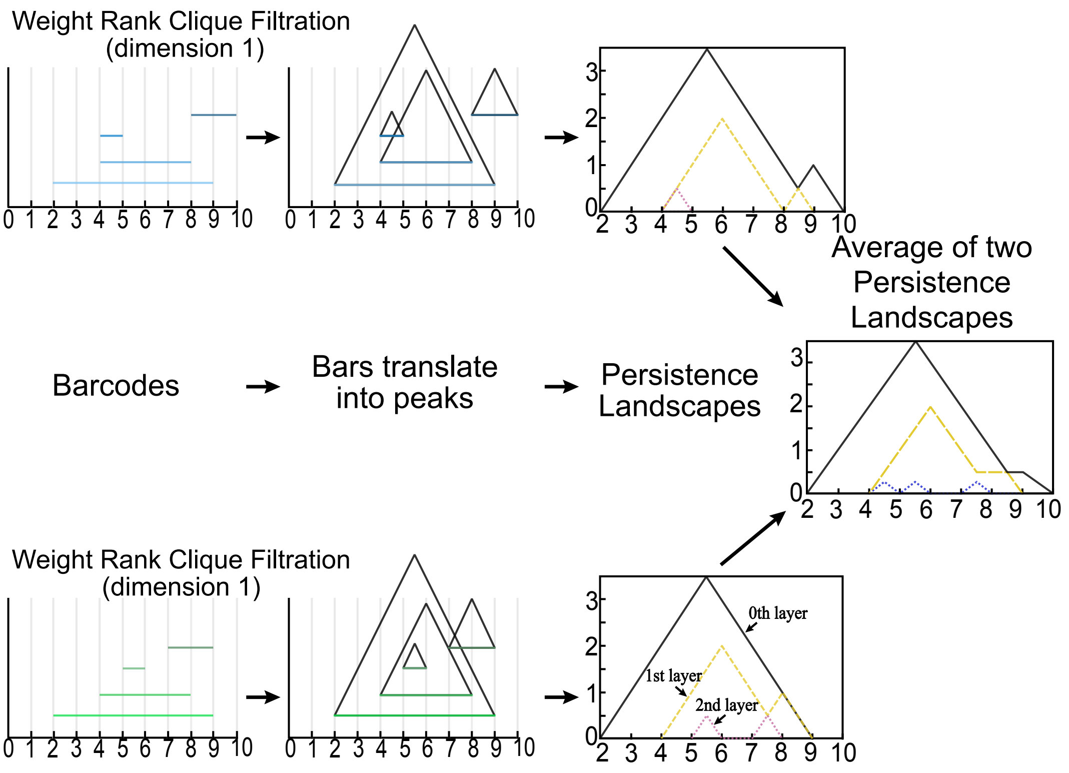


Figure Supp.A.2: Visualisation of landscape averaging. Figure adapted from [Stolz et al., 2017].

##### A.2.4 Statistical tests on persistence (sub)-landscapes

For each dimension, $D$, and two groups, $A$ and $B$ (i.e. HC and SCH), their landscapes denoted $Y_D^A$ and $Y_D^B$ respectively, the distance $W_D$ can be computed using the $L_1$-distance:

$$W_D := L_1\left(Y_D^A, Y_D^B\right). \quad \text{(A.5)}$$

The landscape distance distribution is created by computing the $L_1$ distance between two averaged landscapes $\hat{Y}_D^1$ and $\hat{Y}_D^2$, where $\hat{Y}_D^1$ is created from a random selection of half of the landscapes from both groups; $\hat{Y}_D^2$ is created from the remaining landscapes. This process is then repeated $m = 10000$ times using equal partitions of subjects (10000 permutations). The histogram of these distances provides us with an empirical distribution of persistent (global) averaged landscape distances, $W$, which we shall use as the null distribution in hypothesis testing. The criterion for accepting the alternative hypothesis (the landscapes are significantly different) is that the distance falls in the right tail of the $W$ distribution of distances, with the critical value $\alpha = 0.05$. The $p$-value is computed as the proportion of distances that are as extreme as or more extreme than the landscape distance observed between the two averaged landscapes computed from the original groups, $W_D$, that is

$$p = \frac{|\mathcal{U}_D|}{m} \quad \text{(A.6)}$$

where $|\mathcal{U}_D|$ is the cardinality of the empirically derived set, $\mathcal{U}_D$, of distances greater or equal to $W_D$.

The above permutation test can also be used on averaged sublandscapes in an analogous way. Where each subject's single-barcode landscape, or sublandscape, for a popular shared cycle is used in the averaging procedure. If a subject does not have a particular cycle, then a "flat landscape", i.e. $f_{b,d} = 0$, is used.

#### A.3 Processing the results

**Agglomerative clustering** Agglomerative clustering was used to highlight similarities between cycles' underlying nodal structures. Each cycle was represented by a binary vector of length 94, whose $i$-th entry was 1 if brain node $i$ participates in the cycle, and 0 otherwise. The Hamming distance between two cycles was then computed to quantify their structural dissimilarity. Finally, agglomerative clustering with Ward's minimum variance method [Ward, 1963] was used to cluster the cycles based on their Hamming distances, highlighting sets of cycles with similar node-participation patterns.

**Global persistence landscapes and persistence sub-landscapes** Persistence properties of cycles were analysed at two levels: collectively (via global persistence landscapes) and on an individual cycle level (via persistence sub-landscapes).

A subject's persistence landscapes were created from the topological information derived from the subject's ordered connectivity matrix. For each dimension, $d$, a global persistence landscape was derived from all the $d$-cycles. These global landscapes could then be averaged across groups of subjects, e.g. all subjects with schizophrenia.

Analogously, for each distinct cycle, averaged sublandscapes could be obtained by taking the sublandscape (or single barcode landscape) corresponding to that cycle from each subject and then averaging (the sublandscape was flat if an individual did not have a particular cycle). This enabled decomposition of the global comparison to a cycle-specific sublandscape comparison between the SCH and HC groups in the *COBRE* dataset.

**Landscapes centroids** To highlight the differences between averaged sublandscapes obtained from the SCH and HC groups, we compared their centroids. The centroid $(\bar{x}, \bar{y})$ of the region bounded by the uppermost layer of each persistent sublandscape $f(x)$ and the bounded interval $[0, 1]$ (the range of normalised filtration steps on the x-axis) was computed as follows:

$$\bar{x} = \frac{1}{A}\int_0^1 x f(x) dx, \;\; \bar{y} = \frac{1}{2A}\int_0^1 f^2(x)\, dx; \;\; A := \int_0^1 f(x)\, dx. \quad \text{(A.7)}$$

### B Supplementary results

#### B.1 COBRE cycles

##### B.1.1 Dimension 2 and 3 cycles

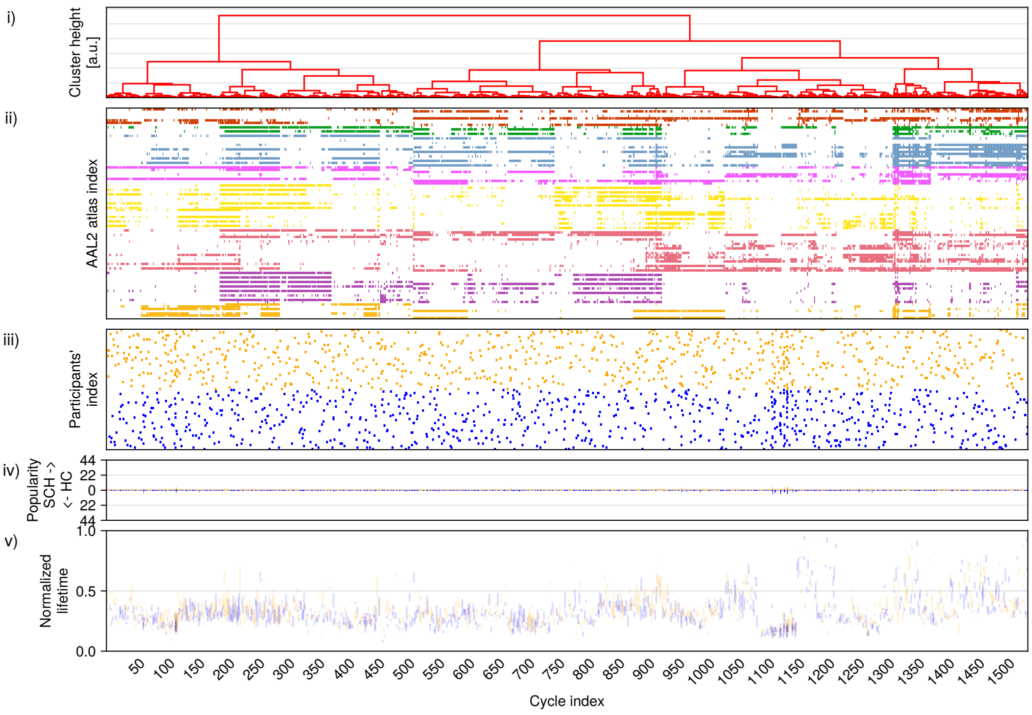


(a) Dimension 2

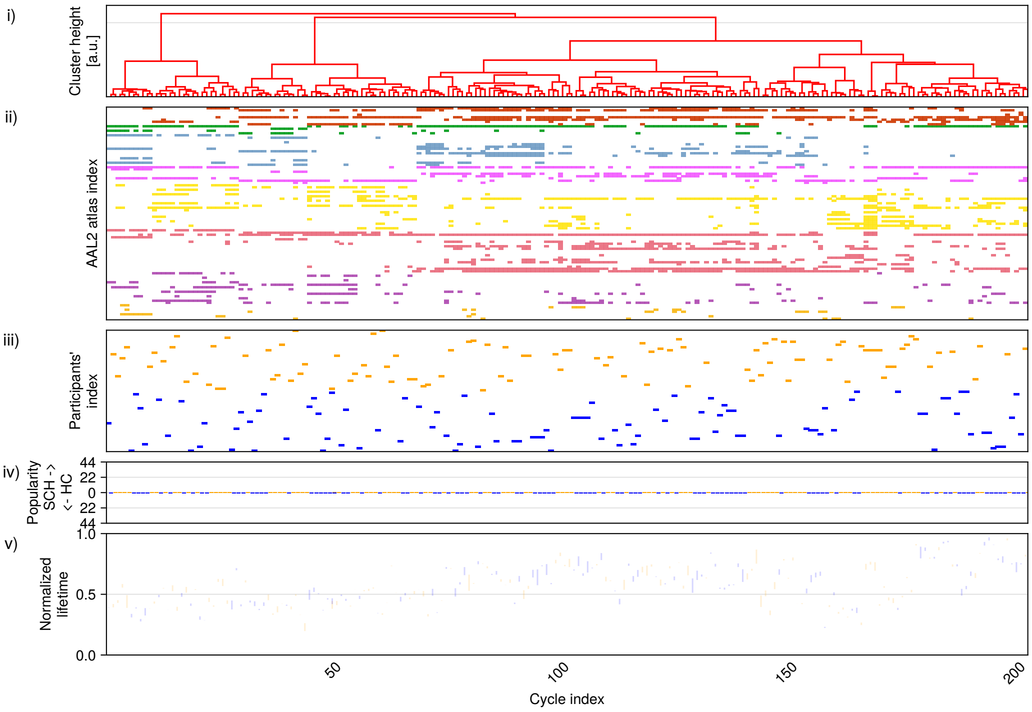


(b) Dimension 3

Figure Supp.B.3: Exploration of structure, popularity, and topological properties of unique cycles in dimensions 2 and 3. The plot description is the same as in Figure 2.

##### B.1.2 Clustering of dim 2 cycles

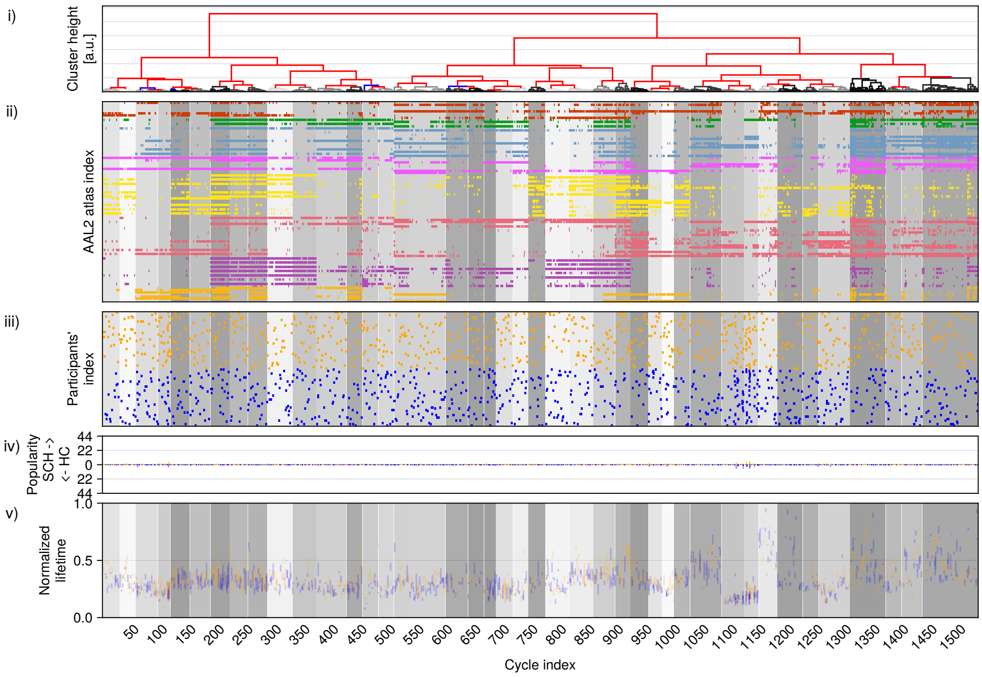


Figure Supp.B.4: Unique cycles structures and popularity presented as a raster plot for dimensions $D = 2$ for COBRE with possible clustering of the cycles. Clustering, as indicated by the background grey in (ii), (iii) and (v), is created by splitting the Ward clustering tree 6 times from the top or until a cluster contains at least 22 cycles- whichever occurs first. The plots are structured as described in 2. Plots (ii) are raster plots showing which brain regions participated in the unique cycles at the moment of the cycle's birth. All unique cycles are aligned with the the horizontal axis and brain regions are marked with 7 colours according to their Yeo's functional network atlas. The brain regions are ordered according to the hierarchical clustering presented at the top part of the plot. Plots (iii) are raster plots showing, in which subjects a cycle was found (orange used for SCH, blue for HC). Plot just below summarizes individual cycle's popularity, i. e. total number of subjects, in which a cycles was found. Figures in (v) present the (vertical) persistence barcodes for each cycle. The vertical axis shows the lifetime normalised to the maximal filtration step in the filtration. All horizontal axis are the same, using cycle indexing from hierarchical clustering.

#### B.2 Persistence sublandscapes for popular cycles in dimension 1

##### B.2.1 Centroids comparison

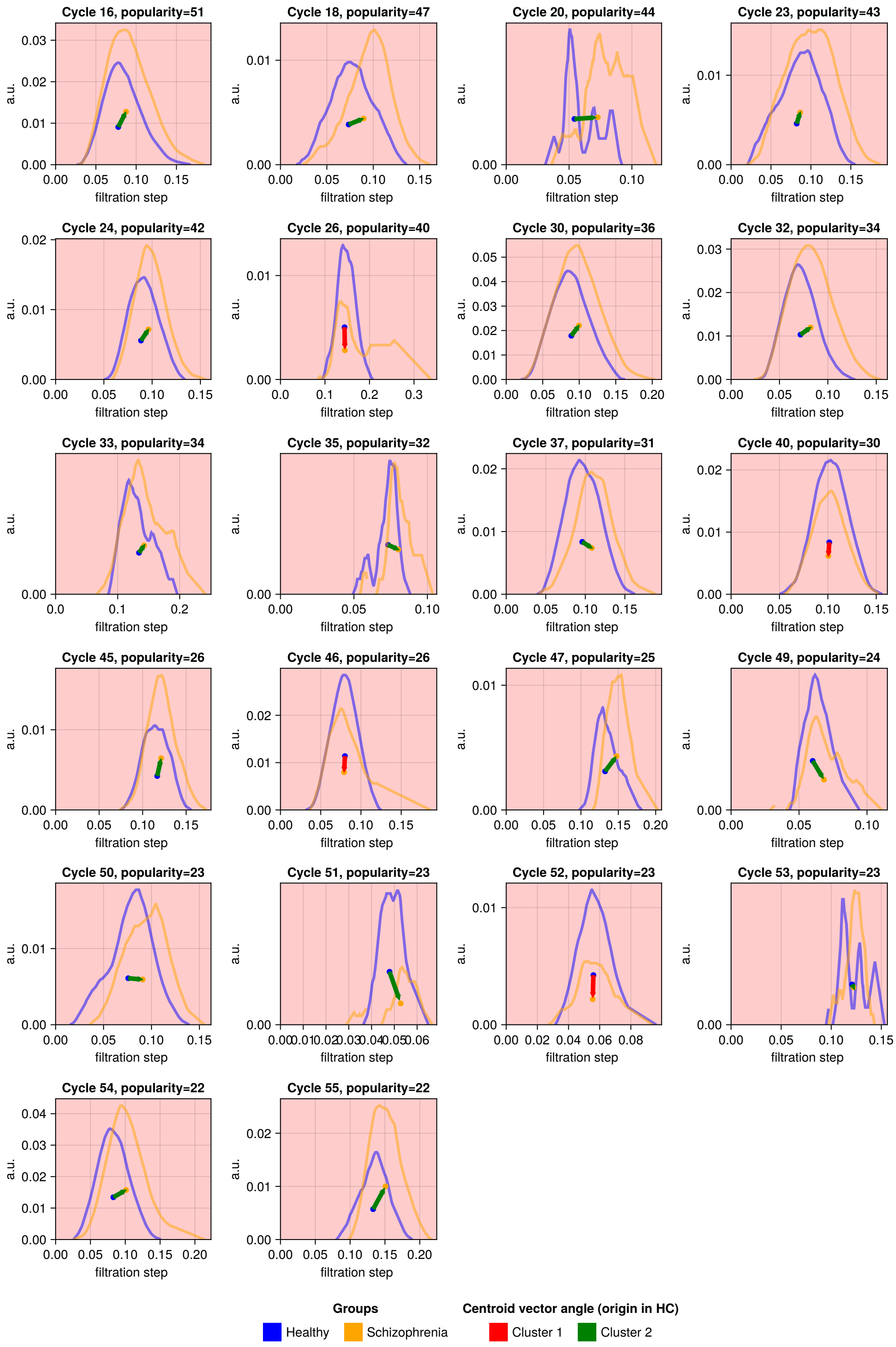


Figure Supp.B.5: Close up on the 22 sublandscapes that are significantly different between the groups. The blue colour is used for the average landscape for healthy controls, orange is used for average landscapes for subjects with schizophrenia. The dots in the middle of the sublandscapes mark the centroids; the vector between the centroid of the HC sublandscape and the SCH sublandscape is shown. By applying agglomeration clustering to the coisne distance matrix computed for all pairs of vectors, we found 2 distinct subgroups of landscapes- for 18 sublandscapes the centroid of $\hat{Y}^i_{SCH}$ is positioned to the right from centroid of $\hat{Y}^i_{HC}$ for cycle $i$ (green colour). For 4 sublandscapes centroid of $\hat{Y}^i_{SCH}$ is positioned above $Y^i_{HC}$, marked with red colour. Another perspective is presented in Figure 4.

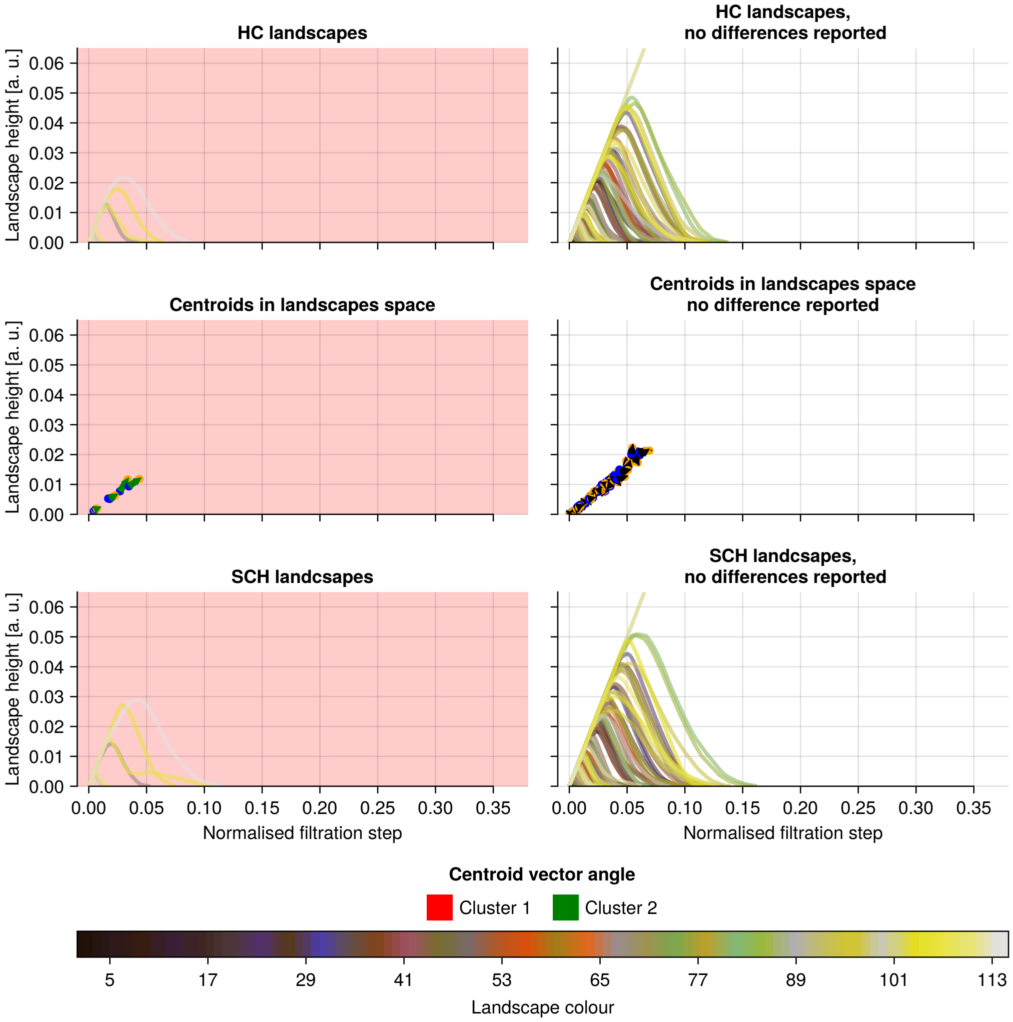


Figure Supp.B.6: Decomposition of the global landscape in terms of popular sublandscapes (data for cycles in dimension 0). In the middle plot, every arrow indicates how the centroids in dimension 0 are dislocated for the 115 popular persistence landscapes that were found to be significantly different between the groups (popular cycles were cycles found in at least 22 subjects). In the top and bottom rows, persistence sublandscapes are presented for the HC group (top) and the SCH group (bottom), for statistically different cycles (first column) and for cycles where no significant difference was found (second column). Persistence sublandscapes are coloured according to the cycle they are derived from, i.e. the same colour between rows indicates the same cycle.

##### B.2.2 Cycle laterality

In Figure Supp.B.7, we decompose the frequency scaffold of popular cycles into one consisting of significant cycles and the other of non-significant cycles. Out of the 123 edges in the significant cycle scaffold, only 6 link the two hemispheres of the brain and do so symmetrically, so they are likely part of the corpus callosum.

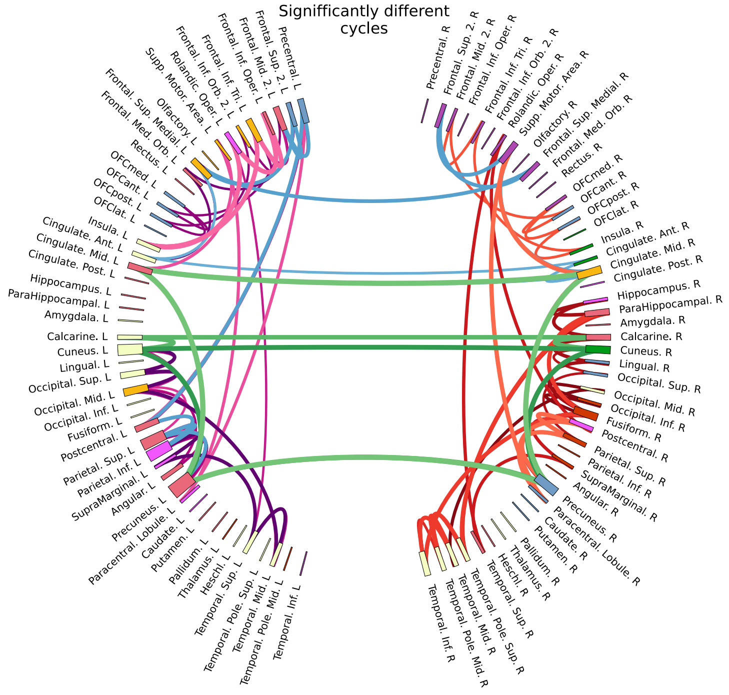


(a) Significantly different cycles

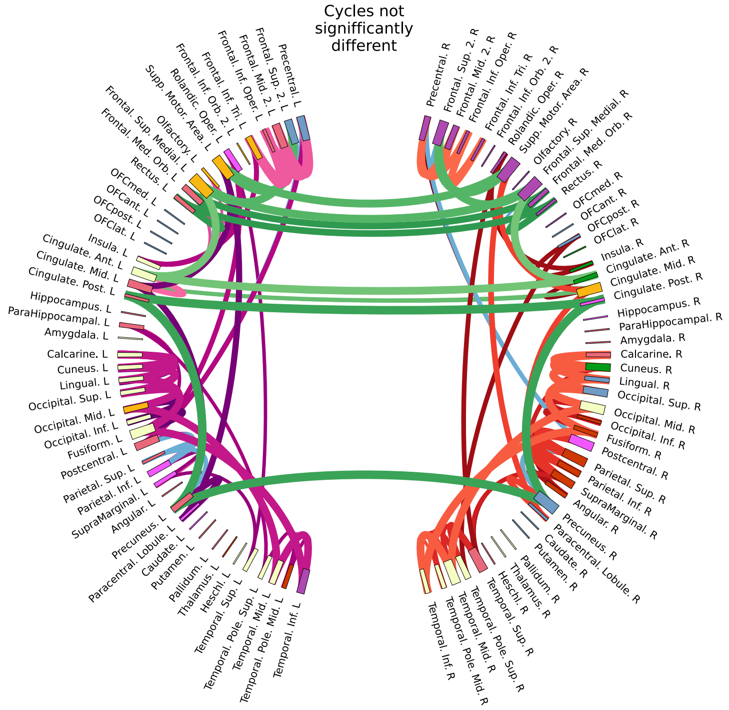


(b) Cycles not significantly different

Figure Supp.B.7: Frequency scaffolds [Petri et al., 2014] created from popular cycles of dimension 1. The scaffold created with significant cycles is presented in the (a); the scaffold created with the remaining popular cycles is presented in (b). Only cycles found in at least 22 subjects are shown. Nodes are coloured according to Yeo network (See Figure 2a for legends) and their size is proportional to number of connections formed. Colours of edges indicate if a cycle is lateralised or across hemispheres: left-only cycles are marked with shades of purple and pink, right-only cycles marked with shades of red and orange, symmetric cycles marked with shades of green, mixed cycles marked with shades of blue. Thickness of connection is proportional to cycle's popularity.

1161 A study by Sizemore et al. (2018) [Sizemore et al., 2018] investigated cycles in the structural connectome of healthy people up to dimension 2. They report 20 2D cavities and 2 3D cavities (described with dimension 1 and 2 cycles, respectively). Among one-dimensional structures, there are cycle representatives that are symmetric (1 cycle), lateralised (9 cycles formed from right-only regions, 4 from left-only regions), and mixed (6 cycles), which demonstrates similar types of structures that we observed. Additionally, 3 of those are found in the popular cycles we report (cycles 16, 37, 47). We demonstrate that persistent features are present in dimension 3 (74 out of 88 subjects had at least 1 cycle). However, their total number is much lower than in lower dimensions, i.e. 202 for dimension 3 and 1625 for dimension 1. We also found 18 cycles in dimension 4 but did not include them in the analysis due to their sparse popularity. Computations for higher dimensions were not feasible due to computational limitations.

We report the following pairs of twin cycles (cycles that were found to be significantly different are marked with $s$): 1 and 8, 5 and 6, 9 and 13, 11 and 12, 14 and 27, 26($s$) and 42, 34 and 47($s$), 37($s$) and 48.

#### B.3 Further analysis of centroid clusters and cycles

In the first of the patterns shown in Figure 4(b), *SCH* cycles are more persistent while being born at a similar time step as *HC* (like landscapes 16, 23, 24, 30, 32, 33, 45) (see Supplementary Figure B.2. The structure is formed at the same filtration step, but the killing of the cycle happens later because the interconnection between the nodes is weaker. In the second, the *SCH* cycles are later born than in *HC* (landscapes 18, 20, 35, 37, 47, 49, 50, 54, 55), which means they are born with weaker connections, or there are other, stronger connections formed before the cycle is formed in the schizophrenia group. For example, cycle 16 (middle cingulate & paracingulate gyri, precuneus, all bilateral) appears at the same filtration step in both groups but persists longer in *SCH*, whereas cycle 18 (cuneus & precuneus, both bilateral) and cycle 37 ( middle occipital gyrus, inferior occipital gyrus, fusiform gyrus, middle temporal gyri, inferior temporal gyri, all right) are born at later step in *SCH*, showing delayed cycle formation.

#### B.4 Dimension 1 cycles, brain regions and connections with literature

The *left precuneus* region (found in 8 cycles) shows several abnormalities in individuals with schizophrenia. This region has decreased centrality in some schizophrenia patients with the genetic subtype, 22q11.2 deletion syndrome [Ottet et al., 2013]. Zhao et al. [Zhao et al., 2018] reported that regional homogeneity (ReHo) measures in the left precuneus are altered in schizophrenia when compared to healthy controls. Additionally, the precuneus is a central hub in the default mode network (DMN) [Utevsky et al., 2014], so changes to the connectivity of this region, as evidenced by white matter changes, may affect how easy it is for the brain to reach the DMN. Increased functional connectivity of the *left precuneus* with other DMN brain regions was reported by [Sambataro et al., 2010] in schizophrenia subjects compared to controls.

The *left* or *right cuneus* regions, although they were found in multiple significantly different cycles, do not appear to be mentioned in studies on structural changes in schizophrenia. However, the *cuneus* is in the occipital lobe and is involved in visual processing, so there may be a relation to the symptom of experiencing hallucinations [Tohid et al., 2015]. There is some evidence from functional studies that subjects with schizophrenia exhibit reduced activation in *right cuneus* in episodic retrieval memory tasks [Ragland et al., 2009].

The third most frequently found region was the *left superior parietal gyrus*, which also exhibits abnormalities in individuals with schizophrenia [Wheeler and Voineskos, 2014]. SCH group is characterised by decreased centrality [Ottet et al., 2013] in this region compared the HC group and Howes et al. 2023 [Howes et al., 2023] mentions lowered levels of N-acetylaspartate (NAA) in the magnetic resonance spectroscopy (MRS) studies in chronic schizophrenia patients (the NAA molecule is found in neuronal tissue and is used as a marker because it gives the largest signal in MRS).

The *left inferior parietal gyrus* shows a progressive decline in regional grey matter volumes in subjects with schizophrenia who have experienced psychotic experiences [Howes et al., 2023, Vita et al., 2012, Merritt et al., 2021]. This decline was particularly notable in the temporal, frontal, cingulate, and parietal cortices.

The *right precuneus* region also exhibits abnormalities in individuals with schizophrenia: Cropley et al. [Cropley et al., 2017] found a reduction of grey matter in this region for schizophrenia patients over the age of 30, while Shen et al. [Shen et al., 2023] reported greater sample entropy for schizophrenia.

The *right middle cingulate & paracingulate gyri* region shows reduced volume in individuals with late-onset schizophrenia [Li et al., 2015]. Also, longitudinal studies report increased volume loss in this region in schizophrenia patients [Howes et al., 2023].

The *left middle occipital gyrus* (found in cycles 33, 51, 52, 54) demonstrates grey matter volume decline in schizophrenia [Zhao et al., 2018], which is accelerated after the age of 30 [Cropley et al., 2017]. Furthermore, Howes et al. [Howes et al., 2023] mentioned significantly lower $[^{11}C] - UCB - J$ (a marker of synaptic density used in positron emission tomography) uptake in this region, with supporting evidence from the study conducted by Osimo et al. [Osimo et al., 2019]. White matter volume reduction in SCH was also observed in this region, as indicated by Canu et al. [Canu et al., 2015] after Witthaus et al. [Witthaus et al., 2008].

The *right fusiform gyrus* (found in cycles 26, 32, 37, 47) shows significant differences in individuals with schizophrenia. Howes et al. [Howes et al., 2023] reported lower grey matter volumes in SCH patients. Interestingly, despite the fact that literature reports changes in both *right and left fusiform gyrus*, only the *right fusiform gyrus* was found in significantly different cycles (*left fusiform gyrus* was only found in cycles not significantly different).

The *superior frontal gyrus medial* region exhibits lower grey matter volumes in SCH patients, as reported by Howes et al. [Howes et al., 2023] after Gao et al. [Gao et al., 2018].

All brain regions from the cycles 16, 18 and 24, are connected via the cingulum bundle; most of the connections of cycle 37 are within the inferior longitudinal fasciculus, and both of those white matter tracts are reported in literature to be affected by schizophrenia [Wheeler and Voineskos, 2014, Sun et al., 2015] (for cycle 37, one of the connections lies within the inferior fronto-occipital fasciculus tract, not reported in the schizophrenia literature).

Table 5: Summary of brain regions participating in low-persistent cycles and their alterations. This table lists brain regions found in cycles, the hemisphere they are located in, and the specific alterations reported in the literature.

| Cycle | Region | Side | How Affected |
|---|---|---|---|
| 20 | Inferior frontal gyrus opercular part | L | Significantly lower grey matter volumes [Gao et al., 2018] |
| 20 | Inferior frontal gyrus triangular part | L | Significantly lower grey matter volumes [Gao et al., 2018] |
| 20 | Rolandic operculum | L | - |
| 20 | Insula | L | Reduced grey matter density [Kuo and Pogue-Geile, 2019] and reduced grey matter volume [Glahn et al., 2008] |
| 35 | Rolandic operculum | R | - |
| 35 | Postcentral gyrus | R | Lower grey matter density relative to controls [Glahn et al., 2008] |
| 35 | Supramarginal gyrus | R | - |
| 35 | Superior temporal gyrus | R | Reduction in grey matter density in drug-free individuals [Kuo and Pogue-Geile, 2019] and greater volume loss [Vita et al., 2012, Gao et al., 2018] |
| 53 | Superior frontal gyrus dorsolateral | L | Reduced cortical thickness in chronic schizophrenia [van Erp et al., 2018] |
| 53 | Superior frontal gyrus medial | L | Reduced cortical thickness in chronic schizophrenia [van Erp et al., 2018] |
| 53 | IFG pars orbitalis Med_Orb | L | - |
| 53 | Anterior cingulate & paracingulate gyri | L | Reduced grey matter density [Glahn et al., 2008] and volume [Brugger and Howes, 2017] |

#### B.5 Permutation test results

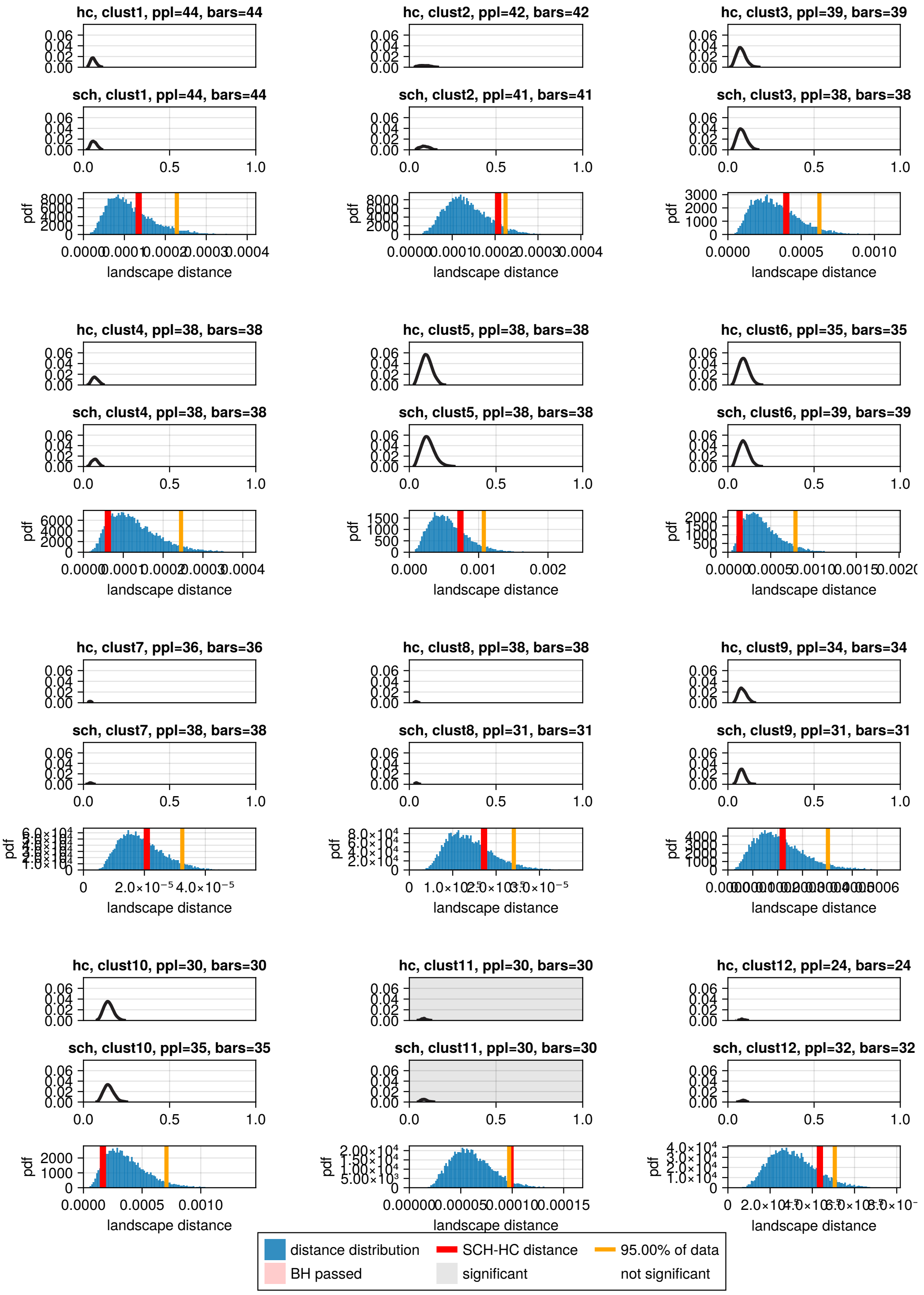


Figure Supp.B.8: Average persistence landsacpes for dimension 1, cycles 1-12. Cycles presented are the cycles that were found in at least 22 subjects

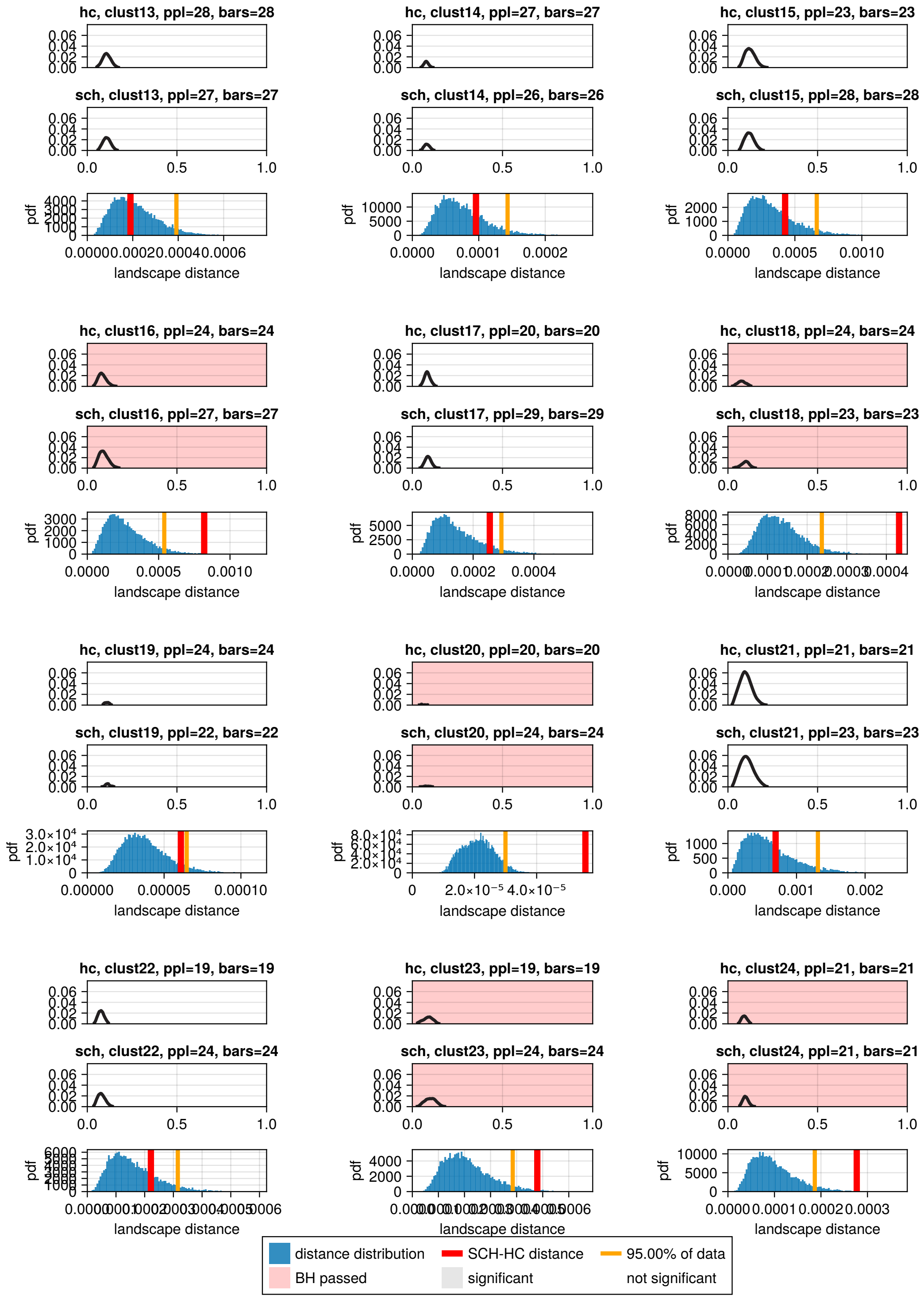


Figure Supp.B.9: Average persistence landsacpes for dimension 1, cycles 13-24. Cycles presented are the cycles that were found in at least 22 subjects

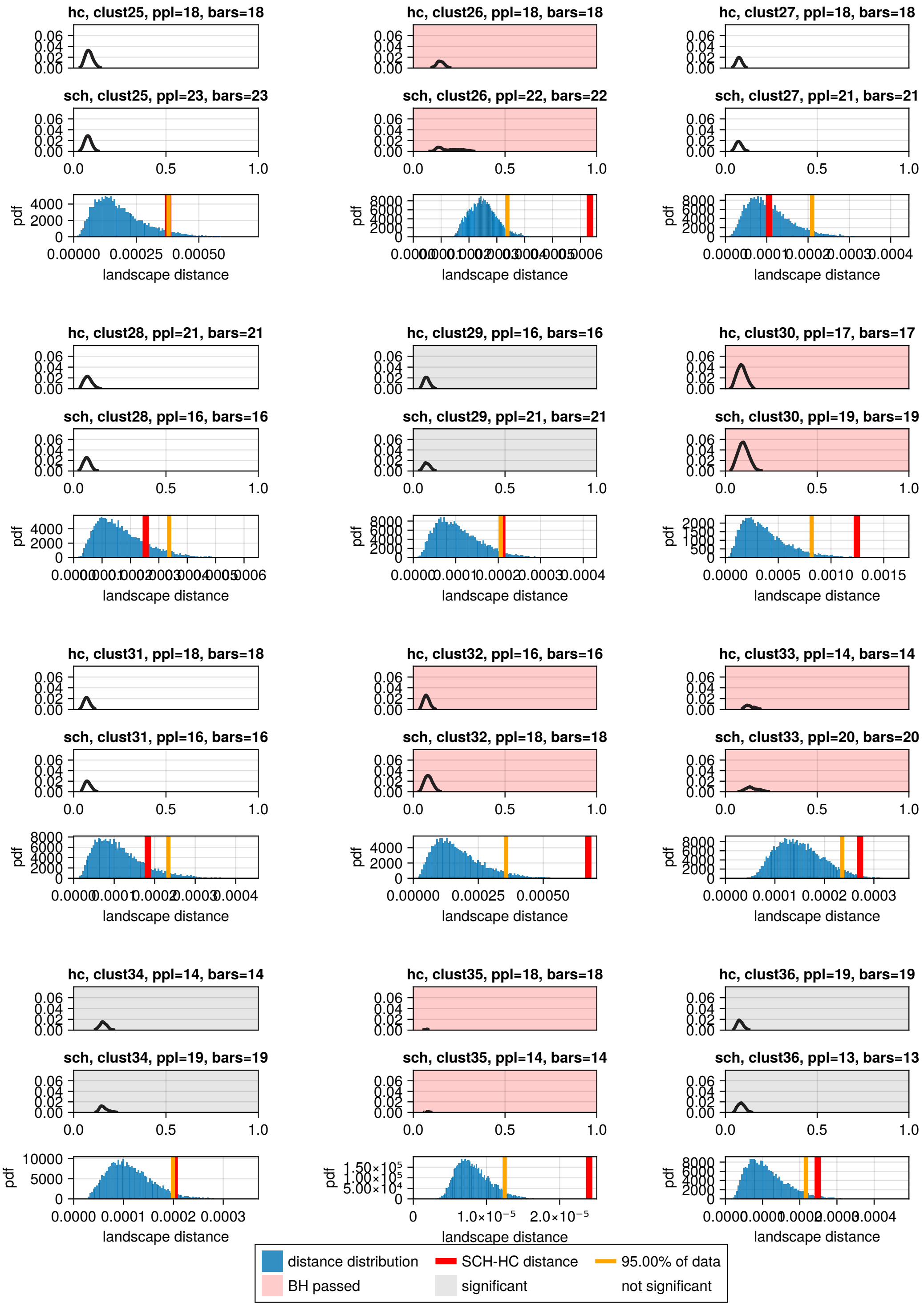


Figure Supp.B.10: Average persistence landsacpes for dimension 1, cycles 25-36. Cycles presented are the cycles that were found in at least 22 subjects

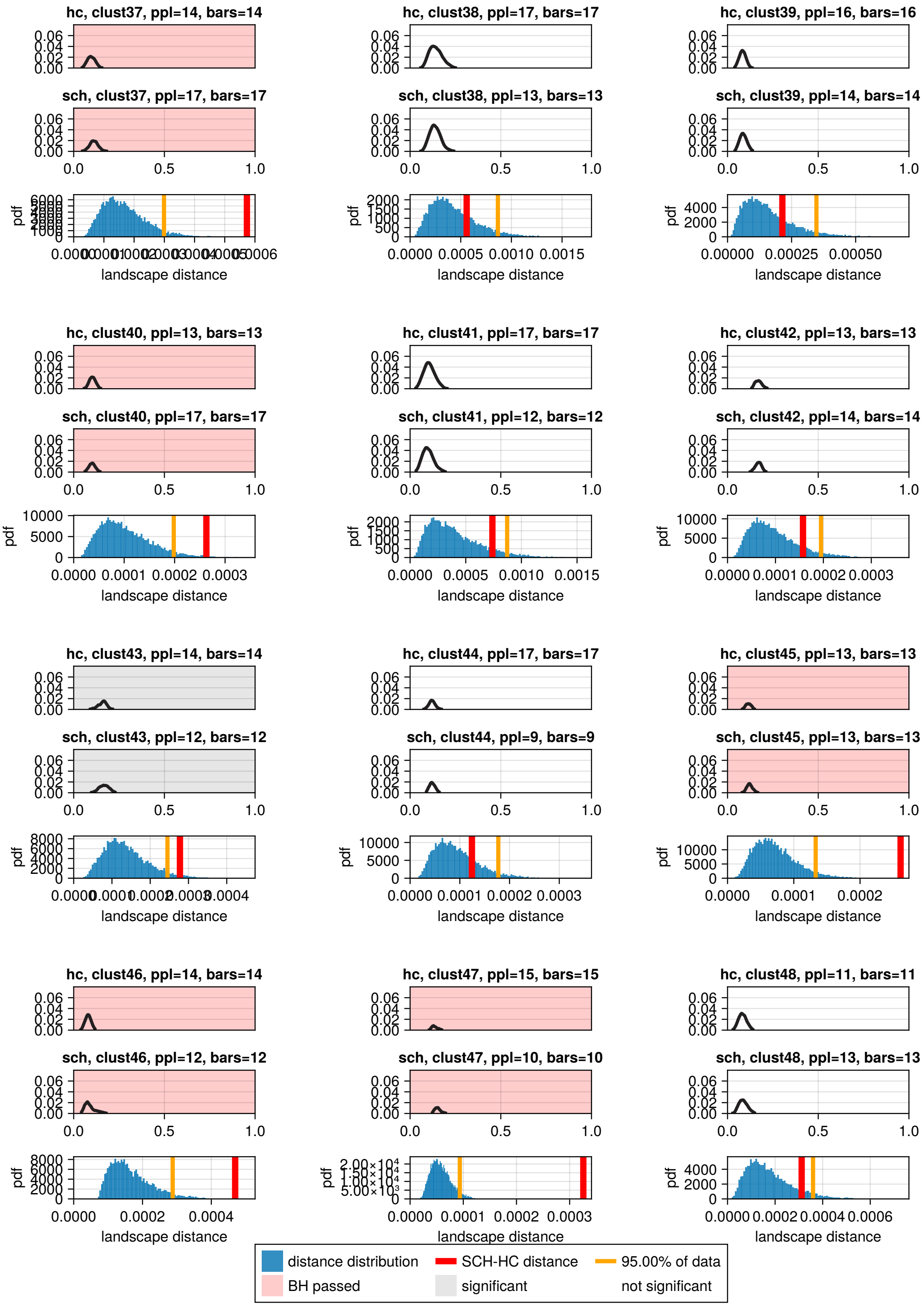


Figure Supp.B.11: Average persistence landsacpes for dimension 1, cycles 37-48. Cycles presented are the cycles that were found in at least 22 subjects

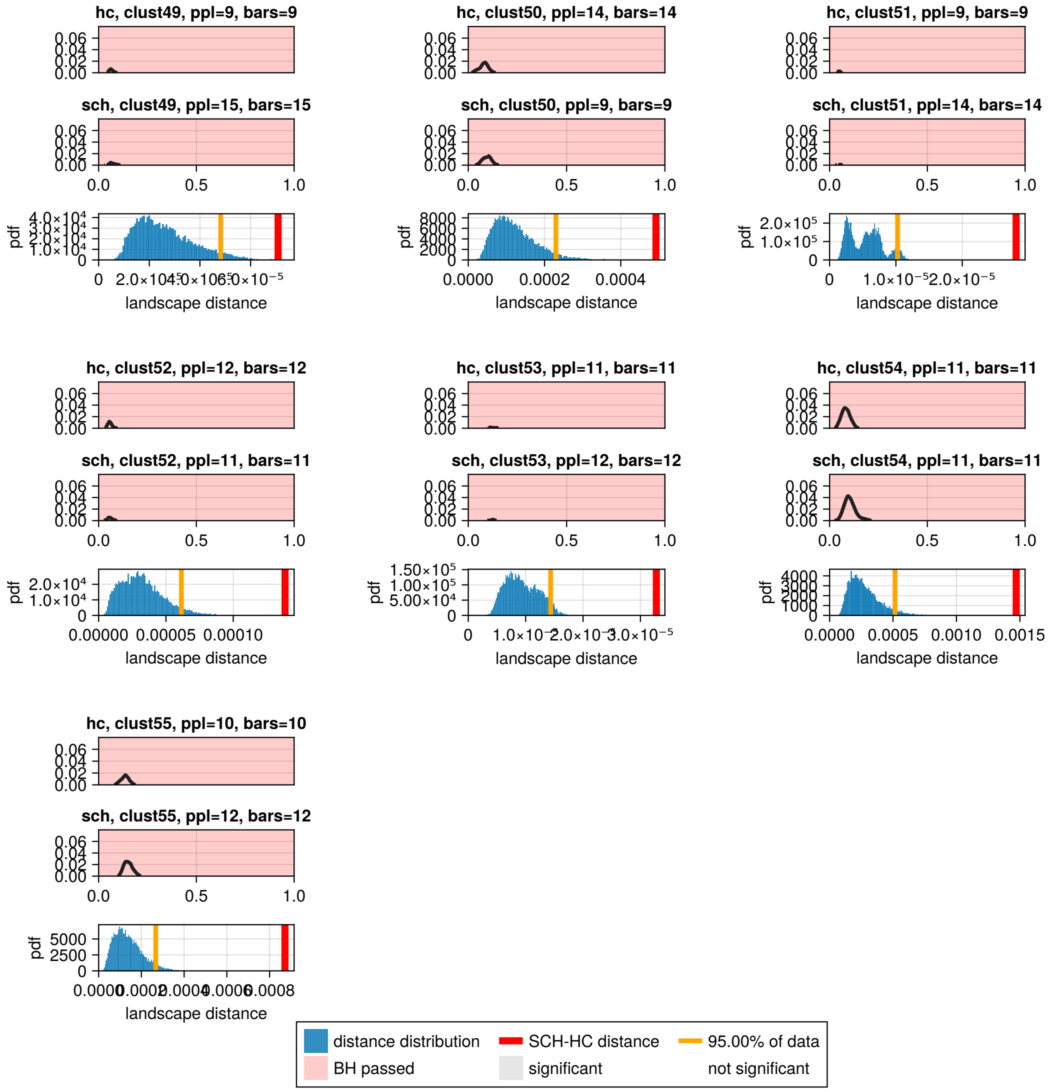


Figure Supp.B.12: Average persistence landsacpes for dimension 1, cycles 49-55. Cycles presented are the cycles tha were found in at least 22 subjects

#### B.6 HCP dataset

##### B.6.1 Sublandscape comparison

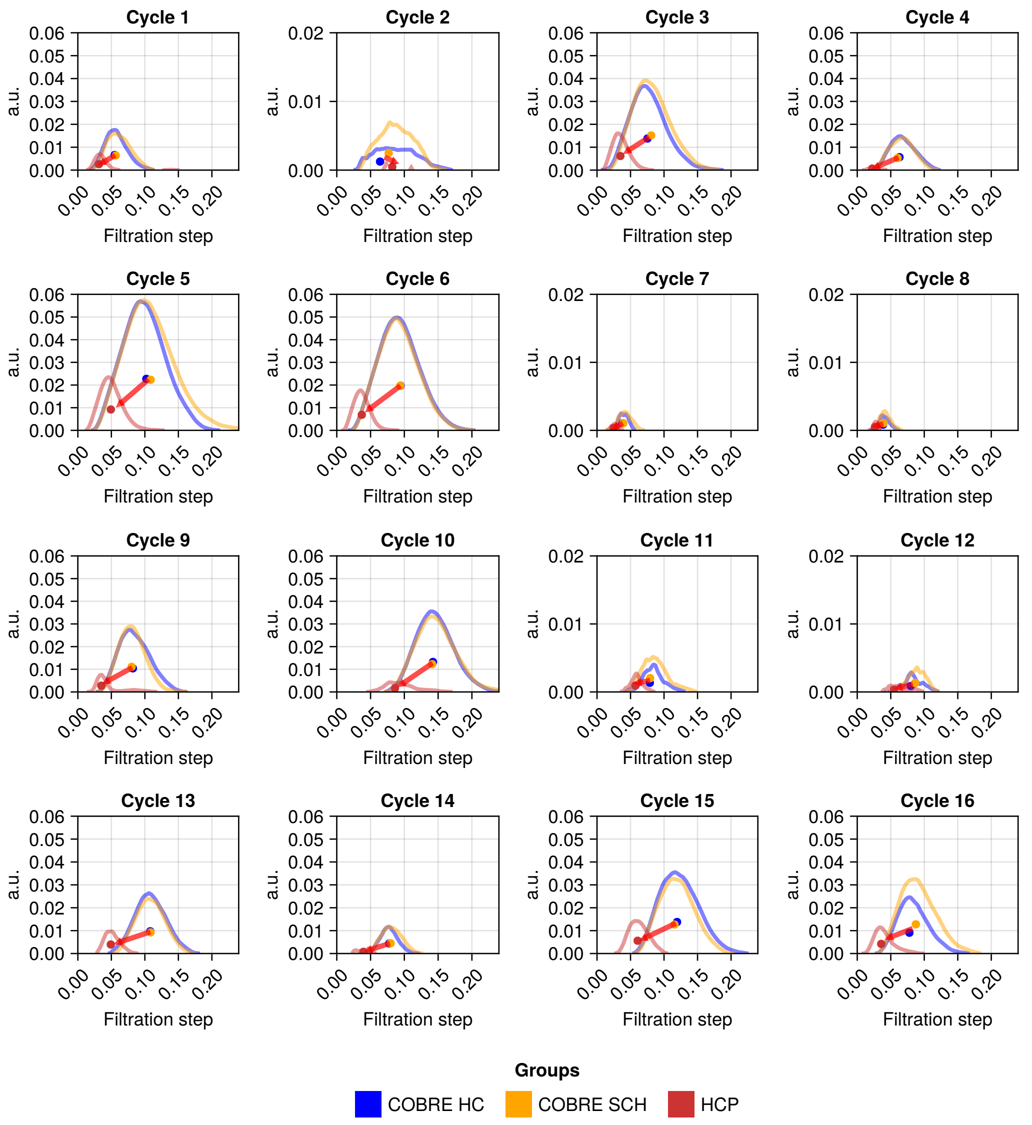


Figure Supp.B.13: Comparison of persistence landscapes for dimension 1 cycles found in two datasets. The cycles shown are 16 most popular cycles from the *COBRE* dataset. Every landscape is the average for one group, as indicated in the inset legend. All landscapes computed from *HCP* data are shifted to the left when compared with persistent landscapes from *COBRE* data, as indicated with the red arrow. The vertical axis represents the landscape's height.

##### B.6.2 Shared cycles

**Addressing the differences between datasets** The differences in cycle persistence between datasets may stem from varying acquisition parameters, which are known to affect the quality of DTI data [Thieleking et al., 2021]. *COBRE* and *HCP* were acquired with different scanners, characterised by different voxel sizes (1.0mm vs 0.7mm for T1 scans; 2.0mm vs 1.25mm for DTI; *COBRE* and *HCP*, respectively; all voxels are isotropic; see Table 4). *COBRE* therefore has lower spatial resolution than *HCP*, (as it has larger voxels), and this results in coarser sampling of fibre tracts and limits the maximum curvature of a tract that can be reconstructed [Caan, 2016]. This leads to *COBRE* having much sparser connectivity matrices. Examples of typical matrices from *COBRE* (HC and SCH) and *HCP* are shown in Figure 7, and the numbers of zero-entries of the matrices are shown in Supplementary Figure Supp.B.19. There are $25.6 \pm 5.7\%$ and $22.6 \pm 5.0\%$ zero entries in the connectivity matrix in *COBRE* (*HC* and *SCH* respectively), while there are almost no zero entries in the matrices from *HCP* (on average: $0.61 \pm 4.53\%$ zeros).

**Sub-landscape comparison between *COBRE* and *HCP*** As there are fewer zero entries in the *HCP* matrix the filtration contains more steps. We therefore expect that the persistence sublandscapes derived from the *HCP* dataset would be shifted toward the beginning of filtration compared to their counterparts from the *COBRE* dataset.

Indeed, this is the case for all shared cycles ($p < 0.01$), as shown in the sublandscapes comparison presented in Supplementary Figure Supp.B.13 (see Supplementary Figure Supp.B.14 for a summary of centroid differences, and Section B.6 for details of statistical tests). The magnitude of this left shift increases with dimension, due to an additive effect - it is only possible for higher-dimensional cycles to emerge when lower-dimensional cycles die.

##### B.6.3 Centroids comparison

A summary for shared cycle sublandscapes, where vectors are drawn from the average centroid of the *COBRE* sublandscape to the centroid of a corresponding *HCP* sublandscape. They are all pointing in the left-downwards direction.

Further, the Kruskal-Wallis test on the distribution of coordinates of centroids showed that cycles found in both *HCP* and *COBRE* have vertical coordinates significantly lower ($p < 0.01$) for all dimensions and horizontal coordinates lower for dimensions 0 and 1. The comparison was not possible for dimension 3, as no cycles are shared between groups. The results of the test are presented in the Table 6.

At the same time, cycles unique to *HCP* (found in *HCP* dataset and not found in *COBRE*), have their birth and death times in the majority placed to the right, compared to the *HCP* cycles that are shared. Visualisation of centroids' position is pictured in Supplementary Figures Supp.B.15 Supp.B.16, Supp.B.17, Supp.B.18 for dimensions 0 to 3 respectively. Since there are many cycles born at the filtration steps, later than for cycles shared between dataset, they result from connections not traced in the *COBRE* dataset.

Table 6: Kruskal-Wallis test results for comparison of vertical and horizontal coordinates of centroids of persistence landscapes.

| | Dimension 0 | | Dimension 1 | | Dimension 2 | |
|---|---|---|---|---|---|---|
| | x | y | x | y | x | **y** |
| *p-value* | **<0.01** | **<0.01** | **<0.01** | **<0.01** | **<0.01** | $= 0.102$ |
| *Number of Observations* | [176, 433] | [176, 433] | [1004, 314] | [1004, 314] | [1186, 66] | [1186, 66] |
| $\chi^2$*-statistic* | 12.8337 | 32.7345 | 107.536 | 47.0329 | 69.5192 | 2.67231 |
| *Rank Sums* | [60731, 125014] | [6494, 120804] | $[7.23182 \times 10^5, 1.4604 \times 10^5]$ | $[7.02508 \times 10^5, 1.66712 \times 10^5]$ | [766866, 17512] | $[7.4770 \times 10^5, 36675]$ |
| *Degrees of Freedom* | 1 | 1 | 1 | 1 | 1 | 1 |

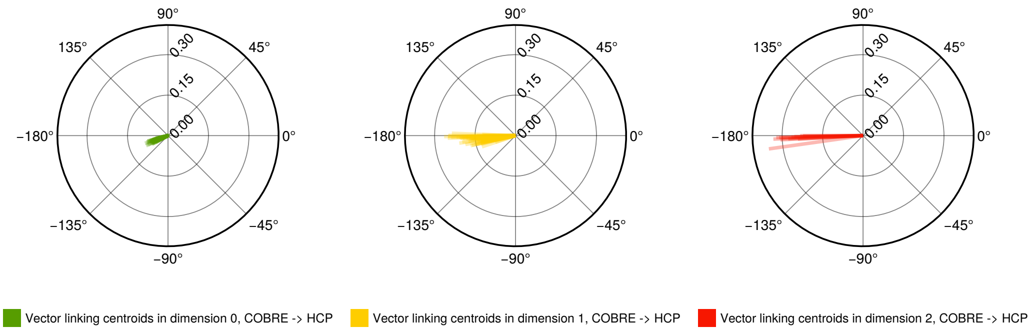


Figure Supp.B.14: Comparison of centroids of shared persistent landscapes for cycles in dimensions 0, 1, 2. Shared cycles are cycles that are found in at least one particiapnt in *COBRE* and in 1 participant in *HCP*. Lines are vectors drawn from the *COBRE* centroid to a centroid in a corresponding *HCP* cycle. There were 292, 116, 12 shared cycles in dimensions 0, 1, 2 respectively. The arrows are pointing down and left, and all the landscapes computed from *HCP* data are shifted to the left when compared with equivalent cycles from *COBRE* data.

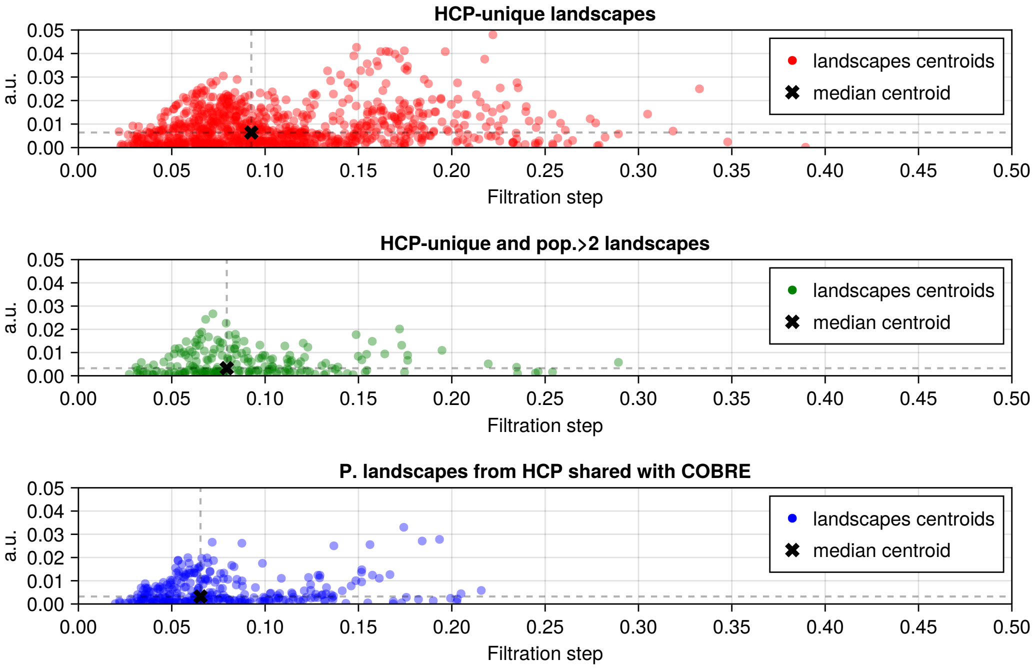


Figure Supp.B.15: Comparison of centroids of *HCP*, dimension 1 cycle not found in *COBRE*, with cycles centroids from *HCP* that are shared with *COBRE*. In all plots, the cross was used to indicate the median of the centroids. The vertical axis represents the landscape's height.

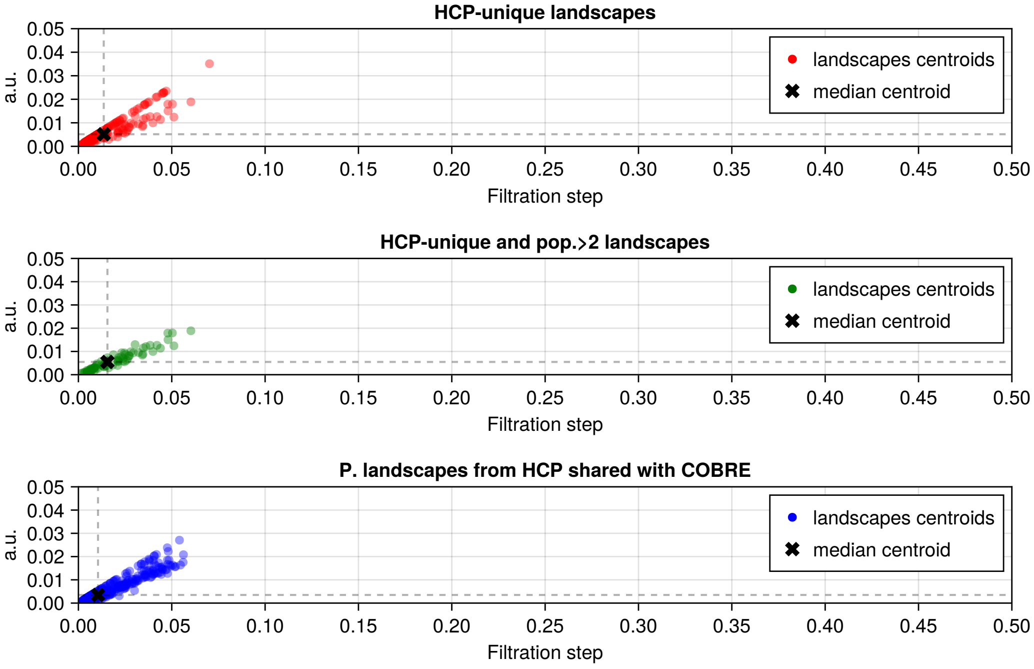


Figure Supp.B.16: Comparison of centroids of HCP, dimension 0 cycles not found in COBRE, with cycle sublandscape centroids from HCP that are shared with COBRE. In all plots, the cross was used to indicate the median of the centroids.

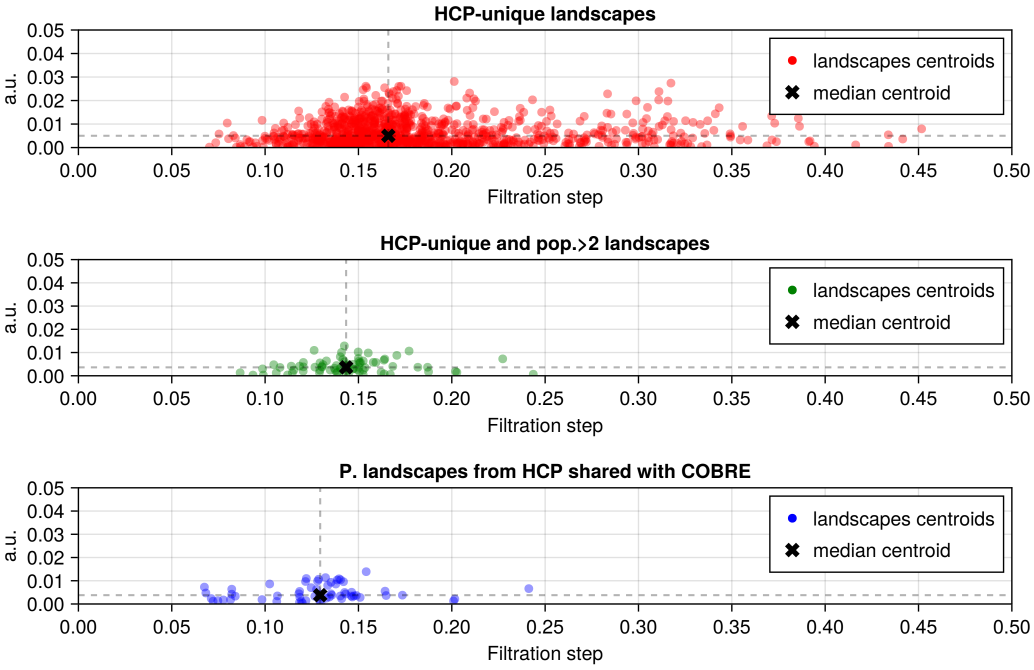


Figure Supp.B.17: Comparison of centroids of HCP, dimension 2 cycle not found in COBRE, with cycles centroids from HCP that are shared with COBRE. In all plots, the cross was used to indicate the median of the centroids.

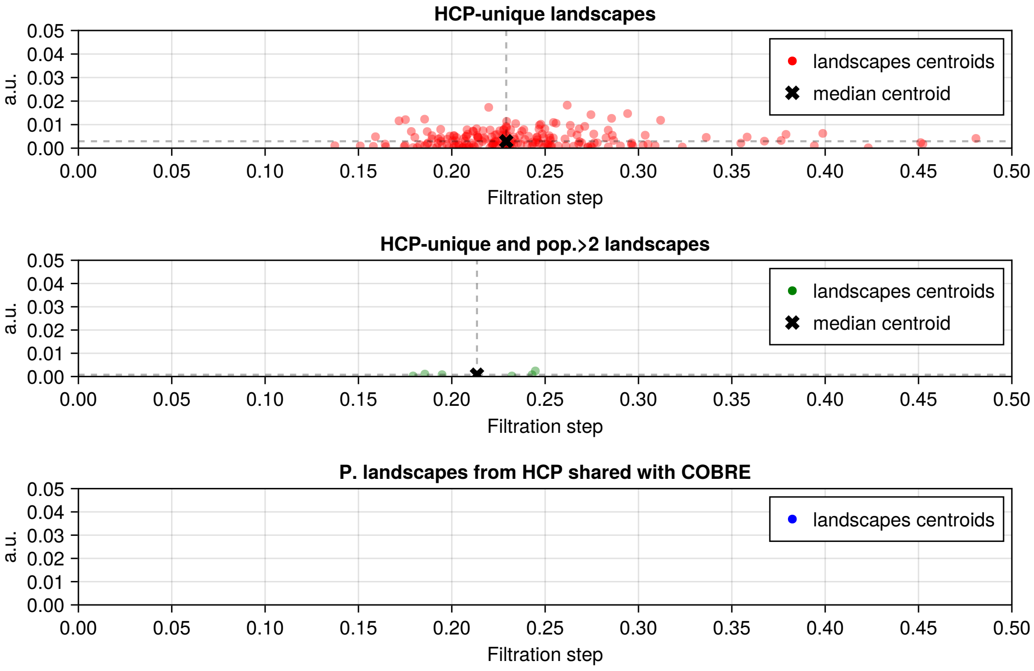


Figure Supp.B.18: Comparison of centroids of HCP, dimension 3 cycle not found in COBRE, with cycles centroids from HCP that are shared with COBRE. In all plots, the cross was used to indicate the median of the centroids.

#### B.7 Popularity significance tests

To assess the association between the presence of a cycle and group membership for each dimension ($D = 0, 1, 2$; note that $D = 3$ was omitted due to the lack of shared cycles), we conducted Fisher's exact test on a $2 \times 2$ contingency table for each cycle found in five or more subjects in the COBRE and HCP datasets. The null hypothesis is that there is no difference in the presence of the cycle between the two groups.

Table 7: Results of Fisher's exact tests for different dimensions for cycles found in 5 or more subjects of the COBRE and HCP datasets. Parameter $\alpha = 0.05$.

| Dimension | #Cycles | #($p < \alpha$) | #Bonferroni-corrected |
|---|---|---|---|
| 0 | 396 | 187 | 77 |
| 1 | 308 | 160 | 64 |
| 2 | 24 | 11 | 1 |

The p-values obtained from individual Fisher's exact tests were adjusted using the Bonferroni correction to account for multiple comparisons, with the significance threshold set at $\alpha = 0.05$. In total, 77, 64, and 1 cycle(s) were found to be statistically significant in dimensions $D = 0, 1, 2$, respectively.

##### B.7.1 Connectivity matrix comparison

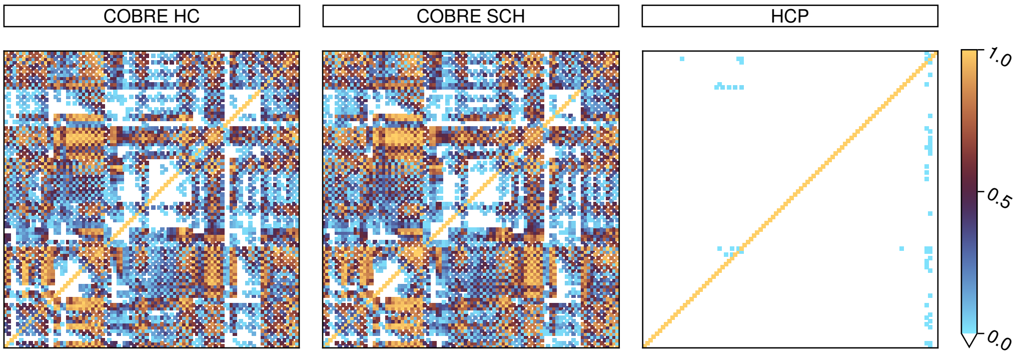


Figure Supp.B.19: Heatmaps of zeros counts in *COBRE* dataset (*HC* left, *SCH* middle) and *HCP* (right). Colours are normalised to maximal number of subjects, that is 44 for *HC* and *SCH* matrices, 88 for *HCP*. There are, on average, $2595.93 \pm 501.76$ zeros in a *COBRE* matrix and $82.01 \pm 416.1$ zeros in *HCP* matrix.

#### B.8 Detailed analysis of null model comparisons

The null models considered here were used to explore the nature of cycles found in the data, that is, they were used to (1) investigate the type of structural connectivity observed with anatomical data and (2) to investigate the importance of relative connections within patients.

We compare the numbers of cycles obtained from null model simulated datasets comprising of 88 matrices to those obtained from *COBRE*. As shown in Figure Supp.B.20, the various null models exhibit very different distributions of cycle type. In particular, they possess far fewer popular dimension 1 and 2 cycles. A cycle is denoted 'popular' if it occurs three or more times in the population of cycles.

Aggregating unique cycles from geometric null models (for unit cubes in $\mathbb{R}^n$ for $n = \{3, 4, 5, 10\}$) did not give any repetition among the cycles in dimensions 1 and above. Also, the total number of cycles in dimensions 0 and 1 is larger than for other NM (except for Random NM). For dimensions 2 and 3, the total number of cycles is matched with the *COBRE* and *HCP* datasets for $n = 5$ (however, the total number of cycles for this model in dimensions 0 and 1 remain much larger than in the datasets).

However, there is a lot of variability in how the samples are generated in the Geometric Null model- any two samples from this model do not have any correlation in the position of the nodes, while the brain is not a randomly connected structure, and it is characterised by small-world topology and conservation of wiring costs [Bullmore and Sporns, 2009, Oldham et al., 2022].

Next, a constrained model was tested, where the weights are larger for nodes that lie closer together, the 'Euclidean-brain' NM. The 'Euclidean-brain' NM with the least level of noise added is the most conservative case, simulating minimal variability among the samples. Its structure has 152, 58, 23 and 0 total unique cycles across dimensions 0 to 3, respectively. As expected, due to the low variability, the majority of the cycles in dimensions 0 and 1 are found in more than half of the generated matrices (90 and 40 cycles, respectively).

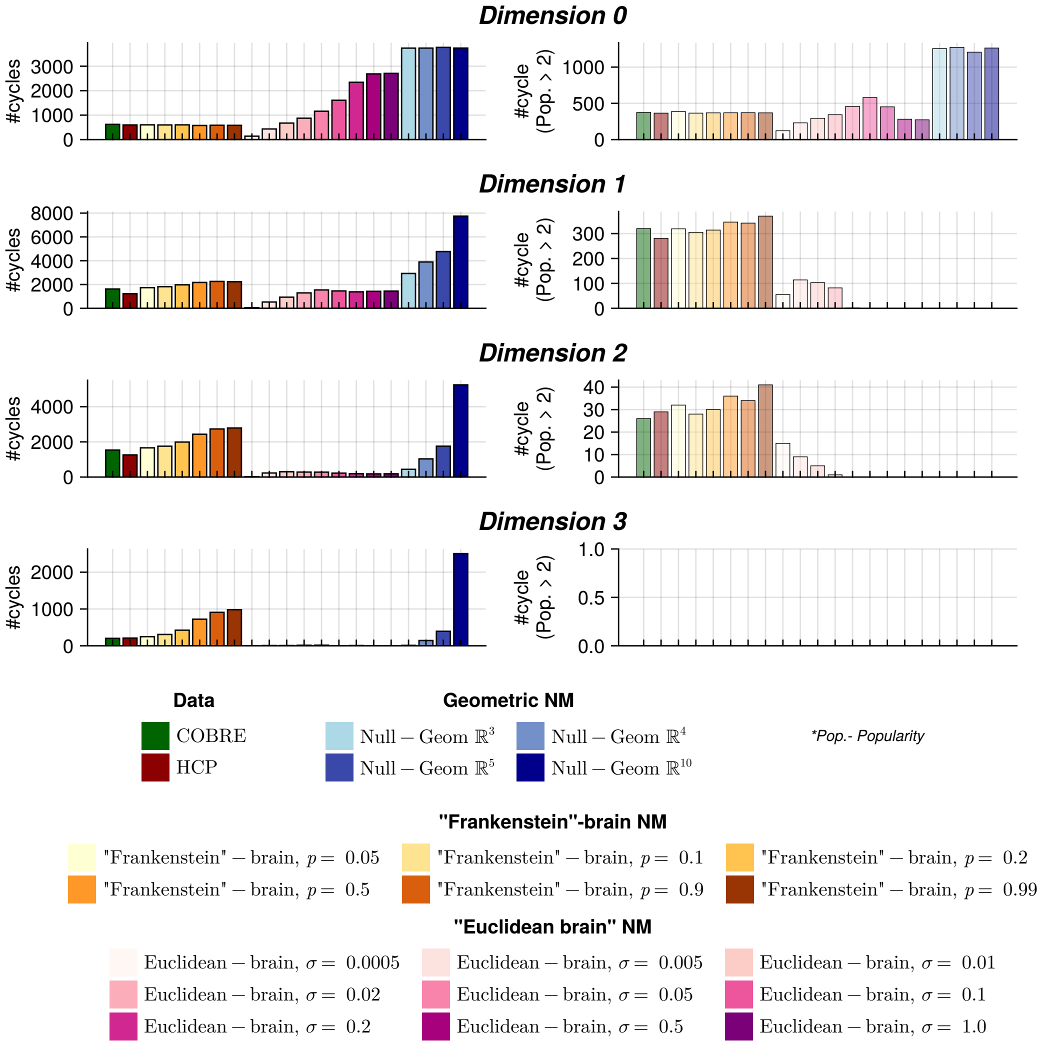


Figure Supp.B.20: Comparison of cycles' popularity. Data is shown for dimensions $d = 0, 1, 2, 3$, from top to bottom. Plots in the left column show the total count of unique cycles found in each dataset (Random NM is not included, as the total number of cycles is by an order of magnitude larger than for Geometric NM). Plots in the right column show the total count of unique cycles that were found in more than 2 subjects in each dataset. Different colours of the bars are used for different data, as shown in the legends below the plots. All datasets contained 88 connectivity structures. The parameter $\sigma$ in 'Euclidean-brain' NM is the standard deviation of the Gaussian noise. The parameter $p$ in 'Frankenstein-brain' NM is the probability of swapping an edge among the matrices. Please see section 4.4 for more details about the models.

The raster plots showing the structure of the cycles are presented in Supp.B.27, visualisations of examples of node distributions in "brain space" is presented in Supplementary Figures Supp.B.30. Increasing the noise level increases the total number of unique cycles across all dimensions (shown in the left column of the Figure Supp.B.20) at the expense of decreases in the popularity of the cycles (right column in the same plot). Interestingly, with the noise level that yields the same number of cycles in dimension 1 as for *COBRE* data ($\sigma = 0.05$), there are almost no popular cycles. Another difference between NM results and the *COBRE* dataset results were that there were almost no cycles (in dimension 2) or none at all (in dimension 3) found across all noise levels.

The 'Euclidean-brain' NM shows that by gradually increasing the noise, the total number of cycles increases, reaching the level found in the datasets and producing popular cycles. This may suggest that the brain may have some underlying structure from which every individual diverges, thus producing a large population of cycles, with some of them found across multiple subjects.

In the 'Euclidean-brain' NM, the distance-based wiring ensures that there are some groups of regions that are forming cycles repeatedly earlier than others, i.e., in Supplementary Figure Supp.B.27 the cycle indexed 1 is formed before cycle indexed 2 and after cycle index 44. We hypothesise that this kind of global relation (i.e. if a connection between regions "A" and "B" are in some relation to connections between any other pair of regions) is also present in the data, and this is explored with the most conservative 'Frankenstein-brain' NM, where the weights are shuffled only within the connections.

In the 'Frankenstein-brain' NM, the total number of cycles in dimensions 1 to 3 increased with the increase of the probability of swapping the connections; however, the total number of popular cycles remained similar (for dimensions 0 and 1) or increased (for dimension 2), showing that this model reproduces the repeatability of the cycles, even after completely randomising connections with the highest probability of swapping connections. This suggests that there exists a global relation of connections across individuals, which can be revealed with PH.

From the 'Euclidean-brain' model study, we see that the cycles found in multiple subjects in the data are due to the underlying connectivity pattern, from which individual brains can diverge due to their unique connectivity. In the 'Euclidean-brain' model, with almost no noise added, far fewer cycles are found that are present in every 'subject', and with the increase of noise added, more cycles are found with less popularity, to a point where all cycles are singletons.

In contrast, we found that the connectivity structure found in humans is unique in the way that it produces many more high-dimensional structures than the 'Euclidean-brain' model (high-dimensional structures were also reported in the study by Sizemore et al. (2018) [Sizemore et al., 2018]).

In the 'Frankenstein-brain' NM, which is closest to that of real data, it was shown that almost as many popular cycles are produced, even though 'Frankenstein-brain' matrices were created by taking original data matrices and randomly swapping a proportion of their entries with those from other data matrices. This suggests that some cycles are present in very much the same way across the whole population, and give rise to universal structures. Across subjects, these nodes join the filtration at approximately the same step, forming the same cycle.

For example, the four most popular cycles in dimension 1 are found in at least 85% of COBRE subjects, but they are also found in all of the instances of 'Frankenstein-brain' NM. The regions that constitute these cycles are combinations of regions like *precuneus*, *anterior and posterior cingulate cortex*, *superior frontal cortex*, which are regarded as "brain hubs" [Bullmore and Sporns, 2009, Van Den Heuvel and Sporns, 2013], that is brain regions characterised by high centrality (details about cycles can be found in Supplementary Tables 39, 40, 41). Three more regions, *insular cortex*, *temporal cortex*, and *lateral parietal cortex*, are brain hubs that occur in other cycles. Therefore, our study indicates that hub nodes are closely interconnected with each other, forming hub-cycles that persist across all permutation models.

The presence of singletons (cycles found in only a single participant), especially if born later, may be due to noise rather than a result of individual variation. Indeed, the mean birth time for singeltons for both *COBRE* and *HCP* is significantly larger than for any cycle that is found in more than one subject (Mann-Whitney U test of the distribution of birth time for singleton cycles and birth time of non-singleton cycles, $p \ll 1$ for both datasets, details presented in the Table 8).

Table 8: Mean birth time of singletons versus non-singletons for *COBRE* and *HCP* datasets. Values are given as mean ± standard deviation.

| Dataset | Singletons | Non-singletons | $p$ | # observations [singeltons vs. not] | Mann-Whitney-U statistic |
|---|---|---|---|---|---|
| *COBRE* | $0.1256 \pm 0.0718$ | $0.0632 \pm 0.0450$ | $p \ll 0.001$ | $[2350, 3084]$ | 1 410 880 |
| *HCP* | $0.0825 \pm 0.0570$ | $0.0336 \pm 0.0330$ | $p \ll 0.001$ | $[1965, 2627]$ | 1 056 440 |

##### B.8.1 Null model reverse filtrations

For *COBRE*, the total number of cycles in the reverse filtration is an order of magnitude larger than for *HCP* or Random NM. This increase relates directly to the increased sparsity of the *COBRE* connectivity matrices- there is a large connected component at the beginning of filtration for each of the ordered matrices, which creates a basis for high-dimensional structures. Another difference is that there are more popular cycles for the reverse filtration of the data than for the Random NM.

Table 9: Total number of cycles for forward and reverse filtration (random matrix added for comparison)

| Dimension | | *COBRE* | rev. *COBRE* | *HCP* | rev. *HCP* | Random NM |
|---|---|---|---|---|---|---|
| 0 | Total Cycles | 622 | 93 | 599 | 780 | 2678 |
| | Popularity $> 2$ | 374 | 18 | 365 | 449 | 288 |
| 1 | Total Cycles | 1625 | 6262 | 1227 | 9515 | 13639 |
| | Popularity $> 2$ | 320 | 435 | 281 | 1984 | 0 |
| 2 | Total Cycles | 1536 | 82118 | 1260 | 31189 | 35412 |
| | Popularity $> 2$ | 26 | 1590 | 29 | 2690 | 0 |
| 3 | Total Cycles | 202 | 323082 | 210 | 45717 | 73624 |
| | Popularity $> 2$ | 0 | 2250 | 0 | 1202 | 0 |

##### B.8.2 Comparison of cycle structure, popularity, and topological properties between COBRE and HCP for dimensions 2 and 3

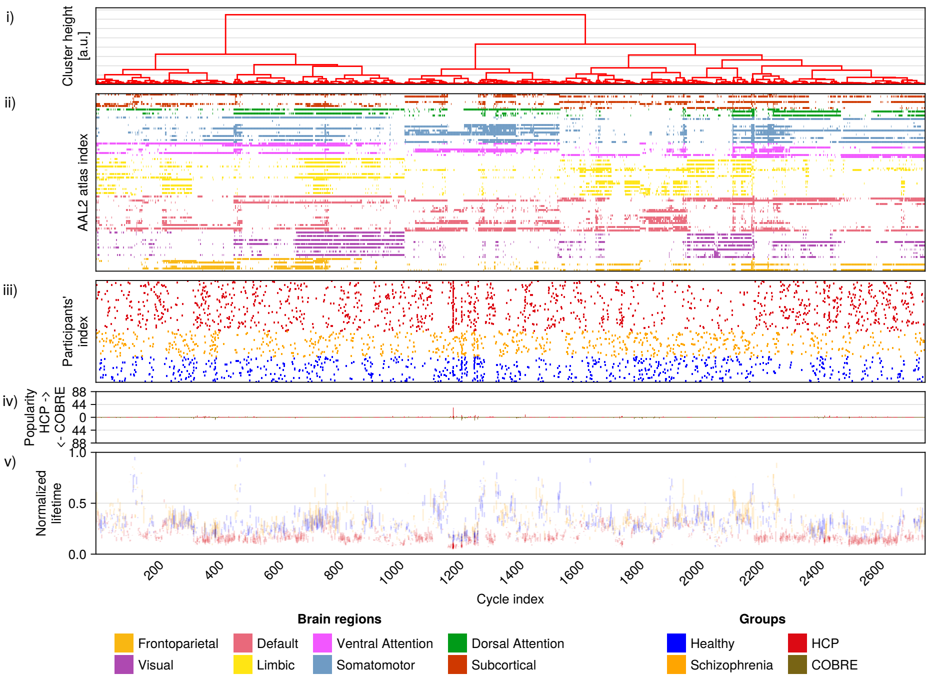


(a) Dimension 2

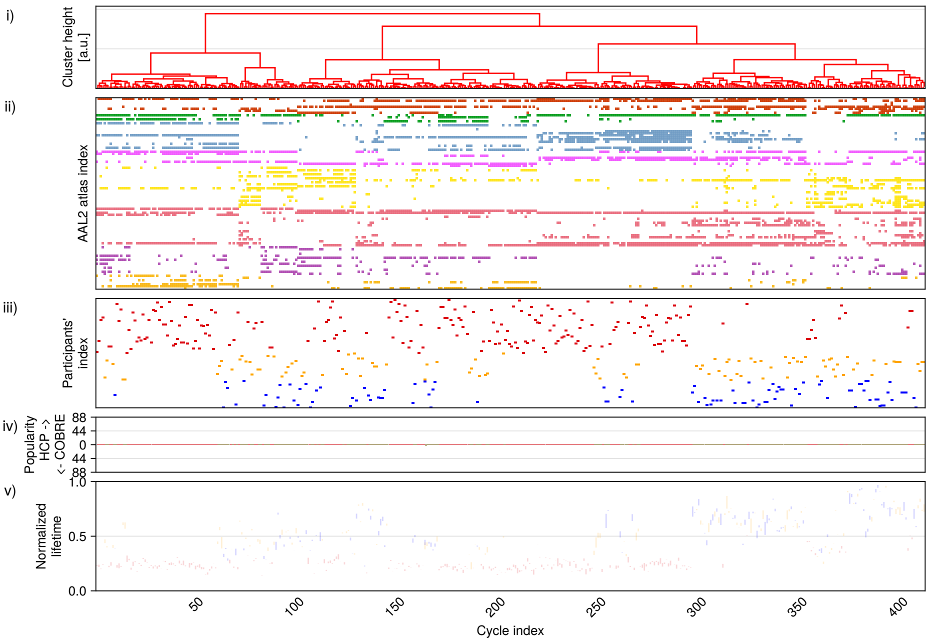


(b) Dimension 3

Figure Supp.B.21: Unique cycles structures and popularity presented as a raster plot for dimensions $d = 2, 3$ for Human Connectome Project (HCP) (red) and COBRE (orange for SCH and blue for HC). For each figure, the plots are structured as described in Figure 5.

#### B.9 Betti curves for forward and reverse for data and null models

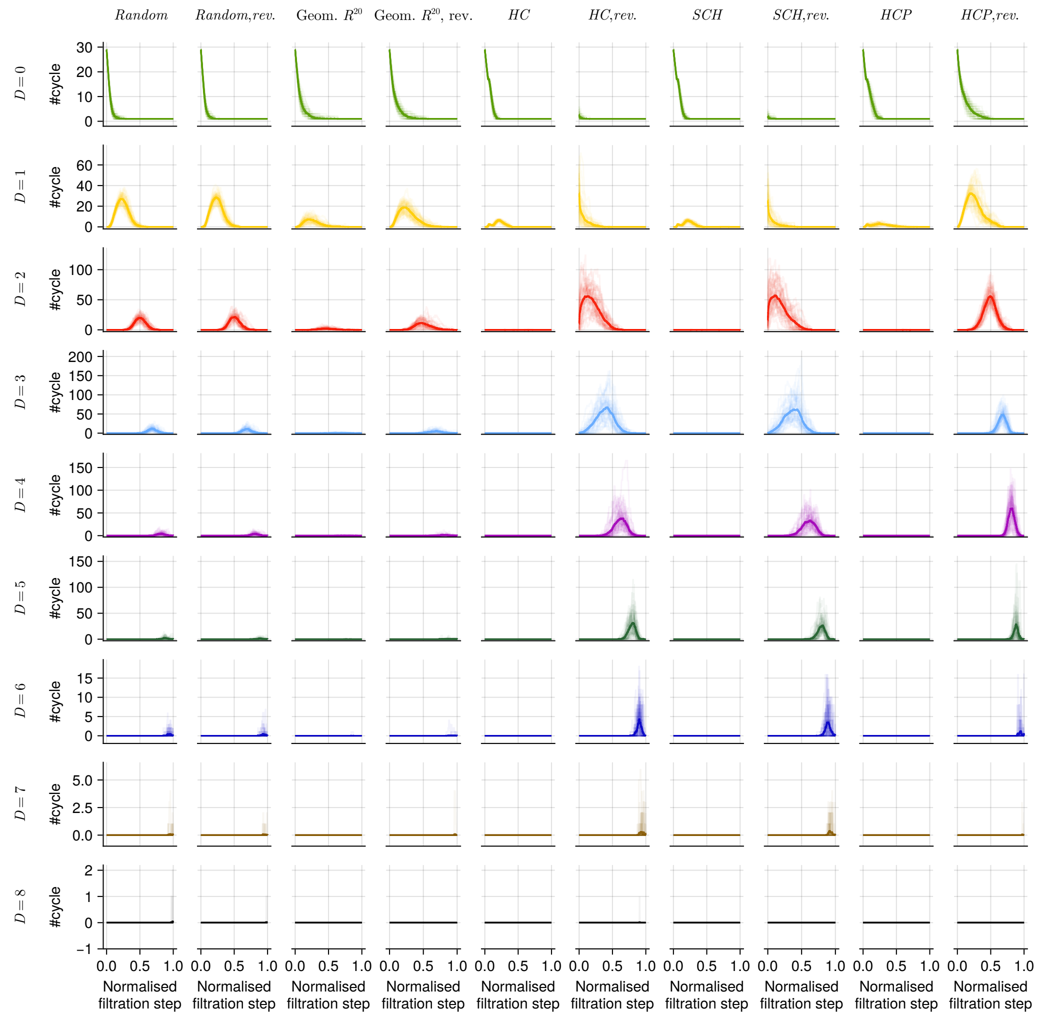


Figure Supp.B.22: Comparison of $\beta$ curves for Random NM, Geometric $R^{20}$ NM, *COBRE*, *HCP* and their reverse filtration for dimension in range $0 \leq D \leq 8, D \in \mathbb{Z}$. Due to computational intensity, matrix size was limited to 30, i.e. for *COBRE* and *HCP* only the first 30 brain regions from AAL2 atlas were used for the computations. We investigated cycles for dimension up to $D = 15$ (no cycles were found for in $D = 8, \ldots, 15$).

##### B.9.1 Shuffling weak connections.

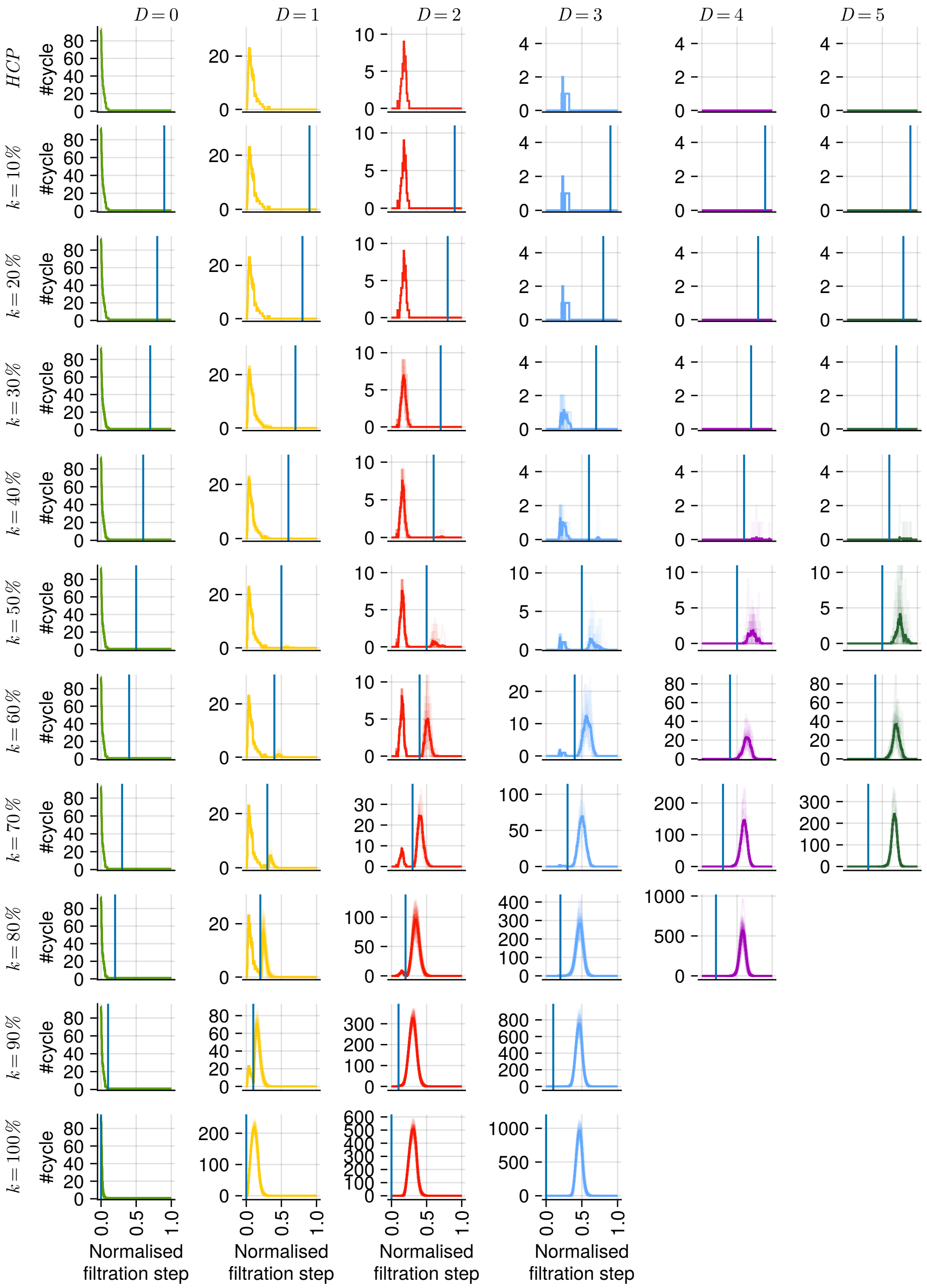


Figure Supp.B.23: Tsunami of randomness. Effect of shuffling of weak connections in *HCP* on the Betti curves. Solid lines are the average Betti curves; faint colours were used for individual Betti curves. All computations were done for $94 \times 94$ matrices. In the first row, Betti curves are shown for the forward filtration from the ordered connectivity matrices for a selected HCP subject. For the following rows, Betti curves were computed for the forward filtration of the connectivity matrix of the same subject, in which $k$ weakest connections (indicated by a blue line in each plot) were shuffled 44 times. Results are shown for the dimensions in the range $0 \leq D \leq 5, D \in \mathbb{Z}$ for $k \leq 70\%$ (missing figures for $D \geq 3$ and $k \geq 80\%$ are due to reaching available computational limits).

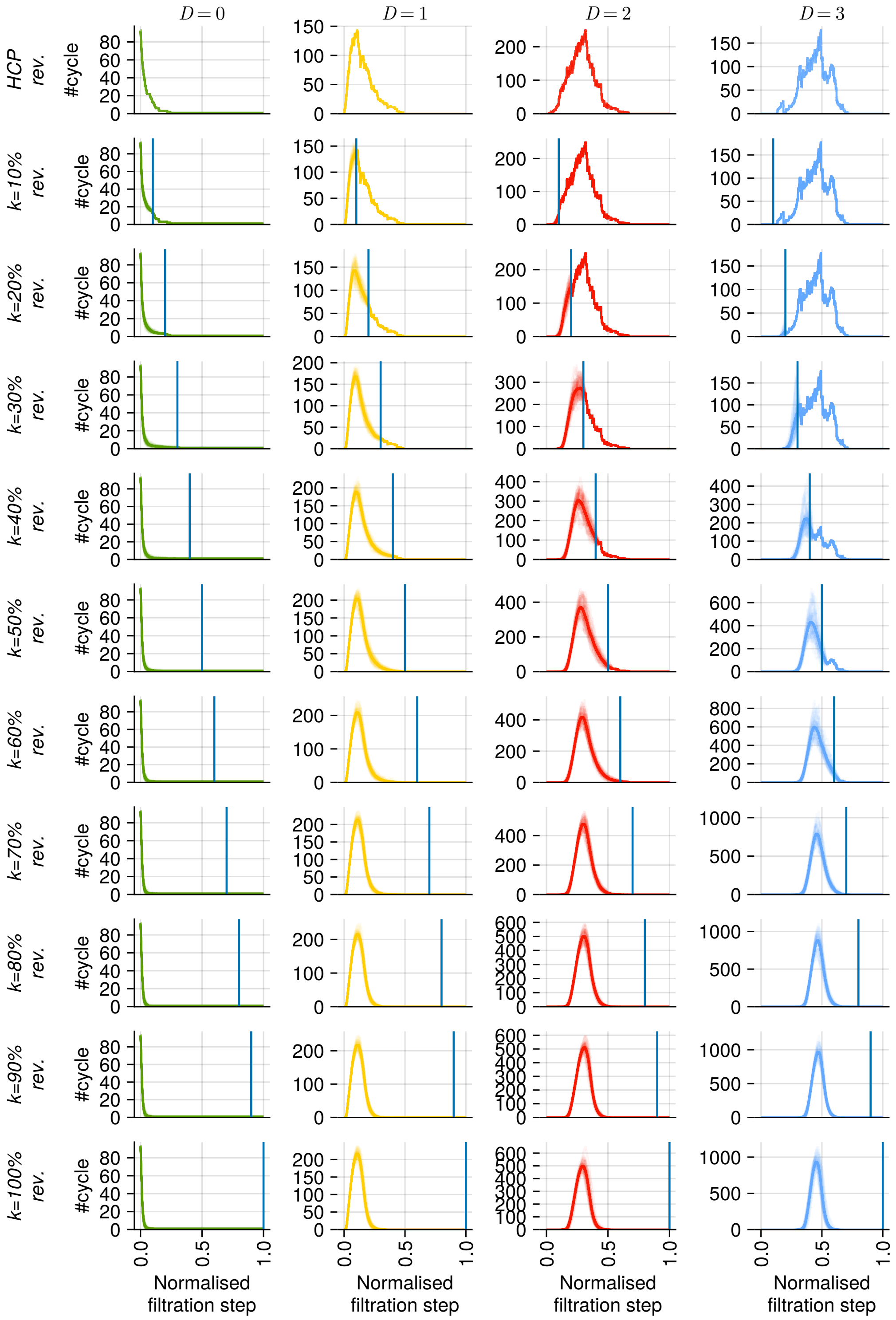


Figure Supp.B.24: Tsunami of randomness. Effect of shuffling of weak connections in the reverse filtration of *HCP* on the Betti curves. Solid lines are the average Betti curves, a faint colour was used for individual Betti curves. All computations were done for matrix size 94. In the first row, Betti curves are shown for the reverse filtration from the ordered connectivity matrices for a selected HCP subject. For the following rows, Betti curves were computed for the reverse filtation of the connectivity matrix of the same subject, in which $k$ weakest connections (indicated by a blue line in each plot) were shuffled 44 times. Results are shown for the dimensions in the range $0 \leq D \leq 3, D \in \mathbb{Z}$.

##### B.9.2 Cycle structure, popularity, and topological properties for null models

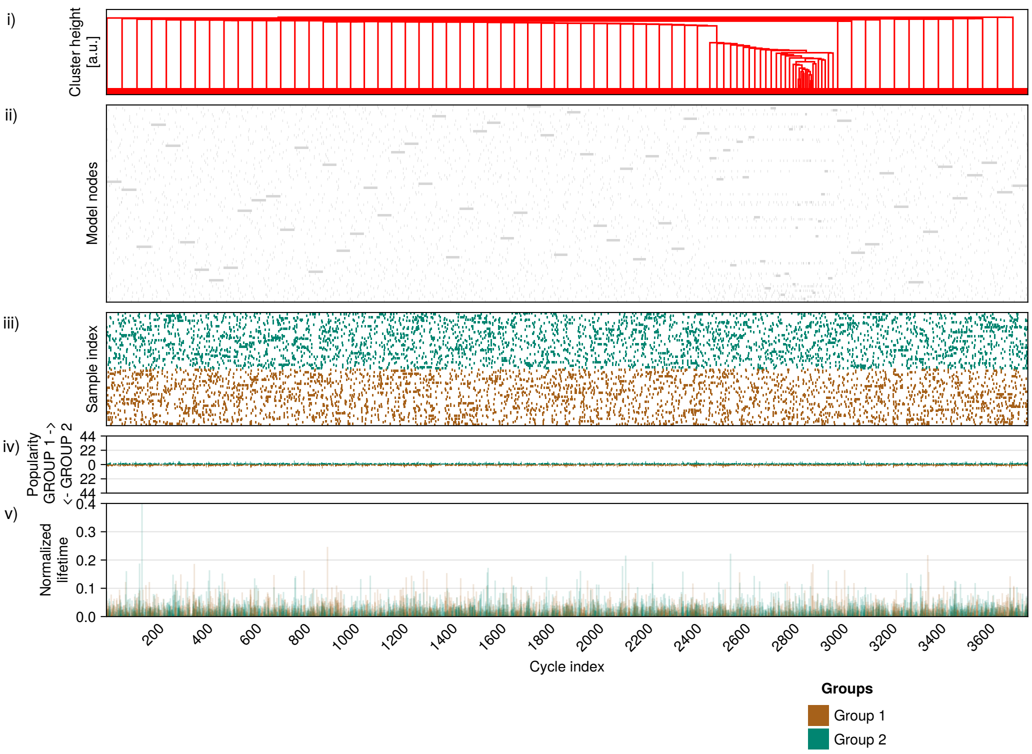


(a) Dimension 0

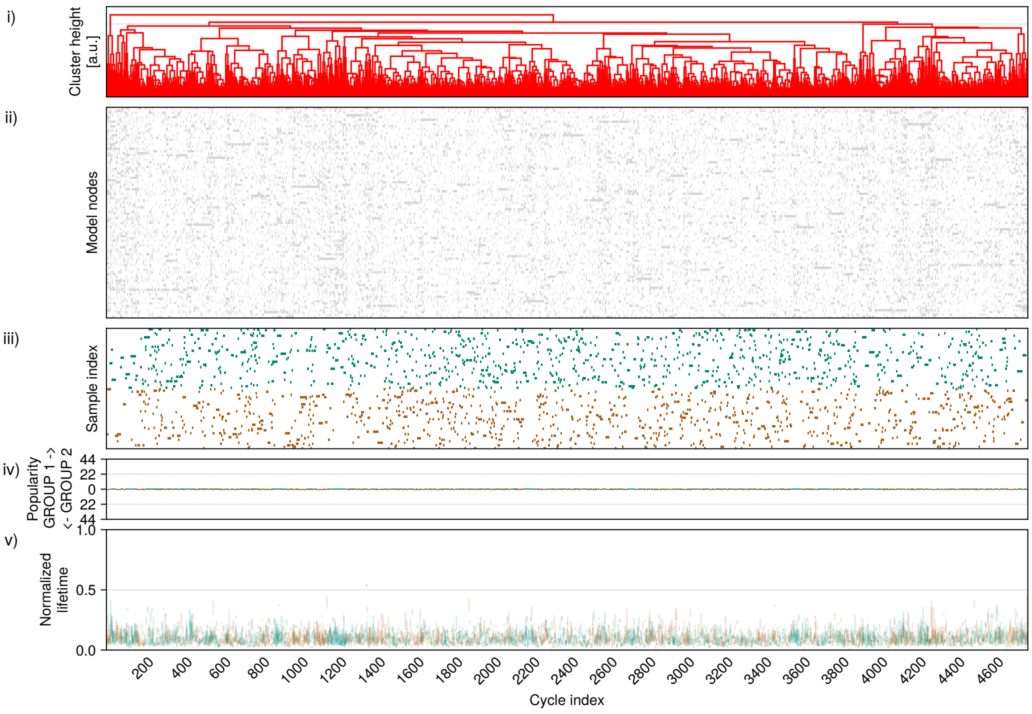


(b) Dimension 1

Figure Supp.B.25: Structure, popularity and topological properties of cycles from null geometric model $\mathbb{R}^5$. The model was used to generate 88 samples that were randomly split into 2 groups of the same size (44 samples in each group); colours used for sample allocation are shown in the legend.

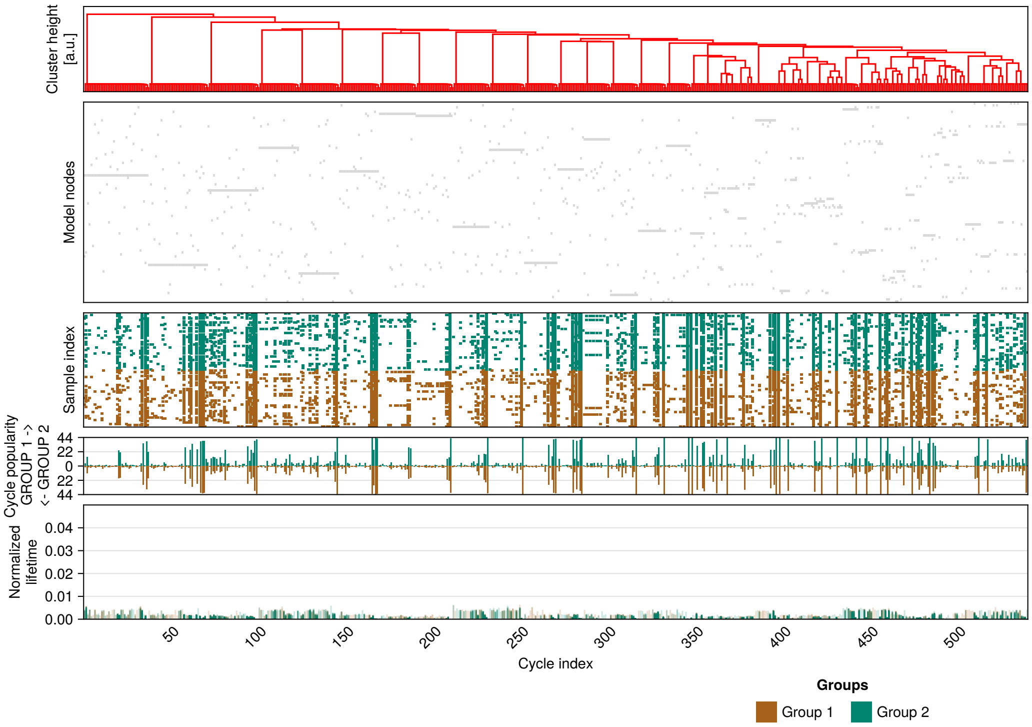


(a) Dimension 0

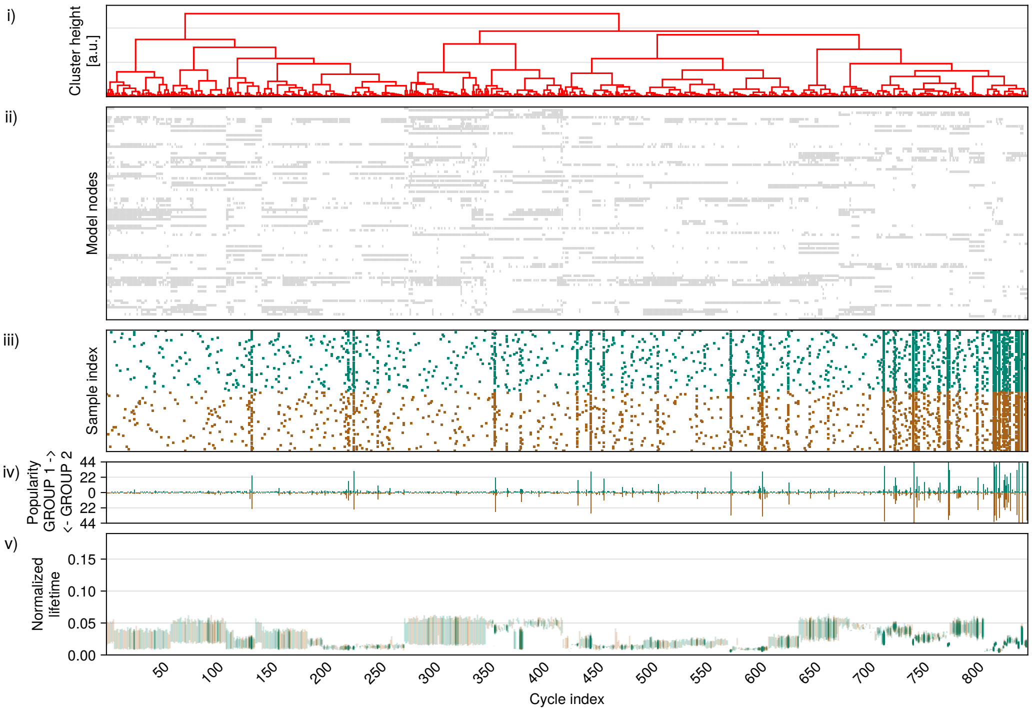


(b) Dimension 1

Figure Supp.B.26: Structure, popularity and topological properties of cycles for ‘Euclidean-brain’ null model, standard deviation of Gaussian distribution $\sigma = 0.005$.

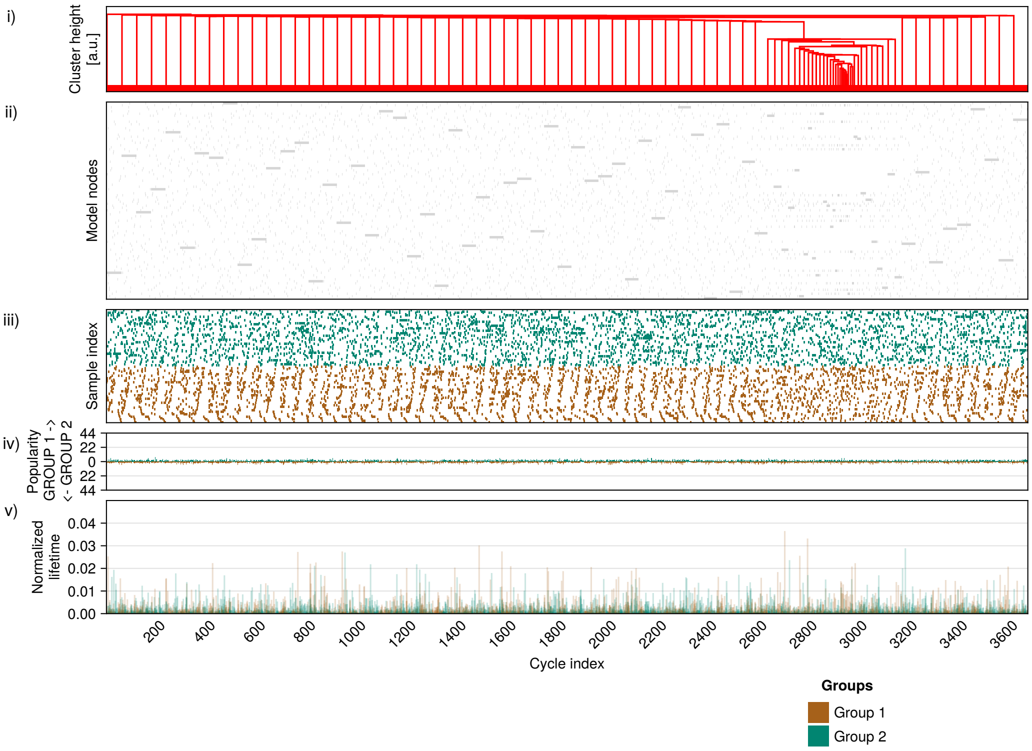


(a) Dimension 0

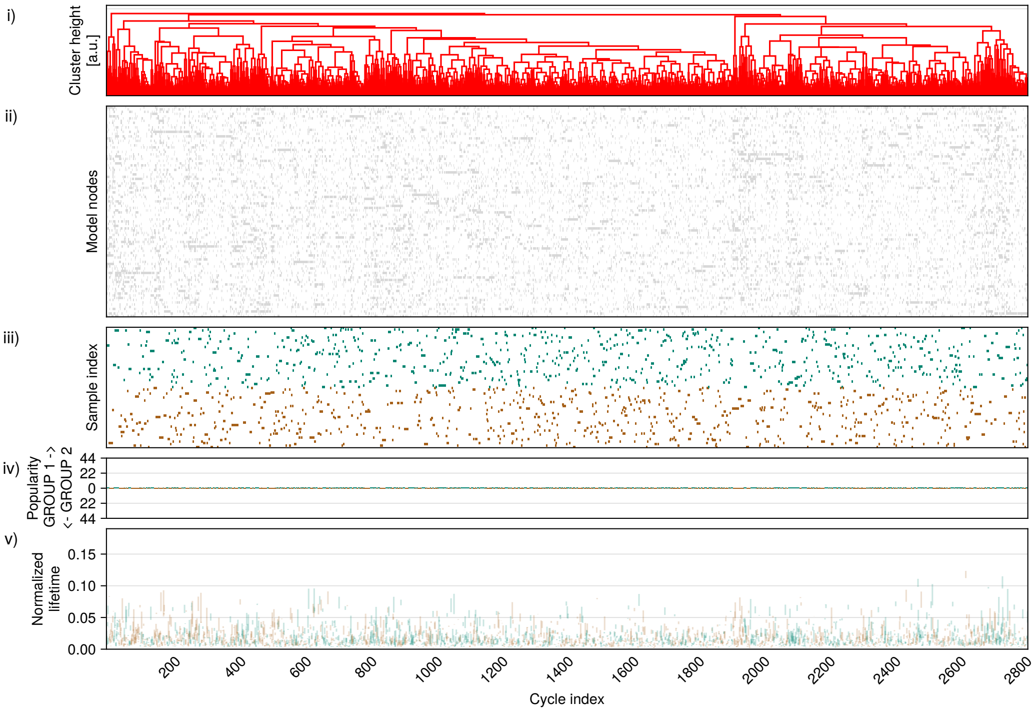


(b) Dimension 1

Figure Supp.B.27: Structure, popularity and topological properties of cycles for 'Euclidean-brain' null model, standard deviation of Gaussian distribution $\sigma = 0.5$.

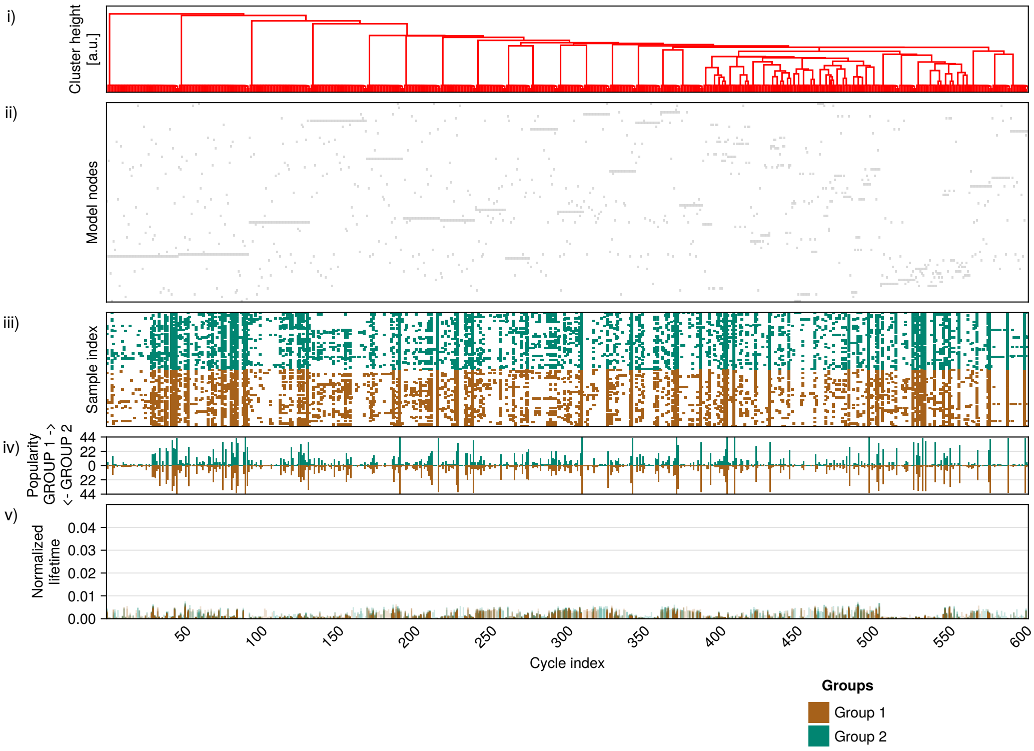


(a) Dimension 0

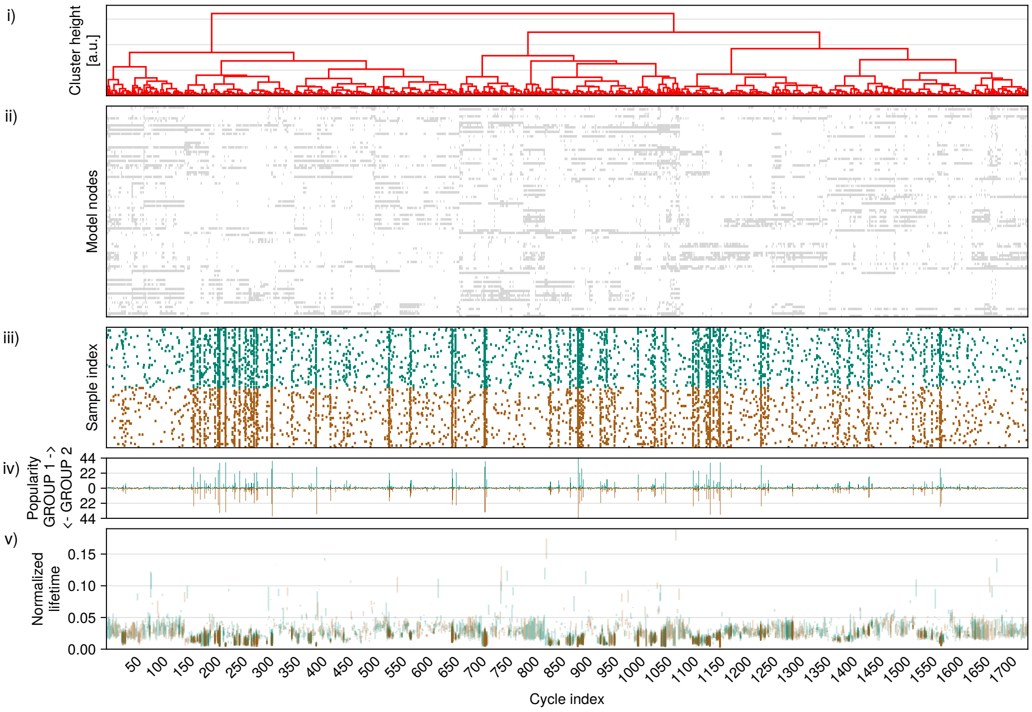


(b) Dimension 1

Figure Supp.B.28: Structure, popularity and topological properties of cycles for ‘Frankenstein-brain’ Null Model, probability of swapping connection $p = 0.05$.

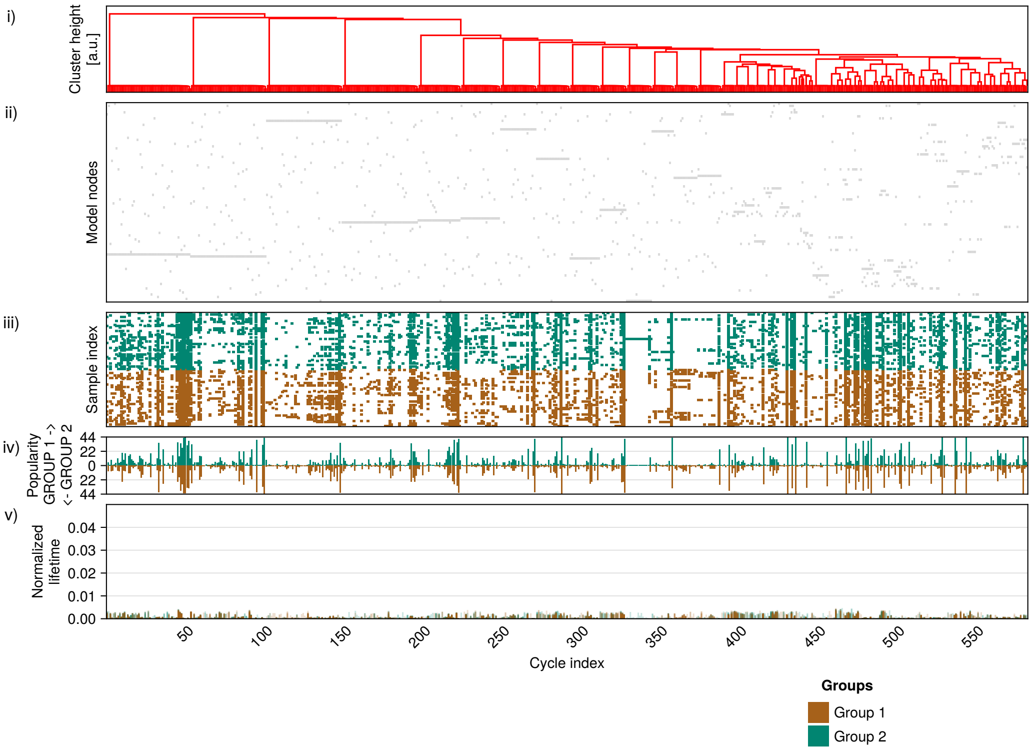


(a) Dimension 0

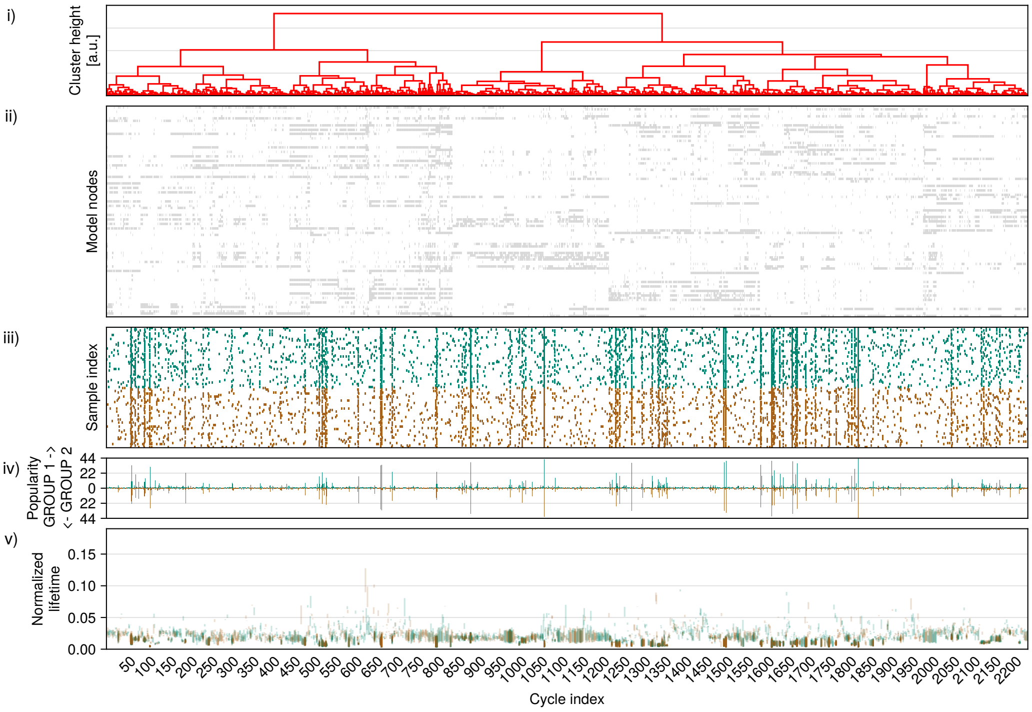


(b) Dimension 1

Figure Supp.B.29: Structure, popularity and topological properties of cycles for 'Frankenstein-brain' Null Model, probability of swapping connection $p = 0.99$.

##### B.9.3 Visualisation of nodes in 'Euclidean-brain' null model

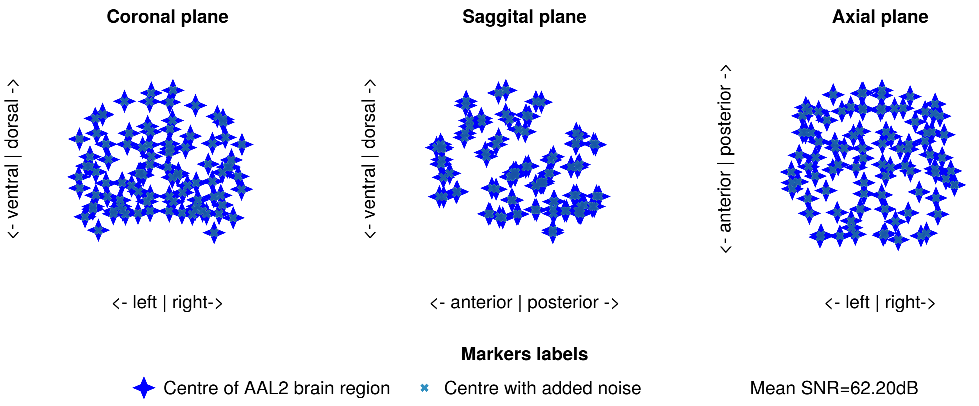


(a) Noise standard deviation $\sigma = 0.0005$, $SNR = 60.20dB$

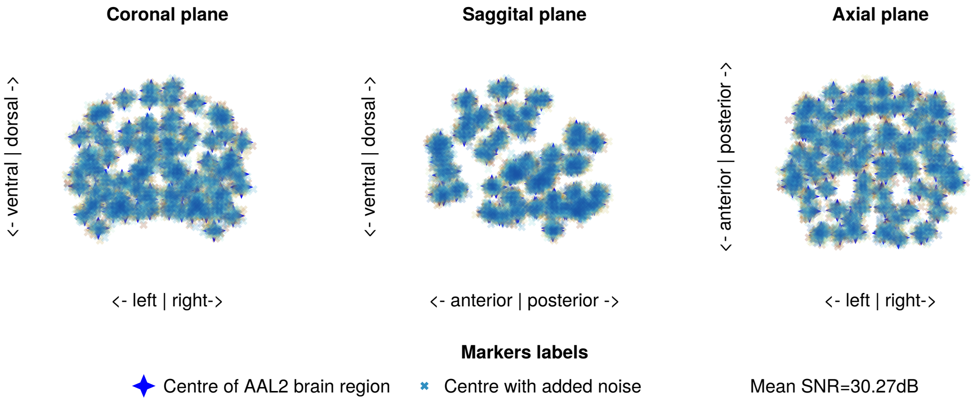


(b) Noise standard deviation $\sigma = 0.02$, $SNR = 30.27dB$

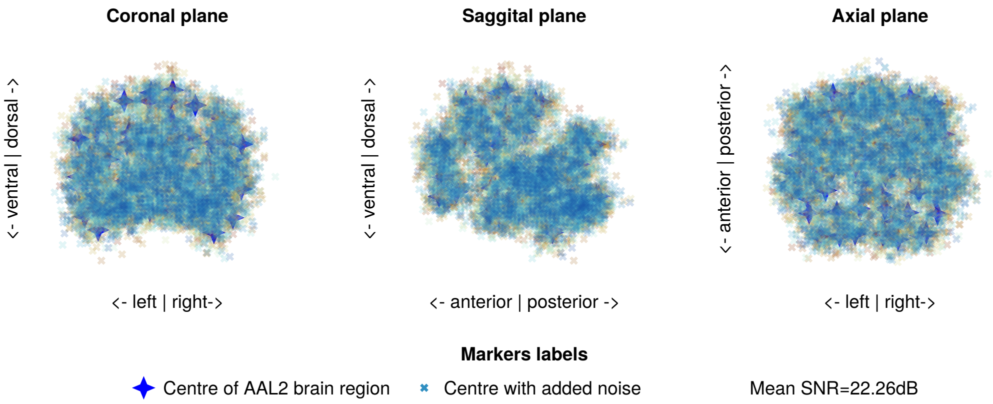


(c) Noise standard deviation $\sigma = 0.05$, $SNR = 22.26dB$

Figure Supp.B.30: Distribution of points with added noise for brain-euclidean null model.

#### B.10 PANSS score interpretation

The individual scores for the PNASS scale are shown in the Figure Supp.B.31.

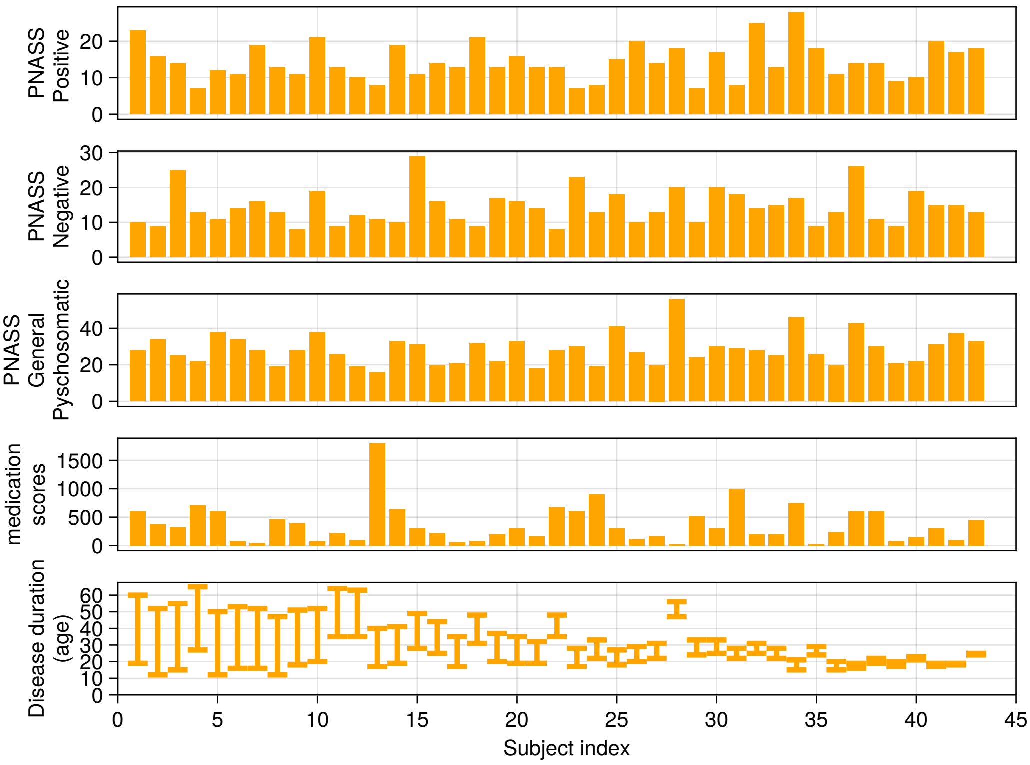


Figure Supp.B.31: participant evaluation with PNASS. The scores are split into 3 types of symptoms- PNASS positive (1st row), PNASS negative (2nd row), general psychosomatic (3rd row). Additionally, the individual medication scores (4th row) and disease duration and onset (5th row) are shown at the bottom of the figure. All results are sorted by the duration of the disease.

#### B.11 Brain regions in popular cycles

##### B.11.1 Significant popular dimension 0 cycles

Table 10: Cycle 68: 20 total schizophrenia subjects, 21 total healthy controls. ID is AAL2 atlas region index.

| | ID | Region | Side | Yeo network |
|---|---|---|---|---|
| 1 | 40 | Posterior cingulate gyrus | R | Default |
| 2 | 71 | Precuneus | L | Default |

Table 11: Cycle 101: 15 total schizophrenia subjects, 11 total healthy controls. ID is AAL2 atlas region index.

| | ID | Region | Side | Yeo network |
|---|---|---|---|---|
| 1 | 61 | Postcentral gyrus | L | Somatomotor |
| 2 | 65 | Inferior parietal gyrus | L | Dorsal Attention |

Table 12: Cycle 107: 10 total schizophrenia subjects, 13 total healthy controls. ID is AAL2 atlas region index.

| | ID | Region | Side | Yeo network |
|---|---|---|---|---|
| 1 | 62 | Postcentral gyrus | R | Somatomotor |
| 2 | 74 | Paracentral lobule | R | Somatomotor |

Table 13: Cycle 108: 11 total schizophrenia subjects, 12 total healthy controls. ID is AAL2 atlas region index.

| | ID | Region | Side | Yeo network |
|---|---|---|---|---|
| 1 | 34 | Insula | R | Ventral Attention |
| 2 | 78 | Lenticular nucleus, Putamen | R | Ventral Attention |

Table 14: Cycle 111: 12 total schizophrenia subjects, 10 total healthy controls. ID is AAL2 atlas region index.

| | ID | Region | Side | Yeo network |
|---|---|---|---|---|
| 1 | 61 | Postcentral gyrus | L | Somatomotor |
| 2 | 71 | Precuneus | L | Default |

Table 15: Cycle 113: 10 total schizophrenia subjects, 12 total healthy controls. ID is AAL2 atlas region index.

| | ID | Region | Side | Yeo network |
|---|---|---|---|---|
| 1 | 39 | Posterior cingulate gyrus | L | Default |
| 2 | 72 | Precuneus | R | Dorsal Attention |

Table 16: Cycle 115: 11 total schizophrenia subjects, 11 total healthy controls. ID is AAL2 atlas region index.

| | ID | Region | Side | Yeo network |
|---|---|---|---|---|
| 1 | 75 | Caudate nucleus | L | Limbic |
| 2 | 76 | Caudate nucleus | R | Subcortical |

##### B.11.2 Significant popular dimension 1 cycles

Table 17: Cycle 16: 27 total schizophrenia subjects, 24 total healthy controls. ID is AAL2 atlas region index.

| | ID | Region | Side | Yeo network |
|---|---|---|---|---|
| 1 | 37 | Middle cingulate & paracingulate gyri | L | Ventral Attention |
| 2 | 38 | Middle cingulate & paracingulate gyri | R | Ventral Attention |
| 3 | 71 | Precuneus | L | Default |
| 4 | 72 | Precuneus | R | Dorsal Attention |

Table 18: Cycle 18: 23 total schizophrenia subjects, 24 total healthy controls. ID is AAL2 atlas region index.

| | ID | Region | Side | Yeo network |
|---|---|---|---|---|
| 1 | 49 | Cuneus | L | Visual |
| 2 | 50 | Cuneus | R | Visual |
| 3 | 71 | Precuneus | L | Default |
| 4 | 72 | Precuneus | R | Dorsal Attention |

Table 19: Cycle 20: 24 total schizophrenia subjects, 20 total healthy controls. ID is AAL2 atlas region index.

| | ID | Region | Side | Yeo network |
|---|---|---|---|---|
| 1 | 7 | Inferior frontal gyrus opercular part | L | Ventral Attention |
| 2 | 9 | Inferior frontal gyrus triangular part | L | Frontoparietal |
| 3 | 13 | Rolandic operculum | L | Somatomotor |
| 4 | 33 | Insula | L | Ventral Attention |

Table 20: Cycle 23: 24 total schizophrenia subjects, 19 total healthy controls. ID is AAL2 atlas region index.

| | ID | Region | Side | Yeo network |
|---|---|---|---|---|
| 1 | 47 | Calcarine fissure and surrounding cortex | L | Visual |
| 2 | 48 | Calcarine fissure and surrounding cortex | R | Visual |
| 3 | 49 | Cuneus | L | Visual |
| 4 | 50 | Cuneus | R | Visual |
| 5 | 71 | Precuneus | L | Default |
| 6 | 72 | Precuneus | R | Dorsal Attention |

Table 21: Cycle 24: 21 total schizophrenia subjects, 21 total healthy controls. ID is AAL2 atlas region index.

| | ID | Region | Side | Yeo network |
|---|---|---|---|---|
| 1 | 16 | Supplementary motor area | R | Somatomotor |
| 2 | 38 | Middle cingulate & paracingulate gyri | R | Ventral Attention |
| 3 | 62 | Postcentral gyrus | R | Somatomotor |
| 4 | 64 | Superior parietal gyrus | R | Dorsal Attention |
| 5 | 72 | Precuneus | R | Dorsal Attention |
| 6 | 74 | Paracentral lobule | R | Somatomotor |

Table 22: Cycle 26: 22 total schizophrenia subjects, 18 total healthy controls. ID is AAL2 atlas region index. A "mirror twin" to Cycle 42 (Table 68).

| | ID | Region | Side | Yeo network |
|---|---|---|---|---|
| 1 | 44 | Parahippocampal gyrus | R | Limbic |
| 2 | 60 | Fusiform gyrus | R | Visual |
| 3 | 88 | Temporal pole superior temporal gyrus | R | Limbic |
| 4 | 90 | Middle temporal gyrus | R | Default |
| 5 | 92 | Temporal pole middle temporal gyrus | R | Limbic |
| 6 | 94 | Inferior temporal gyrus | R | Limbic |

Table 23: Cycle 30: 19 total schizophrenia subjects, 17 total healthy controls. ID is AAL2 atlas region index.

| | ID | Region | Side | Yeo network |
|---|---|---|---|---|
| 1 | 1 | Precentral gyrus | L | Somatomotor |
| 2 | 3 | Superior frontal gyrus dorsolateral | L | Default |
| 3 | 4 | Superior frontal gyrus dorsolateral | R | Default |
| 4 | 5 | Middle frontal gyrus | L | Frontoparietal |
| 5 | 16 | Supplementary motor area | R | Somatomotor |
| 6 | 19 | Superior frontal gyrus medial | L | Default |
| 7 | 20 | Superior frontal gyrus medial | R | Default |
| 8 | 37 | Middle cingulate & paracingulate gyri | L | Ventral Attention |
| 9 | 38 | Middle cingulate & paracingulate gyri | R | Ventral Attention |
| 10 | 61 | Postcentral gyrus | L | Somatomotor |
| 11 | 63 | Superior parietal gyrus | L | Dorsal Attention |
| 12 | 65 | Inferior parietal gyrus | L | Dorsal Attention |
| 13 | 71 | Precuneus | L | Default |

Table 24: Cycle 32: 18 total schizophrenia subjects, 16 total healthy controls. ID is AAL2 atlas region index.

| | ID | Region | Side | Yeo network |
|---|---|---|---|---|
| 1 | 49 | Cuneus | L | Visual |
| 2 | 53 | Superior occipital gyrus | L | Visual |
| 3 | 55 | Middle occipital gyrus | L | Visual |
| 4 | 63 | Superior parietal gyrus | L | Dorsal Attention |
| 5 | 65 | Inferior parietal gyrus | L | Dorsal Attention |
| 6 | 67 | Supramarginal gyrus | L | Ventral Attention |
| 7 | 71 | Precuneus | L | Default |
| 8 | 85 | Superior temporal gyrus | L | Somatomotor |
| 9 | 89 | Middle temporal gyrus | L | Default |

Table 25: Cycle 33: 20 total schizophrenia subjects, 14 total healthy controls. ID is AAL2 atlas region index.

| | ID | Region | Side | Yeo network |
|---|---|---|---|---|
| 1 | 42 | Hippocampus | R | Limbic |
| 2 | 44 | Parahippocampal gyrus | R | Limbic |
| 3 | 48 | Calcarine fissure and surrounding cortex | R | Visual |
| 4 | 52 | Lingual gyrus | R | Visual |
| 5 | 60 | Fusiform gyrus | R | Visual |

Table 26: Cycle 35: 14 total schizophrenia subjects, 18 total healthy controls. ID is AAL2 atlas region index.

| | ID | Region | Side | Yeo network |
|---|---|---|---|---|
| 1 | 14 | Rolandic operculum | R | Somatomotor |
| 2 | 62 | Postcentral gyrus | R | Somatomotor |
| 3 | 68 | Supramarginal gyrus | R | Ventral Attention |
| 4 | 86 | Superior temporal gyrus | R | Somatomotor |

Table 27: Cycle 37: 17 total schizophrenia subjects, 14 total healthy controls. ID is AAL2 atlas region index. A "mirror twin" to Cycle 48 (Table 71).

| | ID | Region | Side | Yeo network |
|---|---|---|---|---|
| 1 | 56 | Middle occipital gyrus | R | Visual |
| 2 | 58 | Inferior occipital gyrus | R | Visual |
| 3 | 60 | Fusiform gyrus | R | Visual |
| 4 | 90 | Middle temporal gyrus | R | Default |
| 5 | 94 | Inferior temporal gyrus | R | Limbic |

Table 28: Cycle 40: 17 total schizophrenia subjects, 13 total healthy controls. ID is AAL2 atlas region index.

| | ID | Region | Side | Yeo network |
|---|---|---|---|---|
| 1 | 1 | Precentral gyrus | L | Somatomotor |
| 2 | 15 | Supplementary motor area | L | Somatomotor |
| 3 | 37 | Middle cingulate & paracingulate gyri | L | Ventral Attention |
| 4 | 61 | Postcentral gyrus | L | Somatomotor |
| 5 | 63 | Superior parietal gyrus | L | Dorsal Attention |
| 6 | 65 | Inferior parietal gyrus | L | Dorsal Attention |
| 7 | 71 | Precuneus | L | Default |
| 8 | 73 | Paracentral lobule | L | Somatomotor |

Table 29: Cycle 46: 12 total schizophrenia subjects, 14 total healthy controls. ID is AAL2 atlas region index.

| | ID | Region | Side | Yeo network |
|---|---|---|---|---|
| 1 | 4 | Superior frontal gyrus dorsolateral | R | Default |
| 2 | 16 | Supplementary motor area | R | Somatomotor |
| 3 | 19 | Superior frontal gyrus medial | L | Default |
| 4 | 20 | Superior frontal gyrus medial | R | Default |
| 5 | 35 | Anterior cingulate & paracingulate gyri | L | Default |
| 6 | 36 | Anterior cingulate & paracingulate gyri | R | Default |
| 7 | 38 | Middle cingulate & paracingulate gyri | R | Ventral Attention |

Table 30: Cycle 45: 13 total schizophrenia subjects, 13 total healthy controls. ID is AAL2 atlas region index.

| | ID | Region | Side | Yeo network |
|---|---|---|---|---|
| 1 | 6 | Middle frontal gyrus | R | Frontoparietal |
| 2 | 10 | Inferior frontal gyrus triangular part | R | Frontoparietal |
| 3 | 26 | Medial orbital gyrus | R | Limbic |
| 4 | 28 | Anterior orbital gyrus | R | Frontoparietal |
| 5 | 30 | Posterior orbital gyrus | R | Limbic |
| 6 | 34 | Insula | R | Ventral Attention |

Table 31: Cycle 47: 10 total schizophrenia subjects, 15 total healthy controls. ID is AAL2 atlas region index. A "mirror twin" to Cycle 34 (Table 63).

| | ID | Region | Side | Yeo network |
|---|---|---|---|---|
| 1 | 44 | Parahippocampal gyrus | R | Limbic |
| 2 | 60 | Fusiform gyrus | R | Visual |
| 3 | 88 | Temporal pole superior temporal gyrus | R | Limbic |
| 4 | 92 | Temporal pole middle temporal gyrus | R | Limbic |
| 5 | 94 | Inferior temporal gyrus | R | Limbic |

Table 32: Cycle 49: 15 total schizophrenia subjects, 9 total healthy controls. ID is AAL2 atlas region index.

| | ID | Region | Side | Yeo network |
|---|---|---|---|---|
| 1 | 50 | Cuneus | R | Visual |
| 2 | 54 | Superior occipital gyrus | R | Visual |
| 3 | 64 | Superior parietal gyrus | R | Dorsal Attention |
| 4 | 72 | Precuneus | R | Dorsal Attention |

Table 33: Cycle 50: 9 total schizophrenia subjects, 14 total healthy controls. ID is AAL2 atlas region index.

| | ID | Region | Side | Yeo network |
|---|---|---|---|---|
| 1 | 47 | Calcarine fissure and surrounding cortex | L | Visual |
| 2 | 48 | Calcarine fissure and surrounding cortex | R | Visual |
| 3 | 49 | Cuneus | L | Visual |
| 4 | 50 | Cuneus | R | Visual |

Table 34: Cycle 51: 14 total schizophrenia subjects, 9 total healthy controls. ID is AAL2 atlas region index.

| | ID | Region | Side | Yeo network |
|---|---|---|---|---|
| 1 | 55 | Middle occipital gyrus | L | Visual |
| 2 | 63 | Superior parietal gyrus | L | Dorsal Attention |
| 3 | 65 | Inferior parietal gyrus | L | Dorsal Attention |
| 4 | 69 | Angular gyrus | L | Default |

Table 35: Cycle 52: 11 total schizophrenia subjects, 12 total healthy controls. ID is AAL2 atlas region index.

| | ID | Region | Side | Yeo network |
|---|---|---|---|---|
| 1 | 49 | Cuneus | L | Visual |
| 2 | 53 | Superior occipital gyrus | L | Visual |
| 3 | 55 | Middle occipital gyrus | L | Visual |
| 4 | 63 | Superior parietal gyrus | L | Dorsal Attention |
| 5 | 65 | Inferior parietal gyrus | L | Dorsal Attention |
| 6 | 69 | Angular gyrus | L | Default |
| 7 | 71 | Precuneus | L | Default |

Table 36: Cycle 53: 12 total schizophrenia subjects, 11 total healthy controls. ID is AAL2 atlas region index.

| | ID | Region | Side | Yeo network |
|---|---|---|---|---|
| 1 | 3 | Superior frontal gyrus dorsolateral | L | Default |
| 2 | 19 | Superior frontal gyrus medial | L | Default |
| 3 | 21 | IFG pars orbitalis Med_Orb | L | Default |
| 4 | 35 | Anterior cingulate & paracingulate gyri | L | Default |

Table 37: Cycle 54: 11 total schizophrenia subjects, 11 total healthy controls. ID is AAL2 atlas region index.

| | ID | Region | Side | Yeo network |
|---|---|---|---|---|
| 1 | 1 | Precentral gyrus | L | Somatomotor |
| 2 | 5 | Middle frontal gyrus | L | Frontoparietal |
| 3 | 9 | Inferior frontal gyrus triangular part | L | Frontoparietal |
| 4 | 13 | Rolandic operculum | L | Somatomotor |
| 5 | 33 | Insula | L | Ventral Attention |
| 6 | 49 | Cuneus | L | Visual |
| 7 | 53 | Superior occipital gyrus | L | Visual |
| 8 | 55 | Middle occipital gyrus | L | Visual |
| 9 | 61 | Postcentral gyrus | L | Somatomotor |
| 10 | 63 | Superior parietal gyrus | L | Dorsal Attention |
| 11 | 65 | Inferior parietal gyrus | L | Dorsal Attention |
| 12 | 71 | Precuneus | L | Default |
| 13 | 85 | Superior temporal gyrus | L | Somatomotor |
| 14 | 89 | Middle temporal gyrus | L | Default |

Table 38: Cycle 55: 12 total schizophrenia subjects, 10 total healthy controls. ID is AAL2 atlas region index.

| | ID | Region | Side | Yeo network |
|---|---|---|---|---|
| 1 | 5 | Middle frontal gyrus | L | Frontoparietal |
| 2 | 9 | Inferior frontal gyrus triangular part | L | Frontoparietal |
| 3 | 11 | IFG pars orbitalis Inf_Orb | L | Default |
| 4 | 25 | Medial orbital gyrus | L | Limbic |
| 5 | 27 | Anterior orbital gyrus | L | Limbic |
| 6 | 29 | Posterior orbital gyrus | L | Limbic |

##### B.11.3 Popular, but not significant dimension 1 cycles

Table 39: Cycle 1: 44 total schizophrenia subjects, 44 total healthy controls. ID is AAL2 atlas region index. A "mirror twin" to Cycle 8 (Table 46).

| | ID | Region | Side | Yeo network |
|---|---|---|---|---|
| 1 | 1 | Precentral gyrus | L | Somatomotor |
| 2 | 5 | Middle frontal gyrus | L | Frontoparietal |
| 3 | 7 | Inferior frontal gyrus opercular part | L | Ventral Attention |
| 4 | 9 | Inferior frontal gyrus triangular part | L | Frontoparietal |

Table 40: Cycle 2: 41 total schizophrenia subjects, 42 total healthy controls. ID is AAL2 atlas region index.

| | ID | Region | Side | Yeo network |
|---|---|---|---|---|
| 1 | 39 | Posterior cingulate gyrus | L | Default |
| 2 | 40 | Posterior cingulate gyrus | R | Default |
| 3 | 71 | Precuneus | L | Default |
| 4 | 72 | Precuneus | R | Dorsal Attention |

Table 41: Cycle 3: 38 total schizophrenia subjects, 39 total healthy controls. ID is AAL2 atlas region index.

| | ID | Region | Side | Yeo network |
|---|---|---|---|---|
| 1 | 3 | Superior frontal gyrus dorsolateral | L | Default |
| 2 | 4 | Superior frontal gyrus dorsolateral | R | Default |
| 3 | 15 | Supplementary motor area | L | Somatomotor |
| 4 | 16 | Supplementary motor area | R | Somatomotor |
| 5 | 19 | Superior frontal gyrus medial | L | Default |
| 6 | 20 | Superior frontal gyrus medial | R | Default |

Table 42: Cycle 4: 38 total schizophrenia subjects, 38 total healthy controls. ID is AAL2 atlas region index.

| | ID | Region | Side | Yeo network |
|---|---|---|---|---|
| 1 | 19 | Superior frontal gyrus medial | L | Default |
| 2 | 20 | Superior frontal gyrus medial | R | Default |
| 3 | 35 | Anterior cingulate & paracingulate gyri | L | Default |
| 4 | 36 | Anterior cingulate & paracingulate gyri | R | Default |

Table 43: Cycle 5: 38 total schizophrenia subjects, 38 total healthy controls. ID is AAL2 atlas region index. A "mirror twin" to Cycle 6 (Table 44).

| | ID | Region | Side | Yeo network |
|---|---|---|---|---|
| 1 | 48 | Calcarine fissure and surrounding cortex | R | Visual |
| 2 | 50 | Cuneus | R | Visual |
| 3 | 52 | Lingual gyrus | R | Visual |
| 4 | 54 | Superior occipital gyrus | R | Visual |
| 5 | 56 | Middle occipital gyrus | R | Visual |
| 6 | 60 | Fusiform gyrus | R | Visual |
| 7 | 90 | Middle temporal gyrus | R | Default |
| 8 | 94 | Inferior temporal gyrus | R | Limbic |

Table 44: Cycle 6: 39 total schizophrenia subjects, 35 total healthy controls. ID is AAL2 atlas region index. A "mirror twin" to Cycle 5 (Table 43).

| | ID | Region | Side | Yeo network |
|---|---|---|---|---|
| 1 | 47 | Calcarine fissure and surrounding cortex | L | Visual |
| 2 | 49 | Cuneus | L | Visual |
| 3 | 51 | Lingual gyrus | L | Visual |
| 4 | 53 | Superior occipital gyrus | L | Visual |
| 5 | 55 | Middle occipital gyrus | L | Visual |
| 6 | 59 | Fusiform gyrus | L | Visual |
| 7 | 89 | Middle temporal gyrus | L | Default |
| 8 | 93 | Inferior temporal gyrus | L | Limbic |

Table 45: Cycle 7: 38 total schizophrenia subjects, 36 total healthy controls. ID is AAL2 atlas region index.

| | ID | Region | Side | Yeo network |
|---|---|---|---|---|
| 1 | 62 | Postcentral gyrus | R | Somatomotor |
| 2 | 64 | Superior parietal gyrus | R | Dorsal Attention |
| 3 | 66 | Inferior parietal gyrus | R | Frontoparietal |
| 4 | 68 | Supramarginal gyrus | R | Ventral Attention |

Table 46: Cycle 8: 31 total schizophrenia subjects, 38 total healthy controls. ID is AAL2 atlas region index. A "mirror twin" to Cycle 1 (Table 39).

| | ID | Region | Side | Yeo network |
|---|---|---|---|---|
| 1 | 2 | Precentral gyrus | R | Somatomotor |
| 2 | 6 | Middle frontal gyrus | R | Frontoparietal |
| 3 | 8 | Inferior frontal gyrus opercular part | R | Frontoparietal |
| 4 | 10 | Inferior frontal gyrus triangular part | R | Frontoparietal |

Table 47: Cycle 9: 31 total schizophrenia subjects, 34 total healthy controls. ID is AAL2 atlas region index. A "mirror twin" to Cycle 13 (Table 51).

| | ID | Region | Side | Yeo network |
|---|---|---|---|---|
| 1 | 47 | Calcarine fissure and surrounding cortex | L | Visual |
| 2 | 49 | Cuneus | L | Visual |
| 3 | 51 | Lingual gyrus | L | Visual |
| 4 | 53 | Superior occipital gyrus | L | Visual |
| 5 | 55 | Middle occipital gyrus | L | Visual |
| 6 | 57 | Inferior occipital gyrus | L | Visual |
| 7 | 59 | Fusiform gyrus | L | Visual |

Table 48: Cycle 10: 35 total schizophrenia subjects, 30 total healthy controls. ID is AAL2 atlas region index.

| | ID | Region | Side | Yeo network |
|---|---|---|---|---|
| 1 | 21 | IFG pars orbitalis Med_Orb | L | Default |
| 2 | 22 | IFG pars orbitalis Med_Orb | R | Default |
| 3 | 23 | Gyrus rectus | L | Limbic |
| 4 | 24 | Gyrus rectus | R | Limbic |

Table 49: Cycle 11: 30 total schizophrenia subjects, 30 total healthy controls. ID is AAL2 atlas region index. A "mirror twin" to Cycle 12 (Table 50).

| | ID | Region | Side | Yeo network |
|---|---|---|---|---|
| 1 | 15 | Supplementary motor area | L | Somatomotor |
| 2 | 37 | Middle cingulate & paracingulate gyri | L | Ventral Attention |
| 3 | 71 | Precuneus | L | Default |
| 4 | 73 | Paracentral lobule | L | Somatomotor |

Table 50: Cycle 12: 32 total schizophrenia subjects, 24 total healthy controls. ID is AAL2 atlas region index. A "mirror twin" to Cycle 11 (Table 49).

| | ID | Region | Side | Yeo network |
|---|---|---|---|---|
| 1 | 16 | Supplementary motor area | R | Somatomotor |
| 2 | 38 | Middle cingulate & paracingulate gyri | R | Ventral Attention |
| 3 | 72 | Precuneus | R | Dorsal Attention |
| 4 | 74 | Paracentral lobule | R | Somatomotor |

Table 51: Cycle 13: 27 total schizophrenia subjects, 28 total healthy controls. ID is AAL2 atlas region index. A "mirror twin" to Cycle 9 (Table 47).

| | ID | Region | Side | Yeo network |
|---|---|---|---|---|
| 1 | 48 | Calcarine fissure and surrounding cortex | R | Visual |
| 2 | 50 | Cuneus | R | Visual |
| 3 | 52 | Lingual gyrus | R | Visual |
| 4 | 54 | Superior occipital gyrus | R | Visual |
| 5 | 56 | Middle occipital gyrus | R | Visual |
| 6 | 58 | Inferior occipital gyrus | R | Visual |
| 7 | 60 | Fusiform gyrus | R | Visual |

Table 52: Cycle 14: 26 total schizophrenia subjects, 27 total healthy controls. ID is AAL2 atlas region index. A "mirror twin" to Cycle 27 (Table 59).

| | ID | Region | Side | Yeo network |
|---|---|---|---|---|
| 1 | 3 | Superior frontal gyrus dorsolateral | L | Default |
| 2 | 15 | Supplementary motor area | L | Somatomotor |
| 3 | 19 | Superior frontal gyrus medial | L | Default |
| 4 | 35 | Anterior cingulate & paracingulate gyri | L | Default |
| 5 | 37 | Middle cingulate & paracingulate gyri | L | Ventral Attention |

Table 53: Cycle 15: 28 total schizophrenia subjects, 23 total healthy controls. ID is AAL2 atlas region index.

| | ID | Region | Side | Yeo network |
|---|---|---|---|---|
| 1 | 86 | Superior temporal gyrus | R | Somatomotor |
| 2 | 88 | Temporal pole superior temporal gyrus | R | Limbic |
| 3 | 90 | Middle temporal gyrus | R | Default |
| 4 | 92 | Temporal pole middle temporal gyrus | R | Limbic |

Table 54: Cycle 17: 29 total schizophrenia subjects, 20 total healthy controls. ID is AAL2 atlas region index.

| | ID | Region | Side | Yeo network |
|---|---|---|---|---|
| 1 | 1 | Precentral gyrus | L | Somatomotor |
| 2 | 5 | Middle frontal gyrus | L | Frontoparietal |
| 3 | 9 | Inferior frontal gyrus triangular part | L | Frontoparietal |
| 4 | 13 | Rolandic operculum | L | Somatomotor |
| 5 | 33 | Insula | L | Ventral Attention |
| 6 | 61 | Postcentral gyrus | L | Somatomotor |

Table 55: Cycle 19: 22 total schizophrenia subjects, 24 total healthy controls. ID is AAL2 atlas region index.

| | ID | Region | Side | Yeo network |
|---|---|---|---|---|
| 1 | 14 | Rolandic operculum | R | Somatomotor |
| 2 | 30 | Posterior orbital gyrus | R | Limbic |
| 3 | 34 | Insula | R | Ventral Attention |
| 4 | 86 | Superior temporal gyrus | R | Somatomotor |
| 5 | 88 | Temporal pole superior temporal gyrus | R | Limbic |

Table 56: Cycle 21: 23 total schizophrenia subjects, 21 total healthy controls. ID is AAL2 atlas region index.

| | ID | Region | Side | Yeo network |
|---|---|---|---|---|
| 1 | 1 | Precentral gyrus | L | Somatomotor |
| 2 | 2 | Precentral gyrus | R | Somatomotor |
| 3 | 3 | Superior frontal gyrus dorsolateral | L | Default |
| 4 | 4 | Superior frontal gyrus dorsolateral | R | Default |
| 5 | 5 | Middle frontal gyrus | L | Frontoparietal |
| 6 | 6 | Middle frontal gyrus | R | Frontoparietal |
| 7 | 19 | Superior frontal gyrus medial | L | Default |
| 8 | 20 | Superior frontal gyrus medial | R | Default |
| 9 | 61 | Postcentral gyrus | L | Somatomotor |
| 10 | 62 | Postcentral gyrus | R | Somatomotor |
| 11 | 63 | Superior parietal gyrus | L | Dorsal Attention |
| 12 | 64 | Superior parietal gyrus | R | Dorsal Attention |
| 13 | 65 | Inferior parietal gyrus | L | Dorsal Attention |
| 14 | 71 | Precuneus | L | Default |
| 15 | 72 | Precuneus | R | Dorsal Attention |

Table 57: Cycle 22: 24 total schizophrenia subjects, 19 total healthy controls. ID is AAL2 atlas region index.

| | ID | Region | Side | Yeo network |
|---|---|---|---|---|
| 1 | 50 | Cuneus | R | Visual |
| 2 | 54 | Superior occipital gyrus | R | Visual |
| 3 | 56 | Middle occipital gyrus | R | Visual |
| 4 | 62 | Postcentral gyrus | R | Somatomotor |
| 5 | 64 | Superior parietal gyrus | R | Dorsal Attention |
| 6 | 68 | Supramarginal gyrus | R | Ventral Attention |
| 7 | 72 | Precuneus | R | Dorsal Attention |
| 8 | 86 | Superior temporal gyrus | R | Somatomotor |
| 9 | 90 | Middle temporal gyrus | R | Default |

Table 58: Cycle 25: 23 total schizophrenia subjects, 18 total healthy controls. ID is AAL2 atlas region index.

| | ID | Region | Side | Yeo network |
|---|---|---|---|---|
| 1 | 3 | Superior frontal gyrus dorsolateral | L | Default |
| 2 | 4 | Superior frontal gyrus dorsolateral | R | Default |
| 3 | 15 | Supplementary motor area | L | Somatomotor |
| 4 | 16 | Supplementary motor area | R | Somatomotor |
| 5 | 19 | Superior frontal gyrus medial | L | Default |
| 6 | 20 | Superior frontal gyrus medial | R | Default |
| 7 | 37 | Middle cingulate & paracingulate gyri | L | Ventral Attention |
| 8 | 38 | Middle cingulate & paracingulate gyri | R | Ventral Attention |

Table 59: Cycle 27: 21 total schizophrenia subjects, 18 total healthy controls. ID is AAL2 atlas region index. A "mirror twin" to Cycle 14 (Table 52).

| | ID | Region | Side | Yeo network |
|---|---|---|---|---|
| 1 | 4 | Superior frontal gyrus dorsolateral | R | Default |
| 2 | 16 | Supplementary motor area | R | Somatomotor |
| 3 | 20 | Superior frontal gyrus medial | R | Default |
| 4 | 36 | Anterior cingulate & paracingulate gyri | R | Default |
| 5 | 38 | Middle cingulate & paracingulate gyri | R | Ventral Attention |

Table 60: Cycle 28: 16 total schizophrenia subjects, 21 total healthy controls. ID is AAL2 atlas region index.

| | ID | Region | Side | Yeo network |
|---|---|---|---|---|
| 1 | 15 | Supplementary motor area | L | Somatomotor |
| 2 | 16 | Supplementary motor area | R | Somatomotor |
| 3 | 37 | Middle cingulate & paracingulate gyri | L | Ventral Attention |
| 4 | 38 | Middle cingulate & paracingulate gyri | R | Ventral Attention |

Table 61: Cycle 29: 21 total schizophrenia subjects, 16 total healthy controls. ID is AAL2 atlas region index.

| | ID | Region | Side | Yeo network |
|---|---|---|---|---|
| 1 | 56 | Middle occipital gyrus | R | Visual |
| 2 | 66 | Inferior parietal gyrus | R | Frontoparietal |
| 3 | 68 | Supramarginal gyrus | R | Ventral Attention |
| 4 | 70 | Angular gyrus | R | Default |
| 5 | 86 | Superior temporal gyrus | R | Somatomotor |
| 6 | 90 | Middle temporal gyrus | R | Default |

Table 62: Cycle 31: 16 total schizophrenia subjects, 18 total healthy controls. ID is AAL2 atlas region index.

| | ID | Region | Side | Yeo network |
|---|---|---|---|---|
| 1 | 50 | Cuneus | R | Visual |
| 2 | 54 | Superior occipital gyrus | R | Visual |
| 3 | 56 | Middle occipital gyrus | R | Visual |
| 4 | 62 | Postcentral gyrus | R | Somatomotor |
| 5 | 64 | Superior parietal gyrus | R | Dorsal Attention |
| 6 | 66 | Inferior parietal gyrus | R | Frontoparietal |
| 7 | 68 | Supramarginal gyrus | R | Ventral Attention |
| 8 | 70 | Angular gyrus | R | Default |
| 9 | 72 | Precuneus | R | Dorsal Attention |

Table 63: Cycle 34: 19 total schizophrenia subjects, 14 total healthy controls. ID is AAL2 atlas region index. A "mirror twin" to Cycle 47 (Table 31).

| | ID | Region | Side | Yeo network |
|---|---|---|---|---|
| 1 | 43 | Parahippocampal gyrus | L | Limbic |
| 2 | 59 | Fusiform gyrus | L | Visual |
| 3 | 87 | Temporal pole superior temporal gyrus | L | Limbic |
| 4 | 91 | Temporal pole middle temporal gyrus | L | Limbic |
| 5 | 93 | Inferior temporal gyrus | L | Limbic |

Table 64: Cycle 36: 13 total schizophrenia subjects, 19 total healthy controls. ID is AAL2 atlas region index.

| | ID | Region | Side | Yeo network |
|---|---|---|---|---|
| 1 | 13 | Rolandic operculum | L | Somatomotor |
| 2 | 61 | Postcentral gyrus | L | Somatomotor |
| 3 | 65 | Inferior parietal gyrus | L | Dorsal Attention |
| 4 | 67 | Supramarginal gyrus | L | Ventral Attention |
| 5 | 85 | Superior temporal gyrus | L | Somatomotor |

Table 65: Cycle 39: 14 total schizophrenia subjects, 16 total healthy controls. ID is AAL2 atlas region index.

| | ID | Region | Side | Yeo network |
|---|---|---|---|---|
| 1 | 1 | Precentral gyrus | L | Somatomotor |
| 2 | 5 | Middle frontal gyrus | L | Frontoparietal |
| 3 | 9 | Inferior frontal gyrus triangular part | L | Frontoparietal |
| 4 | 13 | Rolandic operculum | L | Somatomotor |
| 5 | 33 | Insula | L | Ventral Attention |
| 6 | 61 | Postcentral gyrus | L | Somatomotor |
| 7 | 65 | Inferior parietal gyrus | L | Dorsal Attention |
| 8 | 67 | Supramarginal gyrus | L | Ventral Attention |
| 9 | 85 | Superior temporal gyrus | L | Somatomotor |

Table 66: Cycle 38: 13 total schizophrenia subjects, 17 total healthy controls. ID is AAL2 atlas region index.

| | ID | Region | Side | Yeo network |
|---|---|---|---|---|
| 1 | 86 | Superior temporal gyrus | R | Somatomotor |
| 2 | 88 | Temporal pole superior temporal gyrus | R | Limbic |
| 3 | 90 | Middle temporal gyrus | R | Default |
| 4 | 92 | Temporal pole middle temporal gyrus | R | Limbic |
| 5 | 94 | Inferior temporal gyrus | R | Limbic |

Table 67: Cycle 41: 12 total schizophrenia subjects, 17 total healthy controls. ID is AAL2 atlas region index.

| | ID | Region | Side | Yeo network |
|---|---|---|---|---|
| 1 | 2 | Precentral gyrus | R | Somatomotor |
| 2 | 4 | Superior frontal gyrus dorsolateral | R | Default |
| 3 | 6 | Middle frontal gyrus | R | Frontoparietal |
| 4 | 16 | Supplementary motor area | R | Somatomotor |
| 5 | 38 | Middle cingulate & paracingulate gyri | R | Ventral Attention |
| 6 | 62 | Postcentral gyrus | R | Somatomotor |
| 7 | 64 | Superior parietal gyrus | R | Dorsal Attention |
| 8 | 72 | Precuneus | R | Dorsal Attention |

Table 68: Cycle 42: 14 total schizophrenia subjects, 13 total healthy controls. ID is AAL2 atlas region index. A "mirror twin" to Cycle 26 (Table 22).

| | ID | Region | Side | Yeo network |
|---|---|---|---|---|
| 1 | 43 | Parahippocampal gyrus | L | Limbic |
| 2 | 59 | Fusiform gyrus | L | Visual |
| 3 | 87 | Temporal pole superior temporal gyrus | L | Limbic |
| 4 | 89 | Middle temporal gyrus | L | Default |
| 5 | 91 | Temporal pole middle temporal gyrus | L | Limbic |
| 6 | 93 | Inferior temporal gyrus | L | Limbic |

Table 69: Cycle 44: 9 total schizophrenia subjects, 17 total healthy controls. ID is AAL2 atlas region index.

| | ID | Region | Side | Yeo network |
|---|---|---|---|---|
| 1 | 19 | Superior frontal gyrus medial | L | Default |
| 2 | 20 | Superior frontal gyrus medial | R | Default |
| 3 | 21 | IFG pars orbitalis Med_Orb | L | Default |
| 4 | 22 | IFG pars orbitalis Med_Orb | R | Default |
| 5 | 35 | Anterior cingulate & paracingulate gyri | L | Default |

Table 70: Cycle 43: 12 total schizophrenia subjects, 14 total healthy controls. ID is AAL2 atlas region index.

| | ID | Region | Side | Yeo network |
|---|---|---|---|---|
| 1 | 17 | Olfactory cortex | L | Limbic |
| 2 | 21 | IFG pars orbitalis Med_Orb | L | Default |
| 3 | 23 | Gyrus rectus | L | Limbic |
| 4 | 35 | Anterior cingulate & paracingulate gyri | L | Default |

Table 71: Cycle 48: 13 total schizophrenia subjects, 11 total healthy controls. ID is AAL2 atlas region index. A "mirror twin" to Cycle 37 (Table 27).

| | ID | Region | Side | Yeo network |
|---|---|---|---|---|
| 1 | 55 | Middle occipital gyrus | L | Visual |
| 2 | 57 | Inferior occipital gyrus | L | Visual |
| 3 | 59 | Fusiform gyrus | L | Visual |
| 4 | 89 | Middle temporal gyrus | L | Default |
| 5 | 93 | Inferior temporal gyrus | L | Limbic |

#### B.12 Brain regions in AAL2 atlas

Table 72: A numbered list of brain regions from the *AAL2* [Rolls et al., 2015] atlas and the corresponding large-scale Yeo [Yeo et al., 2011] network to which the brain area belongs.

| *N* | *Brain region* | *Yeo network* | *N* | *Brain region* | *Yeo network* |
|---|---|---|---|---|---|
| 1 | Precentral L | Somatomotor | 48 | Calcarine R | Visual |
| 2 | Precentral R | Somatomotor | 49 | Cuneus L | Visual |
| 3 | Frontal Sup 2 L | Default | 50 | Cuneus R | Visual |
| 4 | Frontal Sup 2 R | Default | 51 | Lingual L | Visual |
| 5 | Frontal Mid 2 L | Frontoparietal | 52 | Lingual R | Visual |
| 6 | Frontal Mid 2 R | Frontoparietal | 53 | Occipital Sup L | Visual |
| 7 | Frontal Inf Oper L | Ventral Attention | 54 | Occipital Sup R | Visual |
| 8 | Frontal Inf Oper R | Frontoparietal | 55 | Occipital Mid L | Visual |
| 9 | Frontal Inf Tri L | Frontoparietal | 56 | Occipital Mid R | Visual |
| 10 | Frontal Inf Tri R | Frontoparietal | 57 | Occipital Inf L | Visual |
| 11 | Frontal Inf Orb 2 L | Default | 58 | Occipital Inf R | Visual |
| 12 | Frontal Inf Orb 2 R | Default | 59 | Fusiform L | Visual |
| 13 | Rolandic Oper L | Somatomotor | 60 | Fusiform R | Visual |
| 14 | Rolandic Oper R | Somatomotor | 61 | Postcentral L | Somatomotor |
| 15 | Supp Motor Area L | Somatomotor | 62 | Postcentral R | Somatomotor |
| 16 | Supp Motor Area R | Somatomotor | 63 | Parietal Sup L | Dorsal Attention |
| 17 | Olfactory L | Limbic | 64 | Parietal Sup R | Dorsal Attention |
| 18 | Olfactory R | Limbic | 65 | Parietal Inf L | Dorsal Attention |
| 19 | Frontal Sup Medial L | Default | 66 | Parietal Inf R | Frontoparietal |
| 20 | Frontal Sup Medial R | Default | 67 | SupraMarginal L | Ventral Attention |
| 21 | Frontal Med Orb L | Default | 68 | SupraMarginal R | Ventral Attention |
| 22 | Frontal Med Orb R | Default | 69 | Angular L | Default |
| 23 | Rectus L | Limbic | 70 | Angular R | Default |
| 24 | Rectus R | Limbic | 71 | Precuneus L | Default |
| 25 | OFCmed L | Limbic | 72 | Precuneus R | Dorsal Attention |
| 26 | OFCmed R | Limbic | 73 | Paracentral Lobule L | Somatomotor |
| 27 | OFCant L | Limbic | 74 | Paracentral Lobule R | Somatomotor |
| 28 | OFCant R | Frontoparietal | 75 | Caudate L | Limbic |
| 29 | OFCpost L | Limbic | 76 | Caudate R | - |
| 30 | OFCpost R | Limbic | 77 | Putamen L | - |
| 31 | OFClat L | Default | 78 | Putamen R | Ventral Attention |
| 32 | OFClat R | Default | 79 | Pallidum L | - |
| 33 | Insula L | Ventral Attention | 80 | Pallidum R | - |
| 34 | Insula R | Ventral Attention | 81 | Thalamus L | - |
| 35 | Cingulate Ant L | Default | 82 | Thalamus R | - |
| 36 | Cingulate Ant R | Default | 83 | Heschl L | Somatomotor |
| 37 | Cingulate Mid L | Ventral Attention | 84 | Heschl R | Somatomotor |
| 38 | Cingulate Mid R | Ventral Attention | 85 | Temporal Sup L | Somatomotor |
| 39 | Cingulate Post L | Default | 86 | Temporal Sup R | Somatomotor |
| 40 | Cingulate Post R | Default | 87 | Temporal Pole Sup L | Limbic |
| 41 | Hippocampus L | - | 88 | Temporal Pole Sup R | Limbic |
| 42 | Hippocampus R | Limbic | 89 | Temporal Mid L | Default |
| 43 | ParaHippocampal L | Limbic | 90 | Temporal Mid R | Default |
| 44 | ParaHippocampal R | Limbic | 91 | Temporal Pole Mid L | Limbic |
| 45 | Amygdala L | Limbic | 92 | Temporal Pole Mid R | Limbic |
| 46 | Amygdala R | - | 93 | Temporal Inf L | Limbic |
| 47 | Calcarine L | Visual | 94 | Temporal Inf R | Limbic |

The AAL2 atlas is a volumetric atlas and a successor of the AAL atlas [Tzourio-Mazoyer et al., 2002], which is popular in schizophrenia structural studies [Gao et al., 2023], the difference being the introduction of the parcellation of the orbitofrontal cortex. Both divide the brain into Regions of Interest (ROI) but omit subcortical structures [Revell et al., 2022]. The exclusion of subcortical structures in the AAL2 atlas is a potential limitation, as their inclusion might provide further insight into disease mechanisms. For instance, the AAL3 atlas is an updated version of the AAL2 atlas, which adds several brain areas, for example, it subdivides the cingulate cortex into more regions [Rolls et al., 2020]. A more fine-grained division of the cingulate cortex might be especially insightful as it is reported to be strongly affected by schizophrenia (reduction of grey matter volume [Brugger and Howes, 2017, Fortea et al., 2021, Merritt et al., 2021], reduction of grey matter density [Glahn et al., 2008]). Another option could be the Schaefer atlas [Schaefer et al., 2018], covering from 100 up to 400 cortical areas- more regions would provide more details about the connections. However, topological computations using parcellation with 200 or more brain regions might be computationally infeasible for higher dimensions.

#### B.13 Strengths, limitations and future directions

The choice of scanners could have an impact on the results if, for example, the differences in Fractional Anisotropy (FA) recorded could result in changes in the ranking of connections obtained. Therefore, our method may be affected by the choice of preprocessing, but to a lesser extent than the choice of scanner. Since we are able to reproduce results across datasets, this shows high robustness to the choice of scanner.

We used Probabilistic Fibre Tracking (PFT) (to obtain the connectivity matrix), which has difficulty tracking long connections [Ambrosen et al., 2020, Reveley et al., 2015], such as cross-hemisphere or sub-cortical connections, and changes to the parameters used in the algorithm can significantly affect the results [Gutierrez et al., 2020]. In particular, it can return false positives [Thomas et al., 2014, Drakesmith et al., 2015, Maier-Hein et al., 2017], that is incorrectly identify certain fibre tracts as real. Our method mitigates this by using the order complex, thereby focusing on the relative connectivity within a subject, rather than scanner and processing-dependent absolute values.

It is the weaker connections that are more likely to be affected by data acquisition and preprocessing variability. In support of this, we demonstrated that shuffling the weak connections in submatrices for *HCP* (matrix size limited to the first 30 *AAL2* brain regions, and dimensions up to 8), the resulting topological properties are not changed for perturbations of a large portion of the connections - there are no changes in cycles for dimensions $D = 0, 1, 2$ for up to 50% of the weights, (for more details, see Supplementary Results B.9.1). Moreover, in *COBRE*, many zero-connections were present because the PFT algorithm was unable to reconstruct weak connections (this can be seen in the sample connectivity matrix in Figure 7). These absent connections in *COBRE* correspond to weak connections in *HCP* (collected with a more modern acquisition system that could detect weaker connections).

Another limitation may come from the choice of the atlas used for brain parcellation. See Section B.12 for a detailed discussion of atlas choice.

Our study was conducted on a diverse population of schizophrenia subjects (see Section B.10). A potential future direction is to focus on topological connectomes of subpopulations (e.g. early onset, no medication patients), or to link our findings with the psychiatric evaluation of the disorder, e.g. establish the possible link with the positive or negative symptoms from the PANAS scale.

White matter connectomic fingerprinting of subjects with treatment-resistant (TR) schizophrenia [Luvsannyam et al., 2022] could direct future neurological investigations of white matter changes and could be further correlated with genetic markers. TR patients, when compared to both healthy controls and treatment responding (TRS) schizophrenia patients, show decreases in white matter volume in frontal, occipital and parietal regions [Molina et al., 2008].

During childhood and adolescence, brain structure alterations progress, and schizophrenia patients' brains experience excessive loss of neurons and synaptic connections [Chung and Cannon, 2015, Haukvik et al., 2013], this process could be tracked with connectomic fingerprinting in longitudinal studies.

Studies [Shen et al., 2023, Cropley et al., 2017] have demonstrated that the progression of white matter changes in schizophrenia patients occurs gradually over time. The changes occur in limited areas at early stages (disease duration below 5 years), however, in later stages (above 25 years of disease), almost all white matter tracts show FA reduction.

While those studies report which of the white matter tracts are are affected, they do not analyse the brain regions whose connectivity is affected with the progression of the disease, for example, they report an average reduction in fractional anisotropy in *inferior fronto-occipital fasciculus*, but do not specify which of the regions it connects are affected at what level. By incorporating our approach in longitudinal studies, it would be possible to learn how the progression of the disease affects parts of the brain tracts.

Our method is a way to create a detailed connectivity profile of the human brain. By extending this approach to a larger population of healthy subjects, we would be able to establish a DTI-based topological connectomic fingerprint. Furthermore, by obtaining fingerprints from patients suffering from various neurological and psychiatric disorders, it would be possible to ascertain the cross-disorder and disorder-specific changes in cycles, fulfilling the idea of an individual profile, and individual disease prognostics. Cross-disorder connectotyping could create detailed maps of a person's connectivity profile, potentially helping to determine their risk or resilience to certain brain disorders.
